# Analysis of nucleotide variations in human g-quadruplex forming regions associated with disease states

**DOI:** 10.1101/2023.01.30.526341

**Authors:** Aryan Neupane, Julia H. Chariker, Eric C. Rouchka

## Abstract

While the role of G4 G quadruplex structures has been identified in cancers and metabolic disorders, single nucleotide variations (SNVs) and their effect on G4s in disease contexts have not been extensively studied. The COSMIC and CLINVAR databases were used to detect SNVs present in G4s to identify sequence level changes and their effect on alteration of G4 secondary structure. 37,515 G4 SNVs in the COSMIC database and 2,115 in CLINVAR were identified. Of those, 7,236 COSMIC (19.3%) and 416 (18%) of the CLINVAR variants result in G4 loss, while 2,728 (COSMIC) and 112 (CLINVAR) SNVs gain a G4 structure. The gene ontology term “GnRH (Gonadotropin-releasing hormone) secretion” is enriched in 21 genes in this pathway that have a G4 destabilizing SNV. Analysis of mutational patterns in the G4 structure show a higher selective pressure (3-fold) in the coding region on the template strand compared to the non-template strand. At the same time, an equal proportion of SNVs were observed among intronic, promoter and enhancer regions across strands. Using GO and pathway enrichment, genes with SNVs for G4 forming propensity in the coding region are enriched for Regulation of Ras protein signal transduction and Src homology 3 (SH3) domain binding.

## INTRODUCTION

G-quadruplexes are stranded secondary structures of nucleic acids rich in guanine. These nucleic acid sequences are characterized by four runs of at least three guanines separated by short loops, which can potentially fold into an intramolecular or intermolecular G-quadruplex structure (1). The tetrad structure of guanine is stacked on top of each other and held together by mixed loops of DNA giving a four-stranded structure that has nucleobases on the inside forming Hoogsteen base pairing and the sugar phosphate backbone on the outside (Figure 1). They are found in G-rich sequences of both DNA and RNA and are stabilized by metal cations such as potassium (K+) or sodium (Na+) (2). The binding energy is held through the H bonding between the guanines, stabilized by π-π interactions and charge interactions between the sixth position of oxygen (O6) and cations (K+, Na+) between the stacks. The structural architecture of a G-quadruplex is quite diverse and can form different topologies based on factors such as the chemical environment, loop length (3,4), and localization in the sequence or structure molecularity (5). The stacking of the guanine tetrads is bound by the loops of nucleotide bases of variable sizes which determine the folding of the secondary structure.

**Figure 1.**
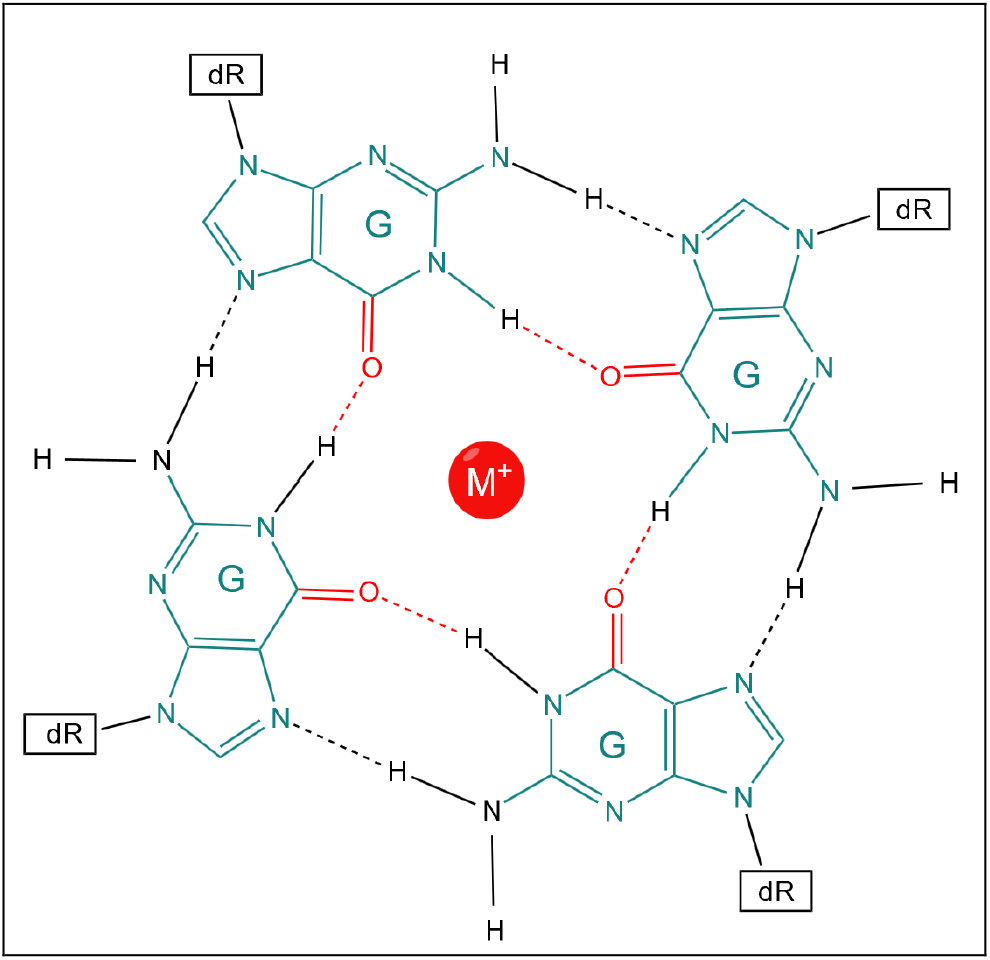
Guanine tetrad formed by Hoogsteen bond formation.

### Functional role of G4 regions

G4 sequences do not always form G4 structures, which can additionally be dependent upon physiological conditions and methylation patterns guided by chromatin structure for their formation (6). However, when they do, they can alter several functional roles. One such perturbed function is transcription which is affected by stalling the replication fork (7-9). In cells that do not have the normal DNA repair machinery, this causes down regulation of several genes and cell cycle arrest (10).

Additionally, G4, G4 stabilizing agents and double-stranded breaks (DSB) facilitate the homologous recombination repair pathway affecting genome instability. Based on the size of the G quadruplex, thermodynamically stable short loop structures within the G4 have been extensively studied to cause instability in replication dependent processes (11). Alteration of DNA polymerase function and helicases in sites of G4 formation has been well established and is used in identification of G quadruplexes *in vivo* (12,13).

While some ligands have shown binding affinity towards G quadruplex structures for treatment of cancer specific cells and transcriptional alteration (14), binding of other ligands that stabilize G4 lead to multiple DNA damage (14), micronuclei formation, delayed replication fork progression (15), and telomeric defects (16-18).

### Mutations within G4 regions

DNA lesions can be mutagenic or lethal, and when they are found in G quadruplex regions, they can alter the secondary structure by changing the guanine tract base pairing or altering the composition of the loop region. A single nucleotide mutation in the G4 present in the promoter region of c-MYC has been shown to change transcription *in vivo* (19). Mass spectroscopy studies using single nucleotide substitution in the central block of parallel G4 forming sequencing found a deleterious effect of G quadruplex stability and association rate (20). A trinucleotide CGG repeat expansion in the untranslated region of the *FMR1* gene has been linked with ataxias and Fragile X Syndrome (21). A T→C SNP at the GC rich region of Apolipoprotein E (APOE) is known to vary G quadruplex structure and has been linked to onset of Alzheimer’s Disease (22). It has been proposed that specific helicases promote genomic stability by actively resolving G4 structures which can be altered by the addition of G4 stabilization ligands presence of specific DSBs (10,12,23). Baral *et al*. identified several eQTL variants in potential G-quadruplex regions (24). Changes in loops of G quadruplexes and stability led to a significant alteration in gene expression among individuals further fueling the structural role of G4s in regulation and binding of transcription factors (23).

Selective mutation of the G rich region to disrupt the G4 structure has been found to alter transcription. The mutation further can hinder the recruitment of transcription factors that overlap the G rich region and function as recognition motifs or bind to the G quadruplex region. Siddiqui-Jain et al. demonstrated that a single G→A mutation destabilizes the folding of G4 in the Pu27 region of MYC which is otherwise repressed, resulting in a threefold increase in transcriptional activity of the gene in tumor cell lines (19). Studies related to 8-oxoguanine in the G quadruplex established the presence of G-A and guanine abasic lesions in G quadruplex structures based on the position in the sequence which can destabilize the secondary structure leaving the unfolded sequence prone to cleavage, leading to further instability in the telomere region (25).

### Study motivation

Given the roles that G4 regions and mutations within them play in transcriptional and translational control, we set out to identify the impacts of mutations in G-quadruplex regions and patterns associated with the variants. This was aided by looking at variants annotated in the COSMIC (26) and CLINVAR (27) databases, which represent mutations associated with cancers (COSMIC) or other clinical relevance (CLINVAR). We identified somatic and germline variations representing SNVs occurring within G quadruplex sequences. Because of their high stability and increased cellular uptake, G quadruplex sequences have interesting diagnostic and therapeutic functions. Understanding how known variants in the genome confer stability or disrupt the G quadruplex sequences will allow a better understanding of G4 structure and function.

## MATERIAL AND METHODS

### Putative and validated G4 identification

Quadparser version 2 (28) with the default parameters was used to identify 175,778 putative G quadruplex regions in the human genome hg38 assembly across both strands. Experimentally validated G4 regions were obtained from an experiment utilizing a method called G4 Seq (GEO accession GSE63874) previously performed by Chambers, et al. (29). The intersection between the putative and experimental G quadruplexes was found using BEDTOOLS (30).

### SNP identification

Cancer-specific curated somatic mutations from the COSMIC database (26) were used for the analysis. COSMIC contains 22,996,215 distinct single nucleotide variants (SNVs) (19,721,019 non-coding variants (NCV) and 5,977,977 coding) from 1.4 million tumor samples. An additional 550,239 germline SNVs from other clinically relevant diseases and disorders were obtained from CLINVAR (27) version (clinvar_20200203.vcf.gz).

For both sets of data, a two-pass analysis was performed. In the first pass, overlaps between the SNVs and putative G quadruplex regions were found to determine potential loss of a G quadruplex structure due to mutations. In the second pass, mutations leading to a G in regions with flanking guanines that could result in the gain of a G quadruplex were detected. In each case, a variant call format (VCF) file describing the coding and non-coding mutations was obtained from COSMIC (26) and CLINVAR (27). Using the VCF, SNVs were filtered using bcftools (31), with insertion and deletion events (INDELs) removed.

### Identification of SNPs affecting G4 formation

A window 30 bases upstream and 30 bases downstream of each variant was used to search for putative G quadruplex sequences. Prospective G4 regions were compared with the Vienna Package RNAfold v2.4.8 to determine changes in G quadruplex stability as a result of the variant (32). The values of ΔMFE (minimum free energy) and ΔED (ensemble diversity) were used as the determining metrics. MFE calculates the stability of the sequence structure based on the binding propensities while centroid distance to ensemble provides the diversity of the sequence structure and alternate structures it can form. G4hunter was also used to compare the G4 scores and the formation of pG4 (33).

Based on the location of a specific SNV inside a G4 region, the relative location of the mutation was calculated as the position of the SNV in the G4 divided by the total length of the sequence. In terms of multiple potential G4 regions, the whole region was used as a single sequence and the relative location of the mutation was calculated. Each SNV was converted into a 3-mer based on its context and changes in the k-mer resulting in a broken GGG quad structure were calculated. For each 3-mer, the number of changes was calculated using one base before and after the location of the variant, respectively. In addition, the SNV in the context of loop and guanine tetrads was analyzed based on the trinucleotide context. The R package annotatr was used for randomized background counts for each annotation (34).

### Enrichment analysis

Based on the G4 identified, the hg38 coordinates of the G4 were used to find the enrichment of transcriptional factors using Remap (35) for Hep-G2, K562, HEK293 and HEK293T cell lines. Further, enrichment analysis of the genes with individual mutations were selected based on the number of SNV per gene, effect of SNVs on the G quadruplex, G4 per gene and samples as specified in the result. Functional annotation enrichment of genes was carried out using DAVID functional annotation (36) while the enrichment analysis of TFs involved was carried out using STRING database (37). In order to analyze the disruption of motifs by each SNV, the R package motifbreakR (38) was used.

## RESULTS

### COSMIC somatic mutations

Using the COSMIC database, 37,515 (0.16% of all COSMIC mutations) distinct single nucleotide somatic mutations were identified within 26,504 pG4 regions from 9,693 genes, 8,998 of which were determined to be protein coding according to ENSEMBL hg38 annotations. The remaining genes were identified as lncRNA (n=540) or miRNA (n=111). The most frequently observed mutation observed in the COSMIC filtered dataset was the transition event G→ A (28%) followed by the transversion event T→G (18%) (Figures 2A and 2B). The variants were expected to be high in number for G→A and G→T (15%) mutations; however, we also identified the T→G transversion to be high in these regions compared with A→G transitions. Comparatively, higher G/C→A/T variants in intragenic CpG islands has been observed due to the spontaneous deamination to the cytosine hypermethylated CpGs within these regions (39,40). However, the effect of these mutations is less studied across G4 regions. We found a lower transition:transversion ratio (p = 0.00001) occurring in the G4 region (1.02), compared to the overall mutations in COSMIC database (1.146) (Supplemental Tables 1 and 2).

**Figure 2.**
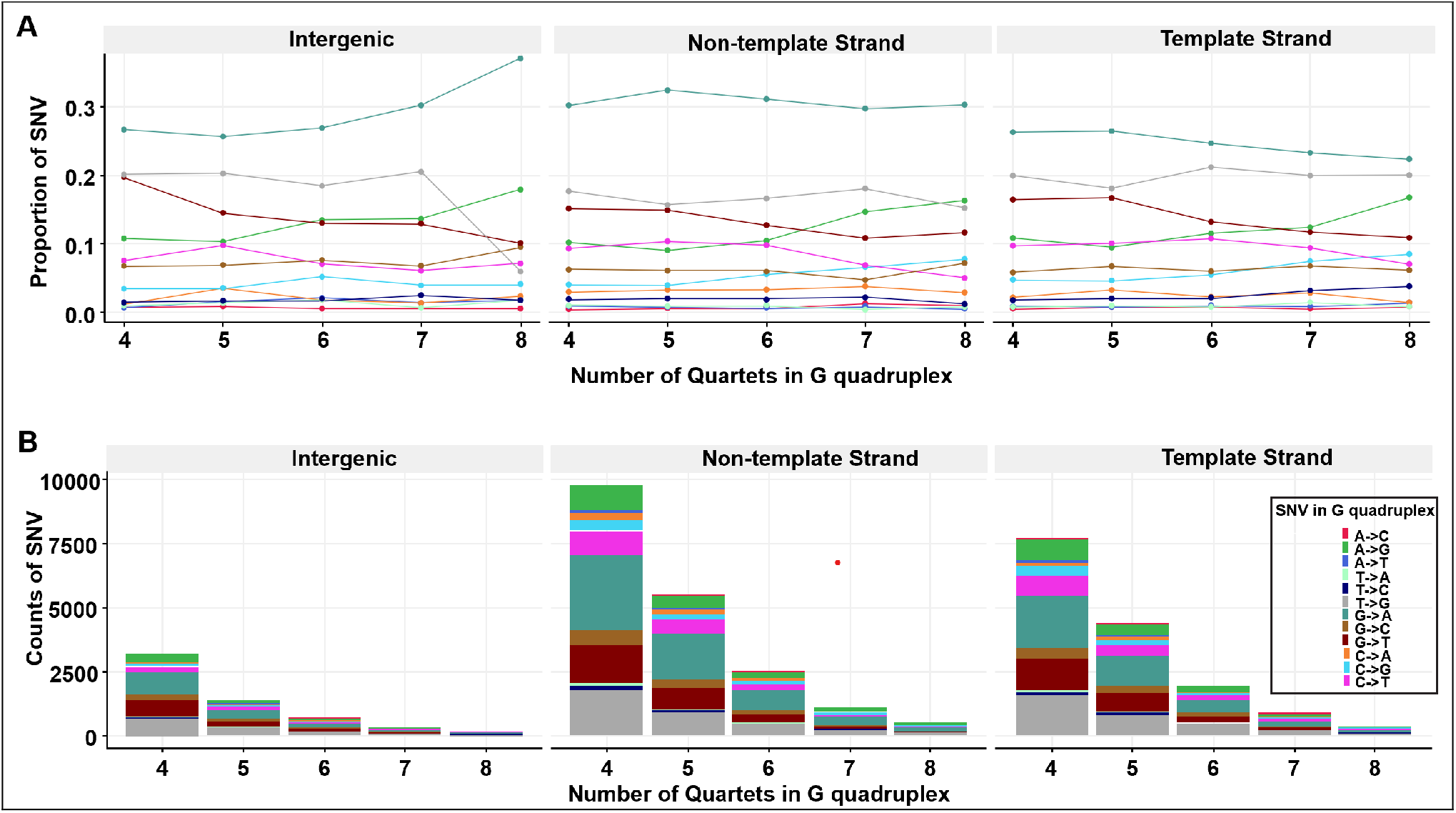
Composition of SNVs in G4 regions from the COSMIC database. Shown is (A) proportion and (B) count of selected SNVs.

Based on the G4Hunter (33) and RNAfold (32) results, we compared the number of SNV events that breaks the G4 structure and changes in the thermodynamic stability based on the minimum fold energy of each sequence. We found 7,236 (19.2% of variants in G4) of the SNVs within the G4Hunter identified G4s result in the loss of a G quadruplex, while 2,728 SNVs led to the gain of a new G quadruplex (Figure 3A, Supplemental Table 3).

**Figure 3.**
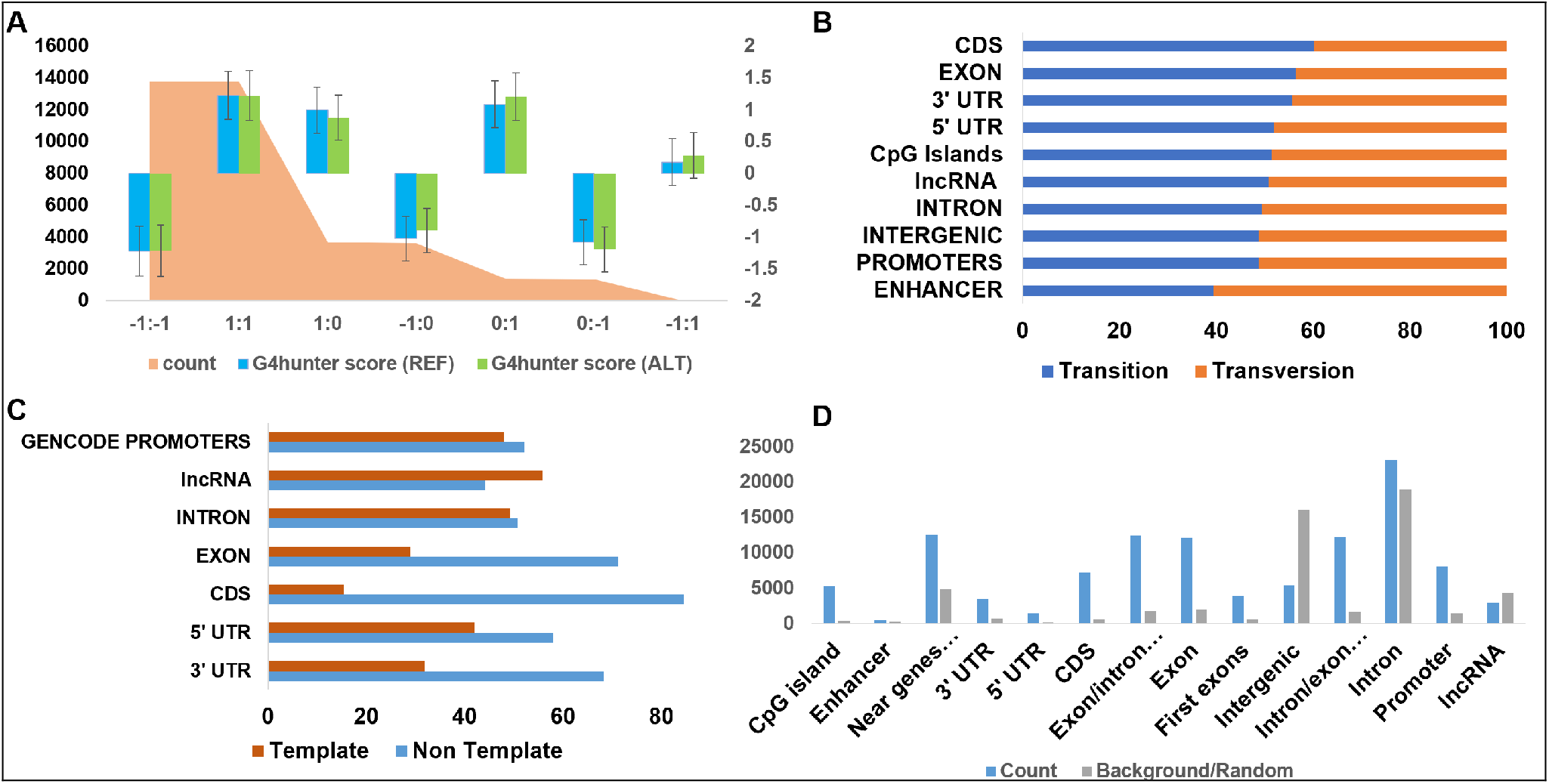
Identified G4 variants relative to functional annotations. Shown is (A) count of change in pG4 with G4Hunter score across both strands before and after mutation (0: absence of pG4; 1: presence of G4 in the forward strand; -1: presence of G4 in the reverse strand); (B) percentage of the type of mutation across annotations from the COSMIC database; (C) percentage of SNVs that occur in a G4 region across the template and non-template strand for functional annotation groups; and (D) count of variants in functional annotations against randomized background count of variants in the human genome.

### CLINVAR germline mutations

Using the CLINVAR database, 5,026 SNVs were identified in pG4 regions out of which 2,155 intersected with experimental G4. Most of these G4 mutations occurring in exons (50%, n=2,559). The remaining variants are found in introns (24%, n=1,251), promoters (11%, n=554), and transcription termination regions (3.5%, n=179). Overall, 13.92% (700 variants) were associated with non-coding RNA, and 84% (n=4,265) SNVs occur in protein coding regions (Figures 3B-3D, Supplemental Table 4).

### Change to G4 stability

RNAfold was used to differentiate the impact of the variant on the stacking. Variants were classified based on the change in stability and formation of available guanines for stacking by combining the sequence pattern analysis of G4Hunter with thermodynamic parameters from RNAfold (Figures 4A-4F). The majority of the SNVs (81%) did not affect the GGG stacking in such a way that the formation of tetrads of guanines was not possible. Though complete breakage of structure does not occur, we found a decrease in the stability of the G quadruplex structure in 40% of these variants. This is due to the presence of additional guanines in the loop that aid the conformational diversity of G quadruplex which can act as extra base for stacking (Figure 4E). We found 10,435 SNVs across the combined COSMIC and CLINVAR mutations that increase the stability (lower the MFE relative to the reference sequence) while 12,061 SNVs brought no change to the MFE. An additional 15,019 variants destabilize the G4. Transversions were more likely to change the structure of the G quadruplex region without disrupting the G stacks and increasing the thermodynamic stability of the structure (17%) compared to transitions (10%). Additionally, transition mutations were found to destabilize the G4 structure at a higher rate (22%) compared to transversions (17%) (Table 1).

**Table 1.** Count and proportion of effect of type of mutation on stability of G4 (COSMIC database).

| Type of SNV | Effect of SNV on MFE | Freq | Percentage |
| --- | --- | --- | --- |
| Transition | Destabilized | 8,600 | 22.93 |
| Transversion | Further stabilized | 6,603 | 17.60 |
| Transition | no change | 6,552 | 17.46 |
| Transversion | Destabilized | 6,419 | 17.11 |
| Transversion | no change | 5,509 | 14.68 |
| Transition | Further stabilized | 3,832 | 10.21 |

**Figure 4.**
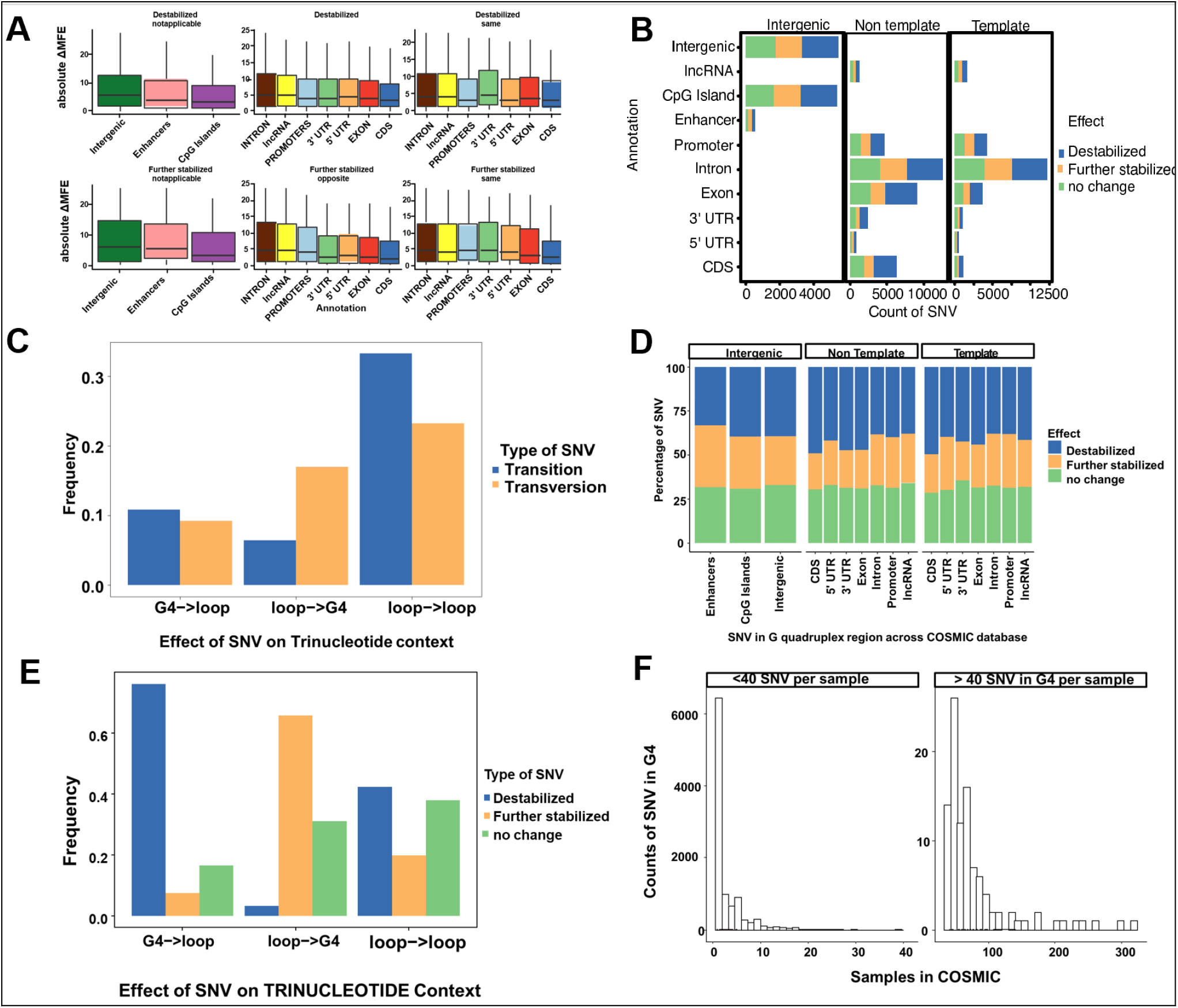
Thermodynamic changes associated with variants in various genomic features. (A) Non-zero delta MFE of G4 across different annotation for destabilized and further stabilized effect by SNV. (B) Count of variants across different regions of the genome and in strand specific or alternative to the coding gene. (C) Proportion of effect on G4 based on the type of mutation. (D) Proportion of variants across different annotation. (E) Proportion of trinucleotide context based on the type of effect on the G4 sequence. (F) Histogram of variants by sample.

### Variants in transcript regions

We find comparatively higher number of mutations in G4 forming exonic regions in 5’UTR, 3’ UTR and CDS regions of protein coding genes when the G4 in formed in the strand opposite the transcribed gene (Figure 3C). The count of SNV around G4 forming regions in intron and promoter regions were proportionate with the transcript opposite or in same strand as the transcript. This shows selection pressure of variants around exon regions as compared to the non-coding regions. Previously, it has been hypothesized the formation of G4 in either strand within the transcribed region, along with nascent RNA would lead to formation of DNA:RNA hybrid R loops in the G quadruplex which results in physically halting the polymerase movement inhibiting further rounds of transcription (41). Additionally, G4 formed on the non-template strand could interfere with the reannealing of the DNA strands increasing the stability of the R loop hybrid.

### Gene component variants

Comparing mutations in different functional groups, G→A mutations are elevated in exons (35.18%) and decreased proportion in promoter region (26.87%). We find a lower percentage of T→G mutations in G4 regions occurring in exons (11.74%) compared to intron, promoter (18%), enhancers (29.84%) and intergenic regions (18%). This pattern of low T→G variants coincides with counts in the CDS region while the 5’ UTR show increased T→G variants (16%). G→A SNVs are found less in enhancers (19%) which are distant from the transcription site and deamination occurring in upstream of transcription site does not affect the G4 region but comparatively have the highest proportion of T→G (29.84%) mutations (Table 2).

**Table 2.**
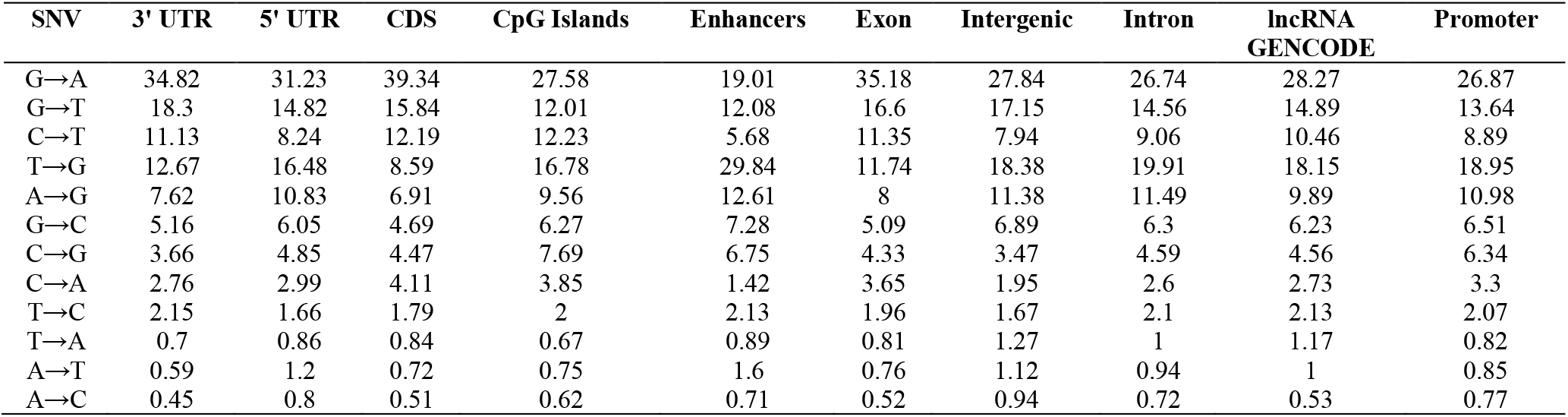
Proportion of SNV by annotation.

| SNV | 3' UTR | 5' UTR | CDS | CpG Islands | Enhancers | Exon | Intergenic | Intron | lncRNA<br>GENCODE | Promoter |
| --- | --- | --- | --- | --- | --- | --- | --- | --- | --- | --- |
| G→A | 34.82 | 31.23 | 39.34 | 27.58 | 19.01 | 35.18 | 27.84 | 26.74 | 28.27 | 26.87 |
| G→T | 18.3 | 14.82 | 15.84 | 12.01 | 12.08 | 16.6 | 17.15 | 14.56 | 14.89 | 13.64 |
| C→T | 11.13 | 8.24 | 12.19 | 12.23 | 5.68 | 11.35 | 7.94 | 9.06 | 10.46 | 8.89 |
| T→G | 12.67 | 16.48 | 8.59 | 16.78 | 29.84 | 11.74 | 18.38 | 19.91 | 18.15 | 18.95 |
| A→G | 7.62 | 10.83 | 6.91 | 9.56 | 12.61 | 8 | 11.38 | 11.49 | 9.89 | 10.98 |
| G→C | 5.16 | 6.05 | 4.69 | 6.27 | 7.28 | 5.09 | 6.89 | 6.3 | 6.23 | 6.51 |
| C→G | 3.66 | 4.85 | 4.47 | 7.69 | 6.75 | 4.33 | 3.47 | 4.59 | 4.56 | 6.34 |
| C→A | 2.76 | 2.99 | 4.11 | 3.85 | 1.42 | 3.65 | 1.95 | 2.6 | 2.73 | 3.3 |
| T→C | 2.15 | 1.66 | 1.79 | 2 | 2.13 | 1.96 | 1.67 | 2.1 | 2.13 | 2.07 |
| T→A | 0.7 | 0.86 | 0.84 | 0.67 | 0.89 | 0.81 | 1.27 | 1 | 1.17 | 0.82 |
| A→T | 0.59 | 1.2 | 0.72 | 0.75 | 1.6 | 0.76 | 1.12 | 0.94 | 1 | 0.85 |
| A→C | 0.45 | 0.8 | 0.51 | 0.62 | 0.71 | 0.52 | 0.94 | 0.72 | 0.53 | 0.77 |

Previously, higher counts of C→T over G→A variants were identified in the non-template strand, which was hypothesized due to cytosine deamination in the nearby 2kb downstream of 5’ end of genes due to higher exposure of single stranded DNA (42). However, we predict the implication of these variants occuring within G quadruplex regions and cause a conformational shift in its structure leading to alteration in expression and binding patterns across these regions. Additionally, 8-oxoguanine formation in G quadruplex binding Sp1 proteins is an important regulator for adipose tissue development and GC rich promoter region with transcription factor sites activating proportional to increasing 8 oxo-G abundance (43).

### Enrichment analysis

#### Gene Ontology

Gene Ontology (GO) enrichment analysis was performed for biological processes (GO:BP) and cellular components (GO:CC). A total of 424 GO:BP categories were determined to be significant (FDR ≤ 0.05) overall (Supplemental Figure 1; Supplemental Table 5), while 425 significant GO:BP enrichments were found for COSMIC alone (Supplemental Figure 2; Supplemental Table 6) and 48 were found for CLINVAR (Supplemental Figure 3; Supplemental Table 7). When this was further broken down into mutations resulting in a loss of a G4, we found 205 significant GO:BP overall (Supplemental Table 8), 75 for COSMIC (Supplemental Table 9) and 25 for CLINVAR (Supplemental Table 10). Among the COSMIC enrichments were synapse organization, axonogenesis, neuron projection guidance, axon guidance, cell-substrate adhesion, neuromuscular process, regulation of neuronprojection development, and xenobiotic glucuronidation. One example gene is the App transcript, which is involved in synapse formation and function in the developing brain. The App transcript is transported to neuronal dendrites, where the transmembrane APP protein plays an integral role in synapse formation and function. However, the translation of App is repressed by the binding of the Fragile X Mental Retardation protein (FMRP) to G-quadruplexes in the App coding region. This repression is thought to occur through direct interaction with the ribosomes, resulting in stalled ribosomal progression on the mRNA (44). Past studies have also shown that this repression can be relieved by synaptic activation of metabotropic glutamate receptors, specifically mGluR5 receptors. This results in the release of FMRP and an increase in APP translation (45).

The CLINVAR enrichments included a number of muscular-related processes, such as striated muscle contraction, neuromuscular process, actin-mediated cell contraction, cardiac conduction, cardiac muscle cell action potential, cardiac muscle cell contraction, membrane depolarization, regulation of actin filament-based movement, muscle tissue morphogenesis, muscle organ morphogenesis, regulation of heart rate, regulation of action potential, cardiac muscle cell action potential involved in contraction, cell communication involved in cardiac conduction, regulation of striated muscle contraction, sensory perception of sound, regulation of heart rate by cardiac conduction, musculoskeletal movement, multicellular organismal movement, transmission of nerve impulse, cardiac muscle tissue morphogenesis, skeletal muscle contraction, and ventricular cardiac muscle cell action potential. Variants leading to a gain of a G4 result in 115 GO:BP enrichments overall (Supplemental Table 11), 22 for COSMIC (Supplemental Table 12) and 2 for CLINVAR (Supplemental Table 13). Among the COSMIC enrichments from genes gaining G4 due to the variants are positive regulation of transcription by RNA polymerase II and actin cytoskeleton organization while the CLINVAR enrichments genes based on loss of G4 include system development, action potential, and cardiac muscle cell action potential. Loss of G4 using COSMIC resulting in similar enriched GO terms as did G4 loss in CLINVAR, included muscle contraction, muscle system process, cardiac muscle contraction, striated muscle contraction, heart contraction, heart process, cardiac muscle cell contraction, cardiac muscle cell action potential involved in contraction, actin-mediated cell contraction, actin filament-based movement, cardiac muscle cell action potential, regulation of heart contraction, action potential, and multicellular organismal signaling.

Among the enriched categories detected were PDZ domain proteins (GIPC2, GRIDZIP, LIMK2, PDLIM7, PDZD7, WHRN, SIPA1L3, PRX, MYO1BA, MAGI2, and MAST) with G4 in coding regions and variants affecting the RGG (arginine-glycine-glycine) domain or G quadruplex stability negatively. Proteins with RGG repeats have been known to bind to G4 structures. Variants in these regions affecting the G4 stability further could affect downstream binding.

GO:CC enrichments yield 128 significant categories overall (129 for COSMIC and 14 for CLINVAR) (Supplemental Figures 4-6; Supplemental Tables 14-16). Among the enriched GO:CC categories detected in COSMIC are collagen containing extracellular matrix, and cell-cell contact zone indicating mutations in these genes affect the adhesion of cells to the extracellular matrix. Other enriched GO:CC terms in CLINVAR include I band, sarcolemma, and myofilament Z disc. Enriched GO:CC terms from Loss of G4 using CLINVAR database include collagen trimer and PCSK9-LDLR complex.

#### KEGG metabolic pathways

KEGG enrichment yielded 96 significant pathways overall as well as 91 COSMIC and 11 CLINVAR (Supplemental Figures 7-9; Supplemental Tables 17-19). Those leading to a loss of G4 yielded 33 significant categories, including 12 and 5 for COSMIC and CLINVAR, respectively (Supplemental Tables 20-22). Among the enriched categories for genes with loss of G4 within CLINVAR are hypertrophic cardiomyopathy, dilated cardiomyopathy, arrhythmogenic right ventricular cardiomyopathy, adrenergic signaling in cardiomyocytes, and acute myeloid leukemia. KEGG enrichments for a gain of G4 resulted in 31, 3, and 0 for overall, COSMIC and CLINVAR respectively (Supplemental Tables 23-24). The enriched terms from gain of G4 in CLINVAR variants are melanoma, phospholipase D signaling pathway and cocaine addiction.

#### INTERPRO protein domains

INTERPRO enrichment yielded 23, 23, and 2 enrichments (Supplemental Tables 25-27). Included were Src homology-3 domain (n=69 FDR=3.73E-03) and Pleckstrin homology-like domain (PH) (n=147, FDR=1.30E-10). The binding affinity of PH domains with the exception for some binding phosphoiniositides with high affinity, majority have unique recognition domains and are known for functional plasticity (46,47).

#### Transcription Factors

We identified the enrichment of transcription factors (TFs) including NFKB1, ZFX, MBD3, ASX1, SUZ12, NCOR1, HMGN3, USF2, EGR1, GTF2F1, KDM4B, HNRNPH1, HNRNPL, NONO, TARDBP, NFATC3, KDM3A, and HOXA3 among others (Supplemental Figures 10-12; Supplemental Table 28). The majority (92%) of these had at least a G quadruplex in their gene structure working in a feed forward regulation of genes. We identified these variants break the motifs for transcription binding sites.

### Trinucleotide context mutation in G quadruplex sequence

Based on the nucleotide context one base pair before and after the mutation, we identified 79% of the variants to be affecting the loop region and 23% of the SNV after the change leads to the formation of GGG in regions with G(A|C|T)G. We find 36% (n=6,810) of the transversion mutations are T→G, while 21% (n=4,070) of the transitions are A→G (Figures 5A-5D). This change becomes more prominent, T→G mutations occurring in context of GTG→GGG occurs in 14% of the SNVs leading to formation of stable G tetrad while GAG→GGG occurs as 6% of the variants. Interestingly, the destabilization of GGG region occurs by GGG→GAG transition in 11% of SNVs (Figure 5E). Previously, it has been reported that the GGG exhibits context dependent specific mutational patterns that preserve the potential for G4 formation (48). We find G→A mutations to be approximately 29% of the total SNVs in the selected G quadruplex regions, with 26% (n=4,144; 11% of the total) of those variants occurring in a context of GGG→GAG with implications of alteration mechanism for G quadruplex sequences (Figure 5E). We observe these patterns throughout different noncoding annotations, except exonic regions and CDS regions. We identify an increased propensity to be able to form stable multiple conformations with de-stabilized structures for 25% of the sequences with the variants while 14% of the variants incurred no change to the stability of the structure (Supplemental Table 29). This approach of analyzing the probable base pairing alternatives for additional guanine Hoogsteen base pairing can help identify the effects of variants within the G4 structure and hence predict the structure change and functionality of G quadruplex in various molecular processes.

**Figure 5.**
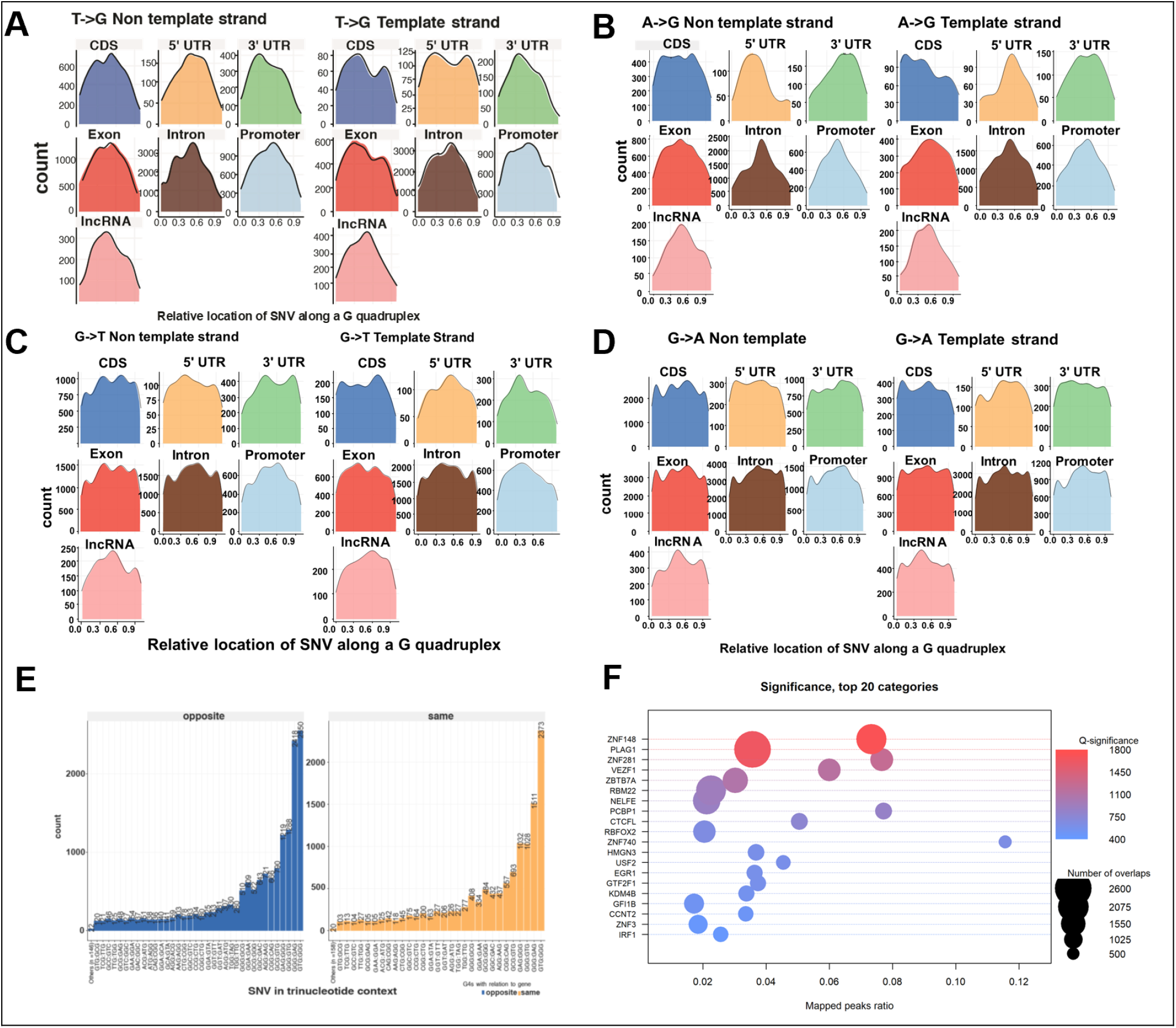
Distribution of SNVs across the G4 regions on the non-template and template strand. Shown are the results for (A) T→G variants; (B) A→G variants; (C) G→T variants; and (D) G→A variants. (E) Distribution of SNVs in trinucleotide contexts relative to the opposite or same strand as the corresponding gene. (F) Significance of the top 20 transcription factors and their genome-wide binding sites.

Based on the position of the mutation in the G quadruplex from the starting point and the length of the sequence, the normalized position for each variant in the G quadruplex was calculated. The relative location of a variant in a G4 is defined as the position of variant divided by the length of the G4. For single nucleotide variants mutating to G either from A or T, we find similar elevated patterns in the center of the G quadruplex. T|A→G mutations show conservation of guanine in the center region with the exception of the CDS and exon in both template and non-template strand across both COSMIC and CLINVAR databases (Figure 5E, Supplemental Figures 13-16). These changes are stricter for SNVs within the 5’UTR across the template and non-template strand in the CLINVAR database, where we observe mutations in the relative center of the G quadruplex for T→G variants as compared to the 5’ UTR COSMIC mutations where we observe mutations across the two extreme loops compared to the center. A→G mutations are observed in a higher proportion at the beginning of G quadruplex in CDS region which provides evidence for mutation pressure in the coding region preferentially protecting the coding sequence. G quadruplexes in UTRs have been reported to be under selection pressure and variants in G4 can account for instability in G4 and diseases (49).

## DISCUSSION

### Variants involved in oxidation

High occurrences of oxidized guanine in G quadruplex structures compared to duplex DNA has been previously established (50). The mutation has been suggested to occur around the external tetrads compared to the central tetrad due to radical trapping antioxidants that slow the efficiency of mutation (51). We identify an increase in counts at the middle stacking of the G quadruplex for A|T→G, implying the functional impact of types of specific variants towards the conformation of the G quadruplex. The observed elevation in counts can be accounted for the presence of tetrads available for variants from guanine to A, C and T. G quadruplex with spare tires can also form alternate structures or exclusion of certain guanosines in case of lesions or substitution in one tetrad region. Base excision repair with APP1 and OGG1 at the promoter of VEGF has shown this mechanism for formation of G quadruplex and this suggests formation of G quadruplex for other genes through a similar mechanism (52,53). Oxidative stress occurring due to the reactive oxygen species (ROS) affects the genome stability and promote mutagenesis, senescence, and other age-related diseases (54). Mutations in GGG regions can destabilize the stacking of guanines, altering the ionization potential affecting the ability of the G region to be further oxidized. G→A, T or C mutations can disrupt the stacking while mutations to G can further stabilize the G quadruplex or allow additional conformations for the stacking. We investigated the change of each type of SNV in each annotation to have the highest change. Based on absolute ΔMFE based on the change, we find pG4 in CDS region and CpG region are least prone to the variants while enhancers and Intergenic G4 are prone to higher stabilizing and destabilizing due to the variants (Figure 4A). G quadruplexes in 3’ UTR in the same strand of coding genes along with introns are prone to the variants and are highly stabilized or destabilized by the variants occurring in COSMIC.

We investigated which SNVs in each annotation had the highest change. The 3’ UTR has a higher incidence of T→G versus A→G SNVs. This implies that T→G mutations are more likely to stabilize G quadruplexes found in the 3’UTR. Putative G4s in CDS and CpG regions are least prone to variants while enhancers and intergenic G4 are show higher changes in stabilization (both stabilizing and destabilizing) due to the SNVs (Figure 4A, Supplemental Figure 17).

### Role of location of SNvs in G4s

The relative position of G→T substitutions along G quadruplex sequences is shown in Figure 5C. The location of this mutation at the beginning of the G quadruplex can disrupt the structural formation; however, further elevated peaks at varying locations leading to additional guanines across the G4 may introduce additional tetrads in introns and exons (Figures 5C and 5D).

The observation of an increased number of G quadruplex stacks resulting from G→T|A substitutions that break up longer runs of G’s is consistent with studies that oxidation of the multiple G’s occur at the start of the G quadruplex tetrads. Our results help establish that the location of mutations and the type of mutation in G rich regions alter the shape and stability of the G quadruplex structure. Previously it has been established that the most sensitive sites are located at the center tetrad (55). For mutations in CLINVAR, we observe a higher mutation rate at the start of G4. The A→G mutations associated with COSMIC variants show a considerable difference in their location relative to the G4 position (Supplemental Figure 18). The escape of 8-oxoG from DNA repair during DNA replication can cause the misincorporation of adenine opposite 8-oxoG leading to the addition of T in place of G. For instance, a sequence with GTTAGGG with 8-oxoG at its fifth position, a mis-incorporation of the A occurs opposite G. Due to the presence of consistent Gs in the region, the true proportions of change in these regions can be hard to monitor over a range of replications. Methylation of cytosine leads to formation of 5-methyl cytosine which are residues for spontaneous transitions. Cytosine deamination might be the primary cause of C→T transition. Further, based on the context, a high proportion of T→G mutations lead to a GTG→GGG structure, supporting the stability of the G quadruplex. It presents a question of whether T→G mutations confer additional stability of G4 in cancer cells. Past studies have highlighted the conditional impact of OG mutations in base pairing with A in mutagenic MutY homolog harboring increased G→T transversions in MUTYH leading to a higher incidence rate of colorectal cancer (56-58). Thymine glycol are non-mutagenic lesions which are highly mutagenic and in regions of DSBs, are cytotoxic. *In vitro* studies have shown it to block replicative and repair DNA polymerases (59).

The OG, thymine glycol and abasic sites formed are repaired by the excision repair pathway. The difference in repair of oxoG sites have been observed in NEIL glycolyases which have been known to remove guanidinohydantoin (Gh) and spiroiminodihydantoin (Sp) from G quadruplex structures in promoter region over parallel conformation (60). However, the glycolysases were not able to remove the oxoG structure from the telomeric G quadruplex or the same G quadruplex structure in antiparallel structures.

### TERT G4 mutations

A study has highlighted that the entire 67 bp G quadruplex associated with the TERT promoter was found to be completely protected from DNase cleavage while the version containing G→T variants was found to be degraded into discrete segments (61,62). Additionally, this region folds into a compact G4 structure without any hairpins in between the G quadruplex stacks. However, based on DMS footprinting studies, formation of hairpins has been predicted (63). Overall, we identified 52 possible SNVs in 39 base pair locations in this 67 bp G quadruplex. The SNV chr5:1,295,113 (G→T) located in the TERT region is present around a G quadruplex in the non-template strand. The SNV was associated with more than twenty-two cancer types including central nervous system, liver, bladder, ovarian, breast, kidney lung, bone, pancreatic, among others. Many of these SNVs destabilize G quadruplexes. Further, with nine tetrads (GGG repeats present) multiple G4 can potentially be formed. With a SNV (G→A), We find the G quadruplex stability with the variant to differ if alternate G4 tetrads are used for the stacking.

### Transcription factor binding

Transcription factor proteins (TFs) known to bind G rich regions including SP1 (64), NF-κB (65), CREB (66), and the methyl-CpG binding domain MBD of methyl-CpG binding protein 2 (MeCP2) (67) had decreased association constants up to 10 fold for transcription factor sites with change of guanine to 8-oxoguanine in model duplex DNA with the donor acceptor pattern change on the imidazole ring in guanine compared to OG. The structure change for guanine for association with CREB was found have a role in epigenetic repression (66). This is supported with our results highlighting reversal of these sequences to a stabilized G4 by change through T→G region in cancer cells. For instance, the variant chr10:122,143,482: G→A significantly affects the binding sites of TFs NHLH1, FOXO3, TAL1, TP53, HES5, HES7, USF2, EGR3, ZNF740, and SP1 among others (Supplemental Table 6). We observe similar observations for an additional 424 SNVs which occur in at least five cancer types in the COSMIC database and disrupt the TF binding site with an average of 15.1 TF per variant.

Local network cluster (STRING) analysis of the enriched TFs yielded terms related to PRC1 complex (4/12 FDR 0.00049) and PcG protein complex (6/25, FDR 6.22e-06), PcG protein complex, and positive regulation of histone H3-K27 methylation (11/59 FDR 1.82 e-10). Polycomb repressive Complex (PRC1) engage in transcriptional control through chromatin modification with histone 2A through a protein ligase Ubiquitylation (68,69). Although the mechanism of PRC1 is under active investigation, recent evidence suggests role of G tracts to selectively remove PCR2 complex from genes during gene activation (70). Polycomb complexes have been associated with repression to maintain cell identity but are associated with actively transcribed loci, and this evidence suggest direct role of G quadruplexes across cell types to regulate expression through structural variation. GO cellular component analysis for the TFs found enriched terms related to Brahma complex, (3/3 FDR 0.00079), Ino80 complex (5/15 FDR 5.35e-05) which are different complexes associated with chromatin remodelling.

Different repair mechanisms including BER, and mismatch repair are required for protecting non-canonical or mismatch base pairs due to polymerase error. Neurogenerative disorders occurring through expansion of CAG→CTG repeats have been associated with MutSβ, a heterodimer involved in mismatch repair. Though the involvement of G quadruplexes in gene transcription and telomere regulation has been studied and proven, the mechanism of base excision repair by DNA glycosylases in G quadruplex and other non-canonical structures is poorly understood. We identified G quadruplexes with SNVs in the genes of CHRNG, GRIN2C, CHAT, ADCY1, GABRG3, CACNG3, PPFIA3, LRTOMT, VAMP2, TSPOAP1, MAPK3,GABRR2, KCNJ6, PICK1, and STX1A, among others. These genes have been associated with several psychiatric disorders, schizophrenia, Bipolar Disorder, Tobacco Use Disorder, Parkinson’s disease, and autism.

Previous research has shown the presence of G quadruplex sequences in various untranslated dendritic mRNAs suggesting the role of G quadruplexes as a neurite localization signal. Deletion of different putative G quadruplex sequence led to severe loss of signal in neurites. It has been hypothesized that the G quadruplex structure being sensitive to cationic, can function in correlation to the neuronal activity in localization and transport as activity dependent changes. Cationic sensitivity could influence the stability and structure and regulate the binding of trans-acting factors (71).

## CONCLUSION

G quadruplexes are formed because of an intricate balance between the folding energy by a nick in the DNA, methylated guanines, and guanines available for stacking. The balance between the hypomethylated and hypermethylated G rich regions near promoters (despite cytosine deamination and cytosine methylation) results in the preserved regions of CpG islands are observed across mammalian genome (72). These previously identified regions as CpG islands can be the preserved G quadruplex regions. Further, methylated guanines CpG islands have been identified within the genes (73) and the methylation susceptibility constraints the G quadruplex formation. We hypothesize these methylation and oxidation patterns are one mechanism by which G quadruplexes can preserve their sequence conformation and the variants occurring in these regions alter the molecular functions downstream.

With the introduction of next generation techniques for identification of G quadruplexes, analysis of variants in these complex region and mechanism of formation of G-quadruplex in different cell types remains uncertain. Our study points out a subset of different genes and G quadruplexes sequences which are affected in cancer cells and consequences of secondary structure forming regions with a nucleotide level investigation. G quadruplex formed in genomic regions participate in gene regulatory pathways to alter gene expression and downstream pathways. Based on large accumulation of published studies, we identify the possible effects of these single nucleotide variants occurring on coding and non-coding regions on the stability of G quadruplexes.

## DATA AVAILABILITY

All code and resulting data is available in the GitHub repository https://github.com/UofLBioinformatics/G4_SNV/. UCSC genome browser tracks can be accessed at: https://bit.ly/G4_SNV.

## ACCESSION NUMBERS

Experimentally validated G4 sequences were obtained from the publicly available dataset generated by Chamers et al. (29) which is available in GEO under accession number GSE63874.

## SUPPLEMENTARY DATA

Supplementary Data are available online.

## ACKNOWLEDGEMENT

We wish to thank members of the Kentucky IDeA Networks of Biomedical Research Excellence (KY INBRE) Bioinformatics Core and the Rouchka and Park labs for their valuable feedback.

