## Supplemental Tables and Figures for "Analysis of nucleotide variations in human g-quadruplex forming regions associated with disease states"

**Supplemental Table 1**. Count of SNVs in overall COSMIC database.

| | | TO | | | |
|---|---|---|---|---|---|
| | | **A** | **C** | **G** | **T** |
| **FROM** | **A** | 0 | 94,513 | 2,006,240 | 1,121,852 |
| | **C** | 1,965,512 | 0 | 990,818 | 3,788,310 |
| | **G** | 3,788,966 | 989,193 | 0 | 2,017,502 |
| | **T** | 1,124,289 | 2,007,011 | 951,101 | 0 |

**Supplemental Table 2**. Counts of SNVs in G4 regions from the COSMIC database.

| | | TO | | | |
|---|---|---|---|---|---|
| | | **A** | **C** | **G** | **T** |
| **FROM** | **A** | 0 | 601 | 7,191 | 768 |
| | **C** | 2,529 | 0 | 2,937 | 8,094 |
| | **G** | 21,349 | 4,450 | 0 | 10,791 |
| | **T** | 742 | 1,608 | 8,861 | 0 |

**Supplemental Table 3.** Changes in putative G4 from the COSMIC database across both strands before and after mutation. (0: absence of pG4; 1: presence of pG4 in forward strand; -1: presence of pG4 in reverse strand)

| G4 with reference allele | G4 with alternate allele | count | % | G4hunter score (reference) | Sd (reference) | G4hunter score (alternate) | Sd (alternate) |
|---|---|---|---|---|---|---|---|
| -1 | -1 | 13,793 | 36.77 | -1.222 | 0.387 | -1.219 | 0.4 |
| -1 | 0 | 3,581 | 9.55 | -1.026 | 0.351 | -0.901 | 0.345 |
| -1 | 1 | 6 | 0.02 | 0.175 | 0.37 | 0.279 | 0.36 |
| 0 | -1 | 1,354 | 3.61 | -1.082 | 0.356 | -1.196 | 0.359 |
| 0 | 1 | 1,374 | 3.66 | 1.085 | 0.368 | 1.201 | 0.371 |
| 1 | 0 | 3,655 | 9.74 | 0.993 | 0.359 | 0.871 | 0.355 |
| 1 | 1 | 13,753 | 36.66 | 1.222 | 0.376 | 1.219 | 0.391 |

**Supplemental Table 4.** Count and proportion of variants in experimentally validated G4 regions for different functional regions.

| | COSMIC | | CLINVAR | |
|---|---|---|---|---|
| **Annotation** | **Count** | **Frequency** | **Count** | **Frequency** |
| **CDS** | 7,569 | 10.02 | 2,281 | 45.38 |
| **5' UTR** | 1,514 | 2 | -- | -- |
| **3' UTR** | 3,669 | 4.86 | 179 | 3.56 |
| **EXON** | 13,034 | 17.26 | 3,014 | 59.97 |
| **INTRON** | 26,014 | 34.44 | 1,251 | 24.89 |
| **PROMOTER** | 9,248 | 12.24 | 554 | 11.02 |
| **ENHANCER** | 563 | 0.75 | -- | -- |
| **CpG ISLAND** | 5,356 | 7.09 | -- | -- |
| **GENCODE lncRNA** | 3,121 | 4.13 | 700 | 13.92 |
| **INTERGENIC** | 5,441 | 7.2 | 761 | 15.14 |

**Supplemental Table 5**. Significant GO:BP enrichments for all COSMIC and CLINVAR G4 mutations.

| GO ID | GO Description | Universe | COSMIC and CLINVAR | Adjusted P-value |
|---|---|---|---|---|
| **GO:0007399** | nervous system development | 1648 | 1087 | 3.01E-48 |
| **GO:0048856** | anatomical structure development | 4152 | 2427 | 6.54E-48 |
| **GO:0032502** | developmental process | 4584 | 2644 | 1.02E-46 |
| **GO:0048731** | system development | 2976 | 1790 | 1.56E-42 |
| **GO:0007275** | multicellular organism development | 3266 | 1938 | 2.46E-41 |
| **GO:0009653** | anatomical structure morphogenesis | 1871 | 1181 | 1.16E-38 |
| **GO:0048699** | generation of neurons | 969 | 670 | 1.07E-37 |
| **GO:0000902** | cell morphogenesis | 718 | 520 | 4.07E-37 |
| **GO:0030154** | cell differentiation | 2860 | 1706 | 5.88E-37 |
| **GO:0048869** | cellular developmental process | 2879 | 1715 | 1.04E-36 |
| **GO:0022008** | neurogenesis | 1096 | 736 | 6.89E-35 |
| **GO:0030182** | neuron differentiation | 923 | 636 | 7.71E-35 |
| **GO:0048468** | cell development | 1355 | 877 | 4.69E-33 |
| **GO:0032501** | multicellular organismal process | 5319 | 2948 | 8.54E-33 |
| **GO:0048858** | cell projection morphogenesis | 459 | 352 | 1.36E-32 |
| **GO:0048666** | neuron development | 729 | 516 | 2.16E-32 |
| **GO:0032989** | cellular component morphogenesis | 549 | 407 | 3.74E-32 |
| **GO:0030030** | cell projection organization | 1164 | 766 | 7.19E-32 |
| **GO:0120039** | plasma membrane bounded cell projection morphogenesis | 455 | 348 | 9.12E-32 |
| **GO:0031175** | neuron projection development | 656 | 471 | 9.40E-32 |
| **GO:0120036** | plasma membrane bounded cell projection organization | 1144 | 754 | 1.22E-31 |
| **GO:0032990** | cell part morphogenesis | 469 | 355 | 8.10E-31 |
| **GO:0048812** | neuron projection morphogenesis | 441 | 337 | 1.53E-30 |
| **GO:0000904** | cell morphogenesis involved in differentiation | 495 | 367 | 1.01E-28 |
| **GO:0023051** | regulation of signaling | 2733 | 1601 | 2.82E-28 |
| **GO:0010646** | regulation of cell communication | 2727 | 1596 | 7.06E-28 |
| **GO:0051128** | regulation of cellular component organization | 1929 | 1170 | 3.23E-27 |
| **GO:0061564** | axon development | 323 | 256 | 5.08E-27 |
| **GO:0030029** | actin filament-based process | 717 | 495 | 1.15E-26 |
| **GO:0050793** | regulation of developmental process | 1810 | 1101 | 5.46E-26 |
| **GO:0048667** | cell morphogenesis involved in neuron differentiation | 381 | 291 | 6.02E-26 |
| **GO:0007409** | axonogenesis | 298 | 237 | 3.00E-25 |
| **GO:0023052** | signaling | 5190 | 2837 | 8.54E-25 |
| **GO:0007154** | cell communication | 5221 | 2844 | 1.92E-23 |
| **GO:0016043** | cellular component organization | 5370 | 2912 | 1.84E-22 |
| **GO:0051239** | regulation of multicellular organismal process | 2117 | 1248 | 3.72E-22 |
| **GO:0048513** | animal organ development | 2176 | 1277 | 1.06E-21 |
| **GO:0030036** | actin cytoskeleton organization | 637 | 435 | 1.52E-21 |
| **GO:0035556** | intracellular signal transduction | 2168 | 1267 | 1.65E-20 |
| **GO:0007155** | cell adhesion | 1216 | 754 | 1.02E-19 |
| **GO:0009966** | regulation of signal transduction | 2464 | 1416 | 2.43E-19 |
| **GO:0007010** | cytoskeleton organization | 1309 | 799 | 2.62E-18 |
| **GO:0031344** | regulation of cell projection organization | 467 | 326 | 7.18E-18 |
| **GO:0050804** | modulation of chemical synaptic transmission | 252 | 195 | 8.78E-18 |
| **GO:0120035** | regulation of plasma membrane bounded cell projection organization | 453 | 317 | 1.42E-17 |
| **GO:0071840** | cellular component organization or biogenesis | 5539 | 2962 | 1.45E-17 |
| **GO:0099177** | regulation of trans-synaptic signaling | 253 | 195 | 1.99E-17 |
| **GO:0048522** | positive regulation of cellular process | 4704 | 2544 | 2.87E-17 |
| **GO:0099536** | synaptic signaling | 522 | 356 | 5.07E-17 |
| **GO:0048523** | negative regulation of cellular process | 3896 | 2135 | 6.91E-17 |
| **GO:0034330** | cell junction organization | 518 | 353 | 8.99E-17 |
| **GO:0010975** | regulation of neuron projection development | 288 | 215 | 1.91E-16 |
| **GO:0022603** | regulation of anatomical structure morphogenesis | 700 | 456 | 2.19E-16 |
| **GO:0045595** | regulation of cell differentiation | 1129 | 693 | 2.71E-16 |
| **GO:0007417** | central nervous system development | 584 | 389 | 4.80E-16 |
| **GO:0050794** | regulation of cellular process | 9523 | 4886 | 6.17E-16 |
| **GO:0048518** | positive regulation of biological process | 5294 | 2827 | 8.88E-16 |
| **GO:0099537** | trans-synaptic signaling | 501 | 340 | 1.25E-15 |
| **GO:0098916** | anterograde trans-synaptic signaling | 495 | 336 | 1.93E-15 |
| **GO:0007268** | chemical synaptic transmission | 495 | 336 | 1.93E-15 |
| **GO:0007165** | signal transduction | 4776 | 2566 | 2.30E-15 |
| **GO:0009987** | cellular process | 14783 | 7306 | 2.42E-15 |
| **GO:0048583** | regulation of response to stimulus | 3327 | 1835 | 3.42E-15 |
| **GO:0097485** | neuron projection guidance | 169 | 137 | 5.04E-15 |
| **GO:0007411** | axon guidance | 169 | 137 | 5.04E-15 |

**Supplemental Table 5 (continued).**

| GO ID | GO Description | Universe | COSMIC and CLINVAR | Adjusted P-value |
|---|---|---|---|---|
| **GO:0065007** | biological regulation | 10721 | 5447 | 5.19E-15 |
| **GO:0048519** | negative regulation of biological process | 4381 | 2365 | 6.60E-15 |
| **GO:0065008** | regulation of biological quality | 2937 | 1634 | 6.88E-15 |
| **GO:0032879** | regulation of localization | 1615 | 947 | 7.50E-15 |
| **GO:0051716** | cellular response to stimulus | 5982 | 3159 | 9.70E-15 |
| **GO:0050789** | regulation of biological process | 10085 | 5143 | 1.21E-14 |
| **GO:0007267** | cell-cell signaling | 1269 | 761 | 1.63E-14 |
| **GO:0051094** | positive regulation of developmental process | 967 | 595 | 6.68E-14 |
| **GO:0051179** | localization | 4343 | 2335 | 1.81E-13 |
| **GO:0050770** | regulation of axonogenesis | 100 | 88 | 2.65E-13 |
| **GO:0051049** | regulation of transport | 1327 | 786 | 3.15E-13 |
| **GO:0048870** | cell motility | 1362 | 804 | 4.26E-13 |
| **GO:0007167** | enzyme-linked receptor protein signaling pathway | 795 | 497 | 5.75E-13 |
| **GO:0009887** | animal organ morphogenesis | 582 | 378 | 7.07E-13 |
| **GO:0050808** | synapse organization | 275 | 198 | 3.52E-12 |
| **GO:0051960** | regulation of nervous system development | 257 | 187 | 3.93E-12 |
| **GO:0016477** | cell migration | 1210 | 717 | 7.09E-12 |
| **GO:0009888** | tissue development | 1239 | 732 | 8.83E-12 |
| **GO:0003012** | muscle system process | 305 | 215 | 8.94E-12 |
| **GO:0055085** | transmembrane transport | 1060 | 635 | 1.60E-11 |
| **GO:0051130** | positive regulation of cellular component organization | 844 | 518 | 1.69E-11 |
| **GO:0006810** | transport | 3620 | 1956 | 1.92E-11 |
| **GO:0065009** | regulation of molecular function | 2121 | 1192 | 2.01E-11 |
| **GO:0040011** | locomotion | 1075 | 641 | 4.38E-11 |
| **GO:0098660** | inorganic ion transmembrane transport | 622 | 394 | 5.04E-11 |
| **GO:0042391** | regulation of membrane potential | 319 | 221 | 7.21E-11 |
| **GO:0051234** | establishment of localization | 3775 | 2029 | 7.97E-11 |
| **GO:0007015** | actin filament organization | 395 | 265 | 8.57E-11 |
| **GO:0045944** | positive regulation of transcription by RNA polymerase II | 952 | 573 | 1.11E-10 |
| **GO:0007420** | brain development | 384 | 258 | 1.44E-10 |
| **GO:0098655** | cation transmembrane transport | 656 | 411 | 1.57E-10 |
| **GO:0023057** | negative regulation of signaling | 1067 | 634 | 1.61E-10 |
| **GO:0061061** | muscle structure development | 407 | 271 | 1.72E-10 |
| **GO:0007169** | transmembrane receptor protein tyrosine kinase signaling pathway | 511 | 330 | 1.77E-10 |
| **GO:0034220** | ion transmembrane transport | 816 | 498 | 2.25E-10 |
| **GO:0016310** | phosphorylation | 1444 | 833 | 2.28E-10 |
| **GO:1902531** | regulation of intracellular signal transduction | 1429 | 825 | 2.39E-10 |
| **GO:0007264** | small GTPase mediated signal transduction | 389 | 260 | 2.86E-10 |
| **GO:0010648** | negative regulation of cell communication | 1060 | 629 | 2.89E-10 |
| **GO:0050767** | regulation of neurogenesis | 208 | 153 | 3.26E-10 |
| **GO:0060322** | head development | 402 | 267 | 4.05E-10 |
| **GO:0040012** | regulation of locomotion | 842 | 511 | 4.05E-10 |
| **GO:0006936** | muscle contraction | 260 | 184 | 4.79E-10 |
| **GO:0022604** | regulation of cell morphogenesis | 240 | 172 | 4.91E-10 |
| **GO:0098662** | inorganic cation transmembrane transport | 572 | 362 | 6.98E-10 |
| **GO:0097435** | supramolecular fiber organization | 710 | 438 | 7.39E-10 |
| **GO:0006811** | ion transport | 1187 | 694 | 9.42E-10 |
| **GO:0030001** | metal ion transport | 678 | 420 | 9.61E-10 |
| **GO:0048167** | regulation of synaptic plasticity | 122 | 98 | 1.02E-09 |
| **GO:2000026** | regulation of multicellular organismal development | 981 | 584 | 1.04E-09 |
| **GO:0045597** | positive regulation of cell differentiation | 624 | 390 | 1.13E-09 |
| **GO:2000145** | regulation of cell motility | 818 | 496 | 1.14E-09 |
| **GO:0030334** | regulation of cell migration | 767 | 468 | 1.35E-09 |
| **GO:0048585** | negative regulation of response to stimulus | 1287 | 745 | 2.06E-09 |
| **GO:0072359** | circulatory system development | 729 | 446 | 2.71E-09 |
| **GO:1903508** | positive regulation of nucleic acid-templated transcription | 1297 | 749 | 3.44E-09 |
| **GO:0045893** | positive regulation of DNA-templated transcription | 1297 | 749 | 3.44E-09 |
| **GO:0051173** | positive regulation of nitrogen compound metabolic process | 2565 | 1404 | 3.55E-09 |
| **GO:0051962** | positive regulation of nervous system development | 147 | 113 | 3.93E-09 |
| **GO:0031346** | positive regulation of cell projection organization | 247 | 174 | 3.98E-09 |
| **GO:0006812** | cation transport | 886 | 530 | 4.38E-09 |
| **GO:0009968** | negative regulation of signal transduction | 1009 | 595 | 6.51E-09 |
| **GO:0032970** | regulation of actin filament-based process | 330 | 222 | 6.52E-09 |

**Supplemental Table 5 (continued).**

| GO ID | GO Description | Universe | COSMIC and CLINVAR | Adjusted P-value |
|---|---|---|---|---|
| **GO:0006468** | protein phosphorylation | 1268 | 732 | 6.97E-09 |
| **GO:0071805** | potassium ion transmembrane transport | 185 | 136 | 7.67E-09 |
| **GO:1902680** | positive regulation of RNA biosynthetic process | 1303 | 750 | 8.22E-09 |
| **GO:0031325** | positive regulation of cellular metabolic process | 2522 | 1378 | 1.21E-08 |
| **GO:0044087** | regulation of cellular component biogenesis | 774 | 467 | 1.61E-08 |
| **GO:0044057** | regulation of system process | 392 | 256 | 1.94E-08 |
| **GO:0060560** | developmental growth involved in morphogenesis | 135 | 104 | 2.36E-08 |
| **GO:0048588** | developmental cell growth | 129 | 100 | 3.06E-08 |
| **GO:0032535** | regulation of cellular component size | 256 | 177 | 3.24E-08 |
| **GO:0060284** | regulation of cell development | 315 | 211 | 4.14E-08 |
| **GO:0051056** | regulation of small GTPase mediated signal transduction | 238 | 166 | 4.61E-08 |
| **GO:0098609** | cell-cell adhesion | 743 | 448 | 5.00E-08 |
| **GO:0031589** | cell-substrate adhesion | 290 | 196 | 6.49E-08 |
| **GO:0001667** | ameboidal-type cell migration | 333 | 220 | 1.12E-07 |
| **GO:0099587** | inorganic ion import across plasma membrane | 112 | 88 | 1.40E-07 |
| **GO:0098659** | inorganic cation import across plasma membrane | 112 | 88 | 1.40E-07 |
| **GO:0009893** | positive regulation of metabolic process | 3177 | 1700 | 1.57E-07 |
| **GO:0051254** | positive regulation of RNA metabolic process | 1433 | 810 | 1.66E-07 |
| **GO:0051240** | positive regulation of multicellular organismal process | 1137 | 655 | 1.76E-07 |
| **GO:0050769** | positive regulation of neurogenesis | 119 | 92 | 2.53E-07 |
| **GO:0010604** | positive regulation of macromolecule metabolic process | 2889 | 1553 | 2.57E-07 |
| **GO:0044093** | positive regulation of molecular function | 1305 | 742 | 2.64E-07 |
| **GO:0031324** | negative regulation of cellular metabolic process | 1858 | 1028 | 2.90E-07 |
| **GO:0060627** | regulation of vesicle-mediated transport | 417 | 266 | 2.97E-07 |
| **GO:0045935** | positive regulation of nucleobase-containing compound metabolic process | 1620 | 905 | 3.26E-07 |
| **GO:0006793** | phosphorus metabolic process | 2354 | 1281 | 3.36E-07 |
| **GO:0051641** | cellular localization | 2721 | 1467 | 3.45E-07 |
| **GO:0035725** | sodium ion transmembrane transport | 129 | 98 | 3.60E-07 |
| **GO:0006813** | potassium ion transport | 200 | 141 | 4.40E-07 |
| **GO:0033043** | regulation of organelle organization | 1015 | 588 | 5.17E-07 |
| **GO:1990138** | neuron projection extension | 112 | 87 | 5.18E-07 |
| **GO:0048638** | regulation of developmental growth | 167 | 121 | 5.36E-07 |
| **GO:0006796** | phosphate-containing compound metabolic process | 2336 | 1270 | 5.65E-07 |
| **GO:0042221** | response to chemical | 2943 | 1577 | 5.88E-07 |
| **GO:0001508** | action potential | 117 | 90 | 6.38E-07 |
| **GO:0034329** | cell junction assembly | 341 | 222 | 6.89E-07 |
| **GO:0023056** | positive regulation of signaling | 1380 | 778 | 7.95E-07 |
| **GO:0032956** | regulation of actin cytoskeleton organization | 296 | 196 | 9.18E-07 |
| **GO:0030516** | regulation of axon extension | 61 | 53 | 9.20E-07 |
| **GO:1905114** | cell surface receptor signaling pathway involved in cell-cell signaling | 388 | 248 | 9.30E-07 |
| **GO:0003015** | heart process | 193 | 136 | 9.57E-07 |
| **GO:0048646** | anatomical structure formation involved in morphogenesis | 751 | 446 | 1.03E-06 |
| **GO:0016049** | cell growth | 358 | 231 | 1.03E-06 |
| **GO:0010647** | positive regulation of cell communication | 1374 | 774 | 1.07E-06 |
| **GO:0006996** | organelle organization | 3147 | 1677 | 1.11E-06 |
| **GO:0061387** | regulation of extent of cell growth | 67 | 57 | 1.13E-06 |
| **GO:0051093** | negative regulation of developmental process | 619 | 374 | 1.58E-06 |
| **GO:0007507** | heart development | 338 | 219 | 1.67E-06 |
| **GO:0048589** | developmental growth | 273 | 182 | 1.68E-06 |
| **GO:0003013** | circulatory system process | 443 | 277 | 2.44E-06 |
| **GO:0042127** | regulation of cell population proliferation | 1218 | 690 | 2.85E-06 |
| **GO:0071495** | cellular response to endogenous stimulus | 974 | 562 | 2.94E-06 |
| **GO:0007265** | Ras protein signal transduction | 278 | 184 | 3.20E-06 |
| **GO:0060047** | heart contraction | 187 | 131 | 3.55E-06 |
| **GO:0009719** | response to endogenous stimulus | 1076 | 615 | 3.74E-06 |
| **GO:0060828** | regulation of canonical Wnt signaling pathway | 211 | 145 | 3.96E-06 |
| **GO:0051493** | regulation of cytoskeleton organization | 450 | 280 | 4.00E-06 |
| **GO:0090066** | regulation of anatomical structure size | 328 | 212 | 4.16E-06 |
| **GO:0031327** | negative regulation of cellular biosynthetic process | 1292 | 727 | 4.83E-06 |
| **GO:0045892** | negative regulation of DNA-templated transcription | 1051 | 601 | 5.22E-06 |
| **GO:0003008** | system process | 1358 | 761 | 5.26E-06 |
| **GO:0006814** | sodium ion transport | 166 | 118 | 5.64E-06 |

**Supplemental Table 5 (continued).**

| GO ID | GO Description | Universe | COSMIC and CLINVAR | Adjusted P-value |
|---|---|---|---|---|
| **GO:0010557** | positive regulation of macromolecule biosynthetic process | 1492 | 830 | 5.80E-06 |
| **GO:0009891** | positive regulation of biosynthetic process | 1593 | 882 | 5.89E-06 |
| **GO:0086001** | cardiac muscle cell action potential | 73 | 60 | 6.04E-06 |
| **GO:0051172** | negative regulation of nitrogen compound metabolic process | 1927 | 1053 | 6.26E-06 |
| **GO:1903507** | negative regulation of nucleic acid-templated transcription | 1056 | 603 | 6.56E-06 |
| **GO:0009890** | negative regulation of biosynthetic process | 1315 | 738 | 6.62E-06 |
| **GO:1902679** | negative regulation of RNA biosynthetic process | 1057 | 603 | 7.97E-06 |
| **GO:0098657** | import into cell | 216 | 147 | 8.46E-06 |
| **GO:0048675** | axon extension | 81 | 65 | 8.67E-06 |
| **GO:0050772** | positive regulation of axonogenesis | 47 | 42 | 9.25E-06 |
| **GO:0060537** | muscle tissue development | 218 | 148 | 9.60E-06 |
| **GO:0050807** | regulation of synapse organization | 124 | 92 | 9.86E-06 |
| **GO:0008283** | cell population proliferation | 1383 | 772 | 9.86E-06 |
| **GO:0001558** | regulation of cell growth | 320 | 206 | 1.11E-05 |
| **GO:0031328** | positive regulation of cellular biosynthetic process | 1568 | 867 | 1.13E-05 |
| **GO:0009790** | embryo development | 505 | 308 | 1.26E-05 |
| **GO:0007166** | cell surface receptor signaling pathway | 2271 | 1225 | 1.38E-05 |
| **GO:0035637** | multicellular organismal signaling | 128 | 94 | 1.60E-05 |
| **GO:0010631** | epithelial cell migration | 261 | 172 | 1.68E-05 |
| **GO:0060070** | canonical Wnt signaling pathway | 254 | 168 | 1.68E-05 |
| **GO:0010632** | regulation of epithelial cell migration | 204 | 139 | 1.92E-05 |
| **GO:0045596** | negative regulation of cell differentiation | 439 | 271 | 2.07E-05 |
| **GO:0050896** | response to stimulus | 7117 | 3620 | 2.09E-05 |
| **GO:0030111** | regulation of Wnt signaling pathway | 274 | 179 | 2.22E-05 |
| **GO:0010558** | negative regulation of macromolecule biosynthetic process | 1251 | 701 | 2.32E-05 |
| **GO:0098739** | import across plasma membrane | 172 | 120 | 2.36E-05 |
| **GO:0034762** | regulation of transmembrane transport | 385 | 241 | 2.39E-05 |
| **GO:0040007** | growth | 515 | 312 | 2.61E-05 |
| **GO:0071310** | cellular response to organic substance | 1734 | 949 | 2.81E-05 |
| **GO:0050803** | regulation of synapse structure or activity | 129 | 94 | 3.02E-05 |
| **GO:0000165** | MAPK cascade | 599 | 357 | 3.11E-05 |
| **GO:0014706** | striated muscle tissue development | 144 | 103 | 3.19E-05 |
| **GO:0035295** | tube development | 616 | 366 | 3.29E-05 |
| **GO:0090132** | epithelium migration | 263 | 172 | 3.75E-05 |
| **GO:0050806** | positive regulation of synaptic transmission | 89 | 69 | 3.90E-05 |
| **GO:0009892** | negative regulation of metabolic process | 2418 | 1295 | 4.08E-05 |
| **GO:0010033** | response to organic substance | 2106 | 1137 | 4.17E-05 |
| **GO:0046777** | protein autophosphorylation | 199 | 135 | 4.63E-05 |
| **GO:0048738** | cardiac muscle tissue development | 138 | 99 | 4.67E-05 |
| **GO:0001505** | regulation of neurotransmitter levels | 143 | 102 | 4.68E-05 |
| **GO:0061337** | cardiac conduction | 91 | 70 | 5.39E-05 |
| **GO:0016358** | dendrite development | 155 | 109 | 5.39E-05 |
| **GO:0090257** | regulation of muscle system process | 157 | 110 | 6.38E-05 |
| **GO:0034765** | regulation of ion transmembrane transport | 325 | 206 | 6.51E-05 |
| **GO:0051253** | negative regulation of RNA metabolic process | 1157 | 649 | 6.64E-05 |
| **GO:0006941** | striated muscle contraction | 142 | 101 | 6.81E-05 |
| **GO:0008361** | regulation of cell size | 119 | 87 | 7.72E-05 |
| **GO:0030155** | regulation of cell adhesion | 614 | 363 | 7.77E-05 |
| **GO:0035239** | tube morphogenesis | 547 | 327 | 7.90E-05 |
| **GO:0090130** | tissue migration | 267 | 173 | 8.81E-05 |
| **GO:0036211** | protein modification process | 2994 | 1581 | 9.04E-05 |
| **GO:0099003** | vesicle-mediated transport in synapse | 121 | 88 | 9.62E-05 |
| **GO:0050905** | neuromuscular process | 73 | 58 | 1.07E-04 |
| **GO:0030048** | actin filament-based movement | 113 | 83 | 1.10E-04 |
| **GO:0042692** | muscle cell differentiation | 243 | 159 | 1.16E-04 |
| **GO:0022607** | cellular component assembly | 2547 | 1355 | 1.30E-04 |
| **GO:0030900** | forebrain development | 155 | 108 | 1.31E-04 |
| **GO:0060078** | regulation of postsynaptic membrane potential | 61 | 50 | 1.38E-04 |
| **GO:0007517** | muscle organ development | 179 | 122 | 1.38E-04 |
| **GO:0009967** | positive regulation of signal transduction | 1255 | 697 | 1.57E-04 |
| **GO:0048639** | positive regulation of developmental growth | 83 | 64 | 1.74E-04 |
| **GO:0048729** | tissue morphogenesis | 348 | 217 | 1.83E-04 |
| **GO:0070887** | cellular response to chemical stimulus | 2288 | 1223 | 1.83E-04 |
| **GO:0043269** | regulation of ion transport | 458 | 277 | 1.97E-04 |

**Supplemental Table 5 (continued).**

| GO ID | GO Description | Universe | COSMIC and CLINVAR | Adjusted P-value |
|---|---|---|---|---|
| **GO:0043269** | regulation of ion transport | 458 | 277 | 1.97E-04 |
| **GO:0035249** | synaptic transmission, glutamatergic | 66 | 53 | 2.05E-04 |
| **GO:0099504** | synaptic vesicle cycle | 114 | 83 | 2.07E-04 |
| **GO:0040008** | regulation of growth | 416 | 254 | 2.14E-04 |
| **GO:0051129** | negative regulation of cellular component organization | 566 | 335 | 2.15E-04 |
| **GO:0021537** | telencephalon development | 101 | 75 | 2.19E-04 |
| **GO:0098900** | regulation of action potential | 48 | 41 | 2.21E-04 |
| **GO:0045934** | negative regulation of nucleobase-containing compound metabolic process | 1273 | 705 | 2.32E-04 |
| **GO:0099565** | chemical synaptic transmission, postsynaptic | 51 | 43 | 2.34E-04 |
| **GO:0051241** | negative regulation of multicellular organismal process | 755 | 435 | 2.53E-04 |
| **GO:0010720** | positive regulation of cell development | 177 | 120 | 2.71E-04 |
| **GO:0140352** | export from cell | 627 | 367 | 2.74E-04 |
| **GO:1902532** | negative regulation of intracellular signal transduction | 436 | 264 | 3.33E-04 |
| **GO:0032409** | regulation of transporter activity | 244 | 158 | 3.37E-04 |
| **GO:0010605** | negative regulation of macromolecule metabolic process | 2244 | 1198 | 3.49E-04 |
| **GO:0008016** | regulation of heart contraction | 164 | 112 | 3.83E-04 |
| **GO:1904062** | regulation of cation transmembrane transport | 291 | 184 | 4.13E-04 |
| **GO:0070727** | cellular macromolecule localization | 1880 | 1013 | 4.24E-04 |
| **GO:0010594** | regulation of endothelial cell migration | 154 | 106 | 4.36E-04 |
| **GO:0043542** | endothelial cell migration | 194 | 129 | 5.00E-04 |
| **GO:0048598** | embryonic morphogenesis | 308 | 193 | 5.37E-04 |
| **GO:0008104** | protein localization | 1874 | 1009 | 5.38E-04 |
| **GO:0044089** | positive regulation of cellular component biogenesis | 416 | 252 | 5.87E-04 |
| **GO:0050790** | regulation of catalytic activity | 1498 | 817 | 6.20E-04 |
| **GO:0198738** | cell-cell signaling by wnt | 343 | 212 | 6.32E-04 |
| **GO:0040017** | positive regulation of locomotion | 472 | 282 | 6.67E-04 |
| **GO:0060048** | cardiac muscle contraction | 111 | 80 | 6.71E-04 |
| **GO:0008154** | actin polymerization or depolymerization | 155 | 106 | 7.08E-04 |
| **GO:1903522** | regulation of blood circulation | 188 | 125 | 7.51E-04 |
| **GO:0030335** | positive regulation of cell migration | 443 | 266 | 7.90E-04 |
| **GO:0051247** | positive regulation of protein metabolic process | 1264 | 696 | 8.30E-04 |
| **GO:0021953** | central nervous system neuron differentiation | 74 | 57 | 8.76E-04 |
| **GO:0030100** | regulation of endocytosis | 159 | 108 | 9.23E-04 |
| **GO:0016055** | Wnt signaling pathway | 339 | 209 | 9.76E-04 |
| **GO:0006836** | neurotransmitter transport | 137 | 95 | 1.04E-03 |
| **GO:0016192** | vesicle-mediated transport | 1326 | 727 | 1.05E-03 |
| **GO:2000147** | positive regulation of cell motility | 463 | 276 | 1.13E-03 |
| **GO:0010243** | response to organonitrogen compound | 613 | 356 | 1.16E-03 |
| **GO:0007163** | establishment or maintenance of cell polarity | 177 | 118 | 1.25E-03 |
| **GO:0030178** | negative regulation of Wnt signaling pathway | 146 | 100 | 1.30E-03 |
| **GO:0099173** | postsynapse organization | 99 | 72 | 1.41E-03 |
| **GO:0006937** | regulation of muscle contraction | 119 | 84 | 1.42E-03 |
| **GO:0007229** | integrin-mediated signaling pathway | 104 | 75 | 1.44E-03 |
| **GO:0070252** | actin-mediated cell contraction | 86 | 64 | 1.50E-03 |
| **GO:0008360** | regulation of cell shape | 116 | 82 | 1.72E-03 |
| **GO:0008015** | blood circulation | 363 | 221 | 1.74E-03 |
| **GO:0060429** | epithelium development | 689 | 395 | 1.79E-03 |
| **GO:0043087** | regulation of GTPase activity | 301 | 187 | 1.79E-03 |
| **GO:0032412** | regulation of ion transmembrane transporter activity | 222 | 143 | 1.84E-03 |
| **GO:0018193** | peptidyl-amino acid modification | 1035 | 575 | 1.97E-03 |
| **GO:0055001** | muscle cell development | 118 | 83 | 2.04E-03 |
| **GO:0070848** | response to growth factor | 516 | 303 | 2.08E-03 |
| **GO:0043085** | positive regulation of catalytic activity | 960 | 536 | 2.08E-03 |
| **GO:0031623** | receptor internalization | 98 | 71 | 2.09E-03 |
| **GO:0051050** | positive regulation of transport | 709 | 405 | 2.15E-03 |
| **GO:0010977** | negative regulation of neuron projection development | 85 | 63 | 2.27E-03 |
| **GO:0031345** | negative regulation of cell projection organization | 125 | 87 | 2.31E-03 |
| **GO:1901699** | cellular response to nitrogen compound | 468 | 277 | 2.40E-03 |
| **GO:0086065** | cell communication involved in cardiac conduction | 58 | 46 | 2.49E-03 |
| **GO:0031532** | actin cytoskeleton reorganization | 105 | 75 | 2.56E-03 |
| **GO:0051146** | striated muscle cell differentiation | 170 | 113 | 2.56E-03 |
| **GO:0051649** | establishment of localization in cell | 1698 | 913 | 2.68E-03 |
| **GO:0046903** | secretion | 638 | 367 | 2.72E-03 |

### Supplemental Table 5 (continued).

| GO ID | GO Description | Universe | COSMIC and CLINVAR | Adjusted P-value |
|---|---|---|---|---|
| **GO:0046578** | regulation of Ras protein signal transduction | 139 | 95 | 2.74E-03 |
| **GO:0031399** | regulation of protein modification process | 1259 | 689 | 2.83E-03 |
| **GO:0071417** | cellular response to organonitrogen compound | 413 | 247 | 2.92E-03 |
| **GO:0048813** | dendrite morphogenesis | 97 | 70 | 3.06E-03 |
| **GO:0060291** | long-term synaptic potentiation | 49 | 40 | 3.09E-03 |
| **GO:0071526** | semaphorin-plexin signaling pathway | 40 | 34 | 3.13E-03 |
| **GO:0050771** | negative regulation of axonogenesis | 43 | 36 | 3.22E-03 |
| **GO:0022898** | regulation of transmembrane transporter activity | 229 | 146 | 3.28E-03 |
| **GO:0032940** | secretion by cell | 571 | 331 | 3.39E-03 |
| **GO:0051174** | regulation of phosphorus metabolic process | 1126 | 620 | 3.43E-03 |
| **GO:0042325** | regulation of phosphorylation | 1004 | 557 | 3.54E-03 |
| **GO:0098901** | regulation of cardiac muscle cell action potential | 30 | 27 | 3.62E-03 |
| **GO:0040013** | negative regulation of locomotion | 278 | 173 | 3.69E-03 |
| **GO:0007416** | synapse assembly | 126 | 87 | 3.77E-03 |
| **GO:0019220** | regulation of phosphate metabolic process | 1125 | 619 | 3.93E-03 |
| **GO:0043408** | regulation of MAPK cascade | 534 | 311 | 3.94E-03 |
| **GO:0045229** | external encapsulating structure organization | 237 | 150 | 4.35E-03 |
| **GO:0051966** | regulation of synaptic transmission, glutamatergic | 51 | 41 | 4.76E-03 |
| **GO:0006898** | receptor-mediated endocytosis | 214 | 137 | 4.77E-03 |
| **GO:0086003** | cardiac muscle cell contraction | 62 | 48 | 5.00E-03 |
| **GO:0098703** | calcium ion import across plasma membrane | 36 | 31 | 5.04E-03 |
| **GO:0086002** | cardiac muscle cell action potential involved in contraction | 48 | 39 | 5.09E-03 |
| **GO:0033036** | macromolecule localization | 2245 | 1186 | 5.15E-03 |
| **GO:0001944** | vasculature development | 492 | 288 | 5.17E-03 |
| **GO:0050890** | cognition | 156 | 104 | 5.19E-03 |
| **GO:0030198** | extracellular matrix organization | 234 | 148 | 5.31E-03 |
| **GO:0060079** | excitatory postsynaptic potential | 45 | 37 | 5.32E-03 |
| **GO:1901888** | regulation of cell junction assembly | 144 | 97 | 5.54E-03 |
| **GO:0050678** | regulation of epithelial cell proliferation | 236 | 149 | 5.67E-03 |
| **GO:1902903** | regulation of supramolecular fiber organization | 311 | 190 | 6.64E-03 |
| **GO:0043062** | extracellular structure organization | 235 | 148 | 7.38E-03 |
| **GO:0002009** | morphogenesis of an epithelium | 275 | 170 | 7.74E-03 |
| **GO:0042592** | homeostatic process | 1249 | 680 | 7.85E-03 |
| **GO:0043412** | macromolecule modification | 3198 | 1659 | 8.65E-03 |
| **GO:0086091** | regulation of heart rate by cardiac conduction | 41 | 34 | 9.11E-03 |
| **GO:0110053** | regulation of actin filament organization | 216 | 137 | 9.48E-03 |
| **GO:0031400** | negative regulation of protein modification process | 409 | 242 | 1.05E-02 |
| **GO:0001568** | blood vessel development | 469 | 274 | 1.06E-02 |
| **GO:0090090** | negative regulation of canonical Wnt signaling pathway | 118 | 81 | 1.13E-02 |
| **GO:0048592** | eye morphogenesis | 81 | 59 | 1.14E-02 |
| **GO:0051246** | regulation of protein metabolic process | 2072 | 1095 | 1.17E-02 |
| **GO:0007423** | sensory organ development | 271 | 167 | 1.19E-02 |
| **GO:2001257** | regulation of cation channel activity | 153 | 101 | 1.31E-02 |
| **GO:0070588** | calcium ion transmembrane transport | 235 | 147 | 1.31E-02 |
| **GO:0007611** | learning or memory | 115 | 79 | 1.37E-02 |
| **GO:0048863** | stem cell differentiation | 155 | 102 | 1.44E-02 |
| **GO:0031401** | positive regulation of protein modification process | 814 | 454 | 1.44E-02 |
| **GO:0060341** | regulation of cellular localization | 766 | 429 | 1.48E-02 |
| **GO:0021954** | central nervous system neuron development | 40 | 33 | 1.51E-02 |
| **GO:0071363** | cellular response to growth factor stimulus | 499 | 289 | 1.51E-02 |
| **GO:1901379** | regulation of potassium ion transmembrane transport | 70 | 52 | 1.53E-02 |
| **GO:0030041** | actin filament polymerization | 129 | 87 | 1.53E-02 |
| **GO:0048514** | blood vessel morphogenesis | 439 | 257 | 1.60E-02 |
| **GO:0050773** | regulation of dendrite development | 62 | 47 | 1.62E-02 |
| **GO:0009792** | embryo development ending in birth or egg hatching | 198 | 126 | 1.68E-02 |
| **GO:0051961** | negative regulation of nervous system development | 85 | 61 | 1.69E-02 |
| **GO:0090596** | sensory organ morphogenesis | 119 | 81 | 1.79E-02 |
| **GO:0001501** | skeletal system development | 289 | 176 | 1.82E-02 |
| **GO:0120031** | plasma membrane bounded cell projection assembly | 483 | 280 | 1.85E-02 |
| **GO:0006816** | calcium ion transport | 313 | 189 | 1.85E-02 |
| **GO:0010634** | positive regulation of epithelial cell migration | 133 | 89 | 1.92E-02 |
| **GO:0031098** | stress-activated protein kinase signaling cascade | 202 | 128 | 1.93E-02 |
| **GO:0009628** | response to abiotic stimulus | 712 | 400 | 1.97E-02 |
| **GO:1990573** | potassium ion import across plasma membrane | 48 | 38 | 1.99E-02 |

**Supplemental Table 5 (continued).**

| GO ID | GO Description | Universe | COSMIC and CLINVAR | Adjusted P-value |
|---|---|---|---|---|
| **GO:0150104** | transport across blood-brain barrier | 87 | 62 | 2.03E-02 |
| **GO:0010232** | vascular transport | 87 | 62 | 2.03E-02 |
| **GO:0007160** | cell-matrix adhesion | 195 | 124 | 2.04E-02 |
| **GO:0032880** | regulation of protein localization | 672 | 379 | 2.06E-02 |
| **GO:0043254** | regulation of protein-containing complex assembly | 332 | 199 | 2.11E-02 |
| **GO:0051258** | protein polymerization | 233 | 145 | 2.14E-02 |
| **GO:0000122** | negative regulation of transcription by RNA polymerase II | 751 | 420 | 2.14E-02 |
| **GO:0007612** | learning | 53 | 41 | 2.48E-02 |
| **GO:0045664** | regulation of neuron differentiation | 96 | 67 | 2.77E-02 |
| **GO:0010959** | regulation of metal ion transport | 298 | 180 | 2.80E-02 |
| **GO:0030010** | establishment of cell polarity | 108 | 74 | 2.86E-02 |
| **GO:0007269** | neurotransmitter secretion | 91 | 64 | 2.87E-02 |
| **GO:0099643** | signal release from synapse | 91 | 64 | 2.87E-02 |
| **GO:0051963** | regulation of synapse assembly | 58 | 44 | 2.89E-02 |
| **GO:0016079** | synaptic vesicle exocytosis | 58 | 44 | 2.89E-02 |
| **GO:0150063** | visual system development | 207 | 130 | 3.03E-02 |
| **GO:0051403** | stress-activated MAPK cascade | 198 | 125 | 3.03E-02 |
| **GO:0061572** | actin filament bundle organization | 136 | 90 | 3.22E-02 |
| **GO:0048762** | mesenchymal cell differentiation | 191 | 121 | 3.23E-02 |
| **GO:0007215** | glutamate receptor signaling pathway | 44 | 35 | 3.51E-02 |
| **GO:0086009** | membrane repolarization | 44 | 35 | 3.51E-02 |
| **GO:0048880** | sensory system development | 213 | 133 | 3.64E-02 |
| **GO:0032878** | regulation of establishment or maintenance of cell polarity | 26 | 23 | 3.65E-02 |
| **GO:0001654** | eye development | 204 | 128 | 3.68E-02 |
| **GO:0048640** | negative regulation of developmental growth | 60 | 45 | 3.78E-02 |
| **GO:0030031** | cell projection assembly | 498 | 286 | 3.80E-02 |
| **GO:0043549** | regulation of kinase activity | 599 | 339 | 3.81E-02 |
| **GO:0050881** | musculoskeletal movement | 41 | 33 | 3.82E-02 |
| **GO:0050879** | multicellular organismal movement | 41 | 33 | 3.82E-02 |
| **GO:1990778** | protein localization to cell periphery | 279 | 169 | 3.89E-02 |
| **GO:0051017** | actin filament bundle assembly | 133 | 88 | 3.94E-02 |
| **GO:0086005** | ventricular cardiac muscle cell action potential | 29 | 25 | 4.10E-02 |
| **GO:0050919** | negative chemotaxis | 35 | 29 | 4.26E-02 |
| **GO:0030517** | negative regulation of axon extension | 32 | 27 | 4.29E-02 |
| **GO:0048878** | chemical homeostasis | 769 | 427 | 4.30E-02 |
| **GO:0042327** | positive regulation of phosphorylation | 663 | 372 | 4.31E-02 |
| **GO:0050768** | negative regulation of neurogenesis | 80 | 57 | 4.32E-02 |
| **GO:0031323** | regulation of cellular metabolic process | 4966 | 2520 | 4.38E-02 |
| **GO:0043009** | chordate embryonic development | 183 | 116 | 4.45E-02 |
| **GO:0010721** | negative regulation of cell development | 109 | 74 | 4.53E-02 |
| **GO:0045785** | positive regulation of cell adhesion | 369 | 217 | 4.75E-02 |
| **GO:0006887** | exocytosis | 267 | 162 | 4.91E-02 |
| **GO:0045216** | cell-cell junction organization | 169 | 108 | 4.96E-02 |
| **GO:0001764** | neuron migration | 87 | 61 | 4.98E-02 |

**Supplemental Table 6.** Significant GO:BP enrichments for all COSMIC G4 mutations.

| GO ID | GO Description | Universe | COSMIC | Adjusted |
|---|---|---|---|---|
| **GO:0007399** | nervous system development | 1648 | 1074 | 3.73E-48 |
| **GO:0048856** | anatomical structure development | 4152 | 2391 | 3.71E-47 |
| **GO:0032502** | developmental process | 4584 | 2606 | 2.53E-46 |
| **GO:0048731** | system development | 2976 | 1762 | 2.21E-41 |
| **GO:0007275** | multicellular organism development | 3266 | 1908 | 2.56E-40 |
| **GO:0048699** | generation of neurons | 969 | 663 | 7.68E-38 |
| **GO:0009653** | anatomical structure morphogenesis | 1871 | 1162 | 2.2E-37 |
| **GO:0000902** | cell morphogenesis | 718 | 515 | 2.79E-37 |
| **GO:0030154** | cell differentiation | 2860 | 1682 | 1.33E-36 |
| **GO:0048869** | cellular developmental process | 2879 | 1691 | 2.15E-36 |
| **GO:0022008** | neurogenesis | 1096 | 728 | 4.84E-35 |
| **GO:0030182** | neuron differentiation | 923 | 629 | 7.45E-35 |
| **GO:0048858** | cell projection morphogenesis | 459 | 349 | 9.15E-33 |
| **GO:0032989** | cellular component morphogenesis | 549 | 404 | 1.13E-32 |
| **GO:0048468** | cell development | 1355 | 865 | 1.38E-32 |
| **GO:0032501** | multicellular organismal process | 5319 | 2904 | 1.87E-32 |
| **GO:0030030** | cell projection organization | 1164 | 758 | 3.34E-32 |
| **GO:0048666** | neuron development | 729 | 510 | 3.54E-32 |
| **GO:0120039** | plasma membrane bounded cell projection morphogenesis | 455 | 345 | 6.29E-32 |
| **GO:0120036** | plasma membrane bounded cell projection organization | 1144 | 746 | 6.38E-32 |
| **GO:0031175** | neuron projection development | 656 | 466 | 1.03E-31 |
| **GO:0032990** | cell part morphogenesis | 469 | 352 | 4.87E-31 |
| **GO:0048812** | neuron projection morphogenesis | 441 | 334 | 1.2E-30 |
| **GO:0023051** | regulation of signaling | 2733 | 1582 | 7.93E-29 |
| **GO:0010646** | regulation of cell communication | 2727 | 1578 | 1.24E-28 |
| **GO:0000904** | cell morphogenesis involved in differentiation | 495 | 363 | 1.5E-28 |
| **GO:0051128** | regulation of cellular component organization | 1929 | 1154 | 5.21E-27 |
| **GO:0061564** | axon development | 323 | 253 | 1.39E-26 |
| **GO:0030029** | actin filament-based process | 717 | 489 | 1.87E-26 |
| **GO:0048667** | cell morphogenesis involved in neuron differentiation | 381 | 288 | 8.11E-26 |
| **GO:0007409** | axonogenesis | 298 | 235 | 2.6E-25 |
| **GO:0050793** | regulation of developmental process | 1810 | 1084 | 2.62E-25 |
| **GO:0023052** | signaling | 5190 | 2793 | 2.45E-24 |
| **GO:0007154** | cell communication | 5221 | 2801 | 3.47E-23 |
| **GO:0016043** | cellular component organization | 5370 | 2874 | 3.66E-23 |
| **GO:0030036** | actin cytoskeleton organization | 637 | 432 | 3E-22 |
| **GO:0051239** | regulation of multicellular organismal process | 2117 | 1227 | 3.02E-21 |
| **GO:0048513** | animal organ development | 2176 | 1257 | 4.1E-21 |
| **GO:0007155** | cell adhesion | 1216 | 749 | 6.57E-21 |
| **GO:0035556** | intracellular signal transduction | 2168 | 1250 | 1.57E-20 |
| **GO:0009966** | regulation of signal transduction | 2464 | 1398 | 1.28E-19 |
| **GO:0007010** | cytoskeleton organization | 1309 | 791 | 7.32E-19 |
| **GO:0050804** | modulation of chemical synaptic transmission | 252 | 194 | 3.06E-18 |
| **GO:0071840** | cellular component organization or biogenesis | 5539 | 2923 | 3.12E-18 |
| **GO:0099177** | regulation of trans-synaptic signaling | 253 | 194 | 6.91E-18 |
| **GO:0048522** | positive regulation of cellular process | 4704 | 2510 | 9.93E-18 |
| **GO:0031344** | regulation of cell projection organization | 467 | 322 | 1.09E-17 |
| **GO:0099536** | synaptic signaling | 522 | 353 | 2.18E-17 |
| **GO:0120035** | regulation of plasma membrane bounded cell projection organization | 453 | 313 | 2.39E-17 |
| **GO:0034330** | cell junction organization | 518 | 349 | 9.31E-17 |
| **GO:0048523** | negative regulation of cellular process | 3896 | 2101 | 2.01E-16 |
| **GO:0048518** | positive regulation of biological process | 5294 | 2788 | 3.67E-16 |
| **GO:0050794** | regulation of cellular process | 9523 | 4812 | 4.06E-16 |
| **GO:0045595** | regulation of cell differentiation | 1129 | 683 | 5.21E-16 |
| **GO:0010975** | regulation of neuron projection development | 288 | 212 | 5.22E-16 |
| **GO:0099537** | trans-synaptic signaling | 501 | 337 | 6.23E-16 |
| **GO:0098916** | anterograde trans-synaptic signaling | 495 | 333 | 9.95E-16 |
| **GO:0007268** | chemical synaptic transmission | 495 | 333 | 9.95E-16 |
| **GO:0048583** | regulation of response to stimulus | 3327 | 1812 | 1.08E-15 |
| **GO:0007417** | central nervous system development | 584 | 383 | 1.53E-15 |
| **GO:0065007** | biological regulation | 10721 | 5366 | 1.62E-15 |
| **GO:0009987** | cellular process | 14783 | 7189 | 2.34E-15 |
| **GO:0022603** | regulation of anatomical structure morphogenesis | 700 | 447 | 2.71E-15 |
| **GO:0007165** | signal transduction | 4776 | 2527 | 3.25E-15 |
| **GO:0007267** | cell-cell signaling | 1269 | 754 | 3.55E-15 |
| **GO:0097485** | neuron projection guidance | 169 | 136 | 3.85E-15 |

**Supplemental Table 6 (continued).**

| GO ID | GO Description | Universe | COSMIC | Adjusted |
|---|---|---|---|---|
| **GO:0007411** | axon guidance | 169 | 136 | 3.85E-15 |
| **GO:0051716** | cellular response to stimulus | 5982 | 3114 | 4.88E-15 |
| **GO:0050789** | regulation of biological process | 10085 | 5066 | 5.3E-15 |
| **GO:0032879** | regulation of localization | 1615 | 935 | 5.33E-15 |
| **GO:0065008** | regulation of biological quality | 2937 | 1610 | 8.9E-15 |
| **GO:0048519** | negative regulation of biological process | 4381 | 2326 | 2.42E-14 |
| **GO:0050770** | regulation of axonogenesis | 100 | 88 | 7.4E-14 |
| **GO:0051094** | positive regulation of developmental process | 967 | 587 | 8.44E-14 |
| **GO:0051049** | regulation of transport | 1327 | 777 | 1.58E-13 |
| **GO:0051179** | localization | 4343 | 2298 | 3.85E-13 |
| **GO:0050808** | synapse organization | 275 | 197 | 1.2E-12 |
| **GO:0051960** | regulation of nervous system development | 257 | 186 | 1.51E-12 |
| **GO:0048870** | cell motility | 1362 | 790 | 1.94E-12 |
| **GO:0007167** | enzyme-linked receptor protein signaling pathway | 795 | 487 | 5.4E-12 |
| **GO:0065009** | regulation of molecular function | 2121 | 1178 | 7.37E-12 |
| **GO:0009887** | animal organ morphogenesis | 582 | 370 | 7.74E-12 |
| **GO:0098660** | inorganic ion transmembrane transport | 622 | 392 | 8.1E-12 |
| **GO:0007015** | actin filament organization | 395 | 264 | 1.69E-11 |
| **GO:0003012** | muscle system process | 305 | 212 | 1.71E-11 |
| **GO:0051130** | positive regulation of cellular component organization | 844 | 511 | 2.1E-11 |
| **GO:0040011** | locomotion | 1075 | 634 | 2.15E-11 |
| **GO:0016477** | cell migration | 1210 | 705 | 2.25E-11 |
| **GO:0098655** | cation transmembrane transport | 656 | 408 | 4.28E-11 |
| **GO:0006810** | transport | 3620 | 1924 | 5E-11 |
| **GO:0055085** | transmembrane transport | 1060 | 624 | 5.85E-11 |
| **GO:0009888** | tissue development | 1239 | 718 | 6.11E-11 |
| **GO:0061061** | muscle structure development | 407 | 269 | 7.1E-11 |
| **GO:0016310** | phosphorylation | 1444 | 824 | 8.45E-11 |
| **GO:0034220** | ion transmembrane transport | 816 | 493 | 1.05E-10 |
| **GO:0042391** | regulation of membrane potential | 319 | 218 | 1.17E-10 |
| **GO:0098662** | inorganic cation transmembrane transport | 572 | 360 | 1.47E-10 |
| **GO:0050767** | regulation of neurogenesis | 208 | 152 | 1.8E-10 |
| **GO:1902531** | regulation of intracellular signal transduction | 1429 | 814 | 2.13E-10 |
| **GO:0045944** | positive regulation of transcription by RNA polymerase II | 952 | 564 | 2.35E-10 |
| **GO:0051234** | establishment of localization | 3775 | 1995 | 2.45E-10 |
| **GO:0007420** | brain development | 384 | 254 | 3.26E-10 |
| **GO:0030001** | metal ion transport | 678 | 416 | 4.56E-10 |
| **GO:0040012** | regulation of locomotion | 842 | 504 | 4.98E-10 |
| **GO:0097435** | supramolecular fiber organization | 710 | 433 | 5.41E-10 |
| **GO:0007264** | small GTPase mediated signal transduction | 389 | 256 | 6.15E-10 |
| **GO:0007169** | transmembrane receptor protein tyrosine kinase signaling pathway | 511 | 324 | 6.85E-10 |
| **GO:0060322** | head development | 402 | 263 | 7.88E-10 |
| **GO:0023057** | negative regulation of signaling | 1067 | 622 | 8.51E-10 |
| **GO:0010648** | negative regulation of cell communication | 1060 | 618 | 9.87E-10 |
| **GO:0051962** | positive regulation of nervous system development | 147 | 113 | 1.03E-09 |
| **GO:0045597** | positive regulation of cell differentiation | 624 | 385 | 1.21E-09 |
| **GO:0006811** | ion transport | 1187 | 684 | 1.24E-09 |
| **GO:0048167** | regulation of synaptic plasticity | 122 | 97 | 1.24E-09 |
| **GO:0006936** | muscle contraction | 260 | 181 | 1.25E-09 |
| **GO:0051173** | positive regulation of nitrogen compound metabolic process | 2565 | 1387 | 1.3E-09 |
| **GO:0030334** | regulation of cell migration | 767 | 462 | 1.33E-09 |
| **GO:0006468** | protein phosphorylation | 1268 | 726 | 1.34E-09 |
| **GO:2000145** | regulation of cell motility | 818 | 489 | 1.52E-09 |
| **GO:0022604** | regulation of cell morphogenesis | 240 | 169 | 1.56E-09 |
| **GO:0071805** | potassium ion transmembrane transport | 185 | 136 | 1.7E-09 |
| **GO:2000026** | regulation of multicellular organismal development | 981 | 575 | 1.88E-09 |
| **GO:0006812** | cation transport | 886 | 524 | 2.72E-09 |
| **GO:1903508** | positive regulation of nucleic acid-templated transcription | 1297 | 739 | 3.11E-09 |
| **GO:0045893** | positive regulation of DNA-templated transcription | 1297 | 739 | 3.11E-09 |
| **GO:0031346** | positive regulation of cell projection organization | 247 | 172 | 4.48E-09 |
| **GO:0048585** | negative regulation of response to stimulus | 1287 | 732 | 6.54E-09 |
| **GO:0031325** | positive regulation of cellular metabolic process | 2522 | 1360 | 6.74E-09 |
| **GO:0060560** | developmental growth involved in morphogenesis | 135 | 104 | 6.79E-09 |
| **GO:1902680** | positive regulation of RNA biosynthetic process | 1303 | 740 | 7.32E-09 |
| **GO:0048588** | developmental cell growth | 129 | 100 | 9.07E-09 |
| **GO:0032970** | regulation of actin filament-based process | 330 | 219 | 9.35E-09 |

**Supplemental Table 6 (continued).**

| GO ID | GO Description | Universe | COSMIC | Adjusted |
|---|---|---|---|---|
| **GO:0009968** | negative regulation of signal transduction | 1009 | 586 | 1.04E-08 |
| **GO:0098609** | cell-cell adhesion | 743 | 445 | 1.17E-08 |
| **GO:0032535** | regulation of cellular component size | 256 | 176 | 1.4E-08 |
| **GO:0044087** | regulation of cellular component biogenesis | 774 | 461 | 1.54E-08 |
| **GO:0044057** | regulation of system process | 392 | 253 | 1.88E-08 |
| **GO:0051056** | regulation of small GTPase mediated signal transduction | 238 | 165 | 2.2E-08 |
| **GO:0098659** | inorganic cation import across plasma membrane | 112 | 88 | 4.74E-08 |
| **GO:0099587** | inorganic ion import across plasma membrane | 112 | 88 | 4.74E-08 |
| **GO:0031589** | cell-substrate adhesion | 290 | 194 | 5.36E-08 |
| **GO:0060284** | regulation of cell development | 315 | 208 | 6.43E-08 |
| **GO:0072359** | circulatory system development | 729 | 434 | 7.08E-08 |
| **GO:0050769** | positive regulation of neurogenesis | 119 | 92 | 8.43E-08 |
| **GO:0006813** | potassium ion transport | 200 | 141 | 1.05E-07 |
| **GO:0031324** | negative regulation of cellular metabolic process | 1858 | 1016 | 1.24E-07 |
| **GO:0060627** | regulation of vesicle-mediated transport | 417 | 264 | 1.34E-07 |
| **GO:0051254** | positive regulation of RNA metabolic process | 1433 | 799 | 1.48E-07 |
| **GO:0044093** | positive regulation of molecular function | 1305 | 733 | 1.63E-07 |
| **GO:0051240** | positive regulation of multicellular organismal process | 1137 | 646 | 1.77E-07 |
| **GO:1990138** | neuron projection extension | 112 | 87 | 1.81E-07 |
| **GO:0009893** | positive regulation of metabolic process | 3177 | 1674 | 1.93E-07 |
| **GO:0010604** | positive regulation of macromolecule metabolic process | 2889 | 1531 | 2.03E-07 |
| **GO:1905114** | cell surface receptor signaling pathway involved in cell-cell signaling | 388 | 247 | 2.48E-07 |
| **GO:0045935** | positive regulation of nucleobase-containing compound metabolic process | 1620 | 893 | 2.49E-07 |
| **GO:0042221** | response to chemical | 2943 | 1557 | 2.51E-07 |
| **GO:0035725** | sodium ion transmembrane transport | 129 | 97 | 3.85E-07 |
| **GO:0023056** | positive regulation of signaling | 1380 | 769 | 4.04E-07 |
| **GO:0030516** | regulation of axon extension | 61 | 53 | 4.35E-07 |
| **GO:0034329** | cell junction assembly | 341 | 220 | 4.37E-07 |
| **GO:0006793** | phosphorus metabolic process | 2354 | 1261 | 4.9E-07 |
| **GO:0051641** | cellular localization | 2721 | 1444 | 5.04E-07 |
| **GO:0033043** | regulation of organelle organization | 1015 | 580 | 5.1E-07 |
| **GO:0061387** | regulation of extent of cell growth | 67 | 57 | 5.2E-07 |
| **GO:0010647** | positive regulation of cell communication | 1374 | 765 | 5.58E-07 |
| **GO:0016049** | cell growth | 358 | 229 | 6.06E-07 |
| **GO:0032956** | regulation of actin cytoskeleton organization | 296 | 194 | 7.37E-07 |
| **GO:0006796** | phosphate-containing compound metabolic process | 2336 | 1250 | 8.5E-07 |
| **GO:0060828** | regulation of canonical Wnt signaling pathway | 211 | 145 | 9.9E-07 |
| **GO:0045892** | negative regulation of DNA-templated transcription | 1051 | 597 | 1E-06 |
| **GO:0051093** | negative regulation of developmental process | 619 | 370 | 1.02E-06 |
| **GO:0048638** | regulation of developmental growth | 167 | 119 | 1.13E-06 |
| **GO:0006996** | organelle organization | 3147 | 1652 | 1.13E-06 |
| **GO:0001667** | ameboidal-type cell migration | 333 | 214 | 1.26E-06 |
| **GO:0048589** | developmental growth | 273 | 180 | 1.54E-06 |
| **GO:1903507** | negative regulation of nucleic acid-templated transcription | 1056 | 598 | 1.85E-06 |
| **GO:0031327** | negative regulation of cellular biosynthetic process | 1292 | 719 | 2.24E-06 |
| **GO:1902679** | negative regulation of RNA biosynthetic process | 1057 | 598 | 2.26E-06 |
| **GO:0042127** | regulation of cell population proliferation | 1218 | 681 | 2.26E-06 |
| **GO:0051172** | negative regulation of nitrogen compound metabolic process | 1927 | 1041 | 2.45E-06 |
| **GO:0090066** | regulation of anatomical structure size | 328 | 210 | 2.86E-06 |
| **GO:0009890** | negative regulation of biosynthetic process | 1315 | 730 | 2.92E-06 |
| **GO:0051493** | regulation of cytoskeleton organization | 450 | 277 | 2.94E-06 |
| **GO:0003015** | heart process | 193 | 133 | 3.77E-06 |
| **GO:0048675** | axon extension | 81 | 65 | 3.8E-06 |
| **GO:0060070** | canonical Wnt signaling pathway | 254 | 168 | 3.9E-06 |
| **GO:0071495** | cellular response to endogenous stimulus | 974 | 553 | 4.87E-06 |
| **GO:0050772** | positive regulation of axonogenesis | 47 | 42 | 5.03E-06 |
| **GO:0003013** | circulatory system process | 443 | 272 | 5.81E-06 |
| **GO:0001508** | action potential | 117 | 87 | 8.02E-06 |
| **GO:0001558** | regulation of cell growth | 320 | 204 | 8.08E-06 |
| **GO:0010557** | positive regulation of macromolecule biosynthetic process | 1492 | 817 | 8.38E-06 |
| **GO:0010558** | negative regulation of macromolecule biosynthetic process | 1251 | 694 | 8.82E-06 |
| **GO:0009891** | positive regulation of biosynthetic process | 1593 | 868 | 8.93E-06 |
| **GO:0009719** | response to endogenous stimulus | 1076 | 604 | 9.26E-06 |
| **GO:0030111** | regulation of Wnt signaling pathway | 274 | 178 | 1E-05 |
| **GO:0050807** | regulation of synapse organization | 124 | 91 | 1.07E-05 |
| **GO:0050896** | response to stimulus | 7117 | 3567 | 1.15E-05 |

### Supplemental Table 6 (continued).

| GO ID | GO Description | Universe | COSMIC | Adjusted |
|---|---|---|---|---|
| **GO:0007265** | Ras protein signal transduction | 278 | 180 | 1.19E-05 |
| **GO:0036211** | protein modification process | 2994 | 1566 | 1.32E-05 |
| **GO:0007166** | cell surface receptor signaling pathway | 2271 | 1207 | 1.39E-05 |
| **GO:0060047** | heart contraction | 187 | 128 | 1.41E-05 |
| **GO:0007507** | heart development | 338 | 213 | 1.5E-05 |
| **GO:0045596** | negative regulation of cell differentiation | 439 | 268 | 1.62E-05 |
| **GO:0051253** | negative regulation of RNA metabolic process | 1157 | 644 | 1.66E-05 |
| **GO:0031328** | positive regulation of cellular biosynthetic process | 1568 | 853 | 1.82E-05 |
| **GO:0008283** | cell population proliferation | 1383 | 759 | 1.9E-05 |
| **GO:0003008** | system process | 1358 | 746 | 2.09E-05 |
| **GO:0000165** | MAPK cascade | 599 | 353 | 2.24E-05 |
| **GO:0048646** | anatomical structure formation involved in morphogenesis | 751 | 433 | 2.29E-05 |
| **GO:0071310** | cellular response to organic substance | 1734 | 936 | 2.3E-05 |
| **GO:0034762** | regulation of transmembrane transport | 385 | 238 | 2.38E-05 |
| **GO:0098657** | import into cell | 216 | 144 | 2.44E-05 |
| **GO:0040007** | growth | 515 | 308 | 2.56E-05 |
| **GO:0006814** | sodium ion transport | 166 | 115 | 2.78E-05 |
| **GO:0008361** | regulation of cell size | 119 | 87 | 3.03E-05 |
| **GO:0050803** | regulation of synapse structure or activity | 129 | 93 | 3.1E-05 |
| **GO:0030155** | regulation of cell adhesion | 614 | 360 | 3.42E-05 |
| **GO:0009790** | embryo development | 505 | 302 | 3.52E-05 |
| **GO:0098739** | import across plasma membrane | 172 | 118 | 4.28E-05 |
| **GO:0001505** | regulation of neurotransmitter levels | 143 | 101 | 4.33E-05 |
| **GO:0010033** | response to organic substance | 2106 | 1120 | 4.51E-05 |
| **GO:0034765** | regulation of ion transmembrane transport | 325 | 204 | 4.69E-05 |
| **GO:0009892** | negative regulation of metabolic process | 2418 | 1275 | 4.96E-05 |
| **GO:0090257** | regulation of muscle system process | 157 | 109 | 5.34E-05 |
| **GO:0060537** | muscle tissue development | 218 | 144 | 5.78E-05 |
| **GO:0050806** | positive regulation of synaptic transmission | 89 | 68 | 5.99E-05 |
| **GO:0046777** | protein autophosphorylation | 199 | 133 | 6.67E-05 |
| **GO:0060078** | regulation of postsynaptic membrane potential | 61 | 50 | 7.28E-05 |
| **GO:0009967** | positive regulation of signal transduction | 1255 | 689 | 8.73E-05 |
| **GO:0045934** | negative regulation of nucleobase-containing compound metabolic process | 1273 | 698 | 9.18E-05 |
| **GO:0022607** | cellular component assembly | 2547 | 1336 | 0.000101 |
| **GO:0007517** | muscle organ development | 179 | 121 | 0.000103 |
| **GO:0099003** | vesicle-mediated transport in synapse | 121 | 87 | 0.000106 |
| **GO:0035249** | synaptic transmission, glutamatergic | 66 | 53 | 0.000106 |
| **GO:0070887** | cellular response to chemical stimulus | 2288 | 1207 | 0.000112 |
| **GO:0016358** | dendrite development | 155 | 107 | 0.000112 |
| **GO:0030900** | forebrain development | 155 | 107 | 0.000112 |
| **GO:0035637** | multicellular organismal signaling | 128 | 91 | 0.000126 |
| **GO:0042692** | muscle cell differentiation | 243 | 157 | 0.000127 |
| **GO:0010632** | regulation of epithelial cell migration | 204 | 135 | 0.000129 |
| **GO:0010631** | epithelial cell migration | 261 | 167 | 0.000131 |
| **GO:0035295** | tube development | 616 | 358 | 0.000132 |
| **GO:0099565** | chemical synaptic transmission, postsynaptic | 51 | 43 | 0.000132 |
| **GO:0070727** | cellular macromolecule localization | 1880 | 1002 | 0.000162 |
| **GO:0140352** | export from cell | 627 | 363 | 0.000185 |
| **GO:0050905** | neuromuscular process | 73 | 57 | 0.000203 |
| **GO:0086001** | cardiac muscle cell action potential | 73 | 57 | 0.000203 |
| **GO:0010720** | positive regulation of cell development | 177 | 119 | 0.000204 |
| **GO:0008104** | protein localization | 1874 | 998 | 0.000208 |
| **GO:0050790** | regulation of catalytic activity | 1498 | 809 | 0.000218 |
| **GO:0099504** | synaptic vesicle cycle | 114 | 82 | 0.000241 |
| **GO:0043269** | regulation of ion transport | 458 | 273 | 0.000241 |
| **GO:0198738** | cell-cell signaling by wnt | 343 | 211 | 0.000251 |
| **GO:0051247** | positive regulation of protein metabolic process | 1264 | 690 | 0.00026 |
| **GO:0061337** | cardiac conduction | 91 | 68 | 0.00026 |
| **GO:0090132** | epithelium migration | 263 | 167 | 0.000269 |
| **GO:0035239** | tube morphogenesis | 547 | 320 | 0.00027 |
| **GO:0048639** | positive regulation of developmental growth | 83 | 63 | 0.000281 |
| **GO:0021537** | telencephalon development | 101 | 74 | 0.000288 |
| **GO:0040008** | regulation of growth | 416 | 250 | 0.000312 |
| **GO:1904062** | regulation of cation transmembrane transport | 291 | 182 | 0.000353 |
| **GO:0032409** | regulation of transporter activity | 244 | 156 | 0.000361 |
| **GO:0010605** | negative regulation of macromolecule metabolic process | 2244 | 1180 | 0.000368 |

### Supplemental Table 6 (continued).

| GO ID | GO Description | Universe | COSMIC | Adjusted |
|---|---|---|---|---|
| **GO:0048729** | tissue morphogenesis | 348 | 213 | 0.000371 |
| **GO:0006941** | striated muscle contraction | 142 | 98 | 0.000389 |
| **GO:0016055** | Wnt signaling pathway | 339 | 208 | 0.000396 |
| **GO:1902532** | negative regulation of intracellular signal transduction | 436 | 260 | 0.000441 |
| **GO:0014706** | striated muscle tissue development | 144 | 99 | 0.000447 |
| **GO:0040017** | positive regulation of locomotion | 472 | 279 | 0.000478 |
| **GO:0044089** | positive regulation of cellular component biogenesis | 416 | 249 | 0.000511 |
| **GO:0031399** | regulation of protein modification process | 1259 | 685 | 0.000532 |
| **GO:0090130** | tissue migration | 267 | 168 | 0.000573 |
| **GO:0008154** | actin polymerization or depolymerization | 155 | 105 | 0.000608 |
| **GO:0030335** | positive regulation of cell migration | 443 | 263 | 0.000628 |
| **GO:0051129** | negative regulation of cellular component organization | 566 | 328 | 0.000642 |
| **GO:0098900** | regulation of action potential | 48 | 40 | 0.000679 |
| **GO:0016192** | vesicle-mediated transport | 1326 | 718 | 0.000692 |
| **GO:0048738** | cardiac muscle tissue development | 138 | 95 | 0.000696 |
| **GO:0018193** | peptidyl-amino acid modification | 1035 | 570 | 0.000741 |
| **GO:0030100** | regulation of endocytosis | 159 | 107 | 0.000773 |
| **GO:0042325** | regulation of phosphorylation | 1004 | 554 | 0.000779 |
| **GO:0051174** | regulation of phosphorus metabolic process | 1126 | 616 | 0.000825 |
| **GO:0051050** | positive regulation of transport | 709 | 402 | 0.000828 |
| **GO:2000147** | positive regulation of cell motility | 463 | 273 | 0.000841 |
| **GO:0043085** | positive regulation of catalytic activity | 960 | 531 | 0.000914 |
| **GO:0030048** | actin filament-based movement | 113 | 80 | 0.00094 |
| **GO:0019220** | regulation of phosphate metabolic process | 1125 | 615 | 0.000956 |
| **GO:0031623** | receptor internalization | 98 | 71 | 0.000992 |
| **GO:0006836** | neurotransmitter transport | 137 | 94 | 0.000993 |
| **GO:0031345** | negative regulation of cell projection organization | 125 | 87 | 0.000996 |
| **GO:0010977** | negative regulation of neuron projection development | 85 | 63 | 0.001143 |
| **GO:0032412** | regulation of ion transmembrane transporter activity | 222 | 142 | 0.001145 |
| **GO:0030178** | negative regulation of Wnt signaling pathway | 146 | 99 | 0.001179 |
| **GO:0031532** | actin cytoskeleton reorganization | 105 | 75 | 0.001196 |
| **GO:0010243** | response to organonitrogen compound | 613 | 351 | 0.001236 |
| **GO:0048598** | embryonic morphogenesis | 308 | 189 | 0.001311 |
| **GO:0008016** | regulation of heart contraction | 164 | 109 | 0.0015 |
| **GO:0006937** | regulation of muscle contraction | 119 | 83 | 0.001537 |
| **GO:0051241** | negative regulation of multicellular organismal process | 755 | 424 | 0.001593 |
| **GO:0007416** | synapse assembly | 126 | 87 | 0.00167 |
| **GO:0007229** | integrin-mediated signaling pathway | 104 | 74 | 0.001771 |
| **GO:0099173** | postsynapse organization | 99 | 71 | 0.001812 |
| **GO:0046903** | secretion | 638 | 363 | 0.001834 |
| **GO:0007163** | establishment or maintenance of cell polarity | 177 | 116 | 0.001975 |
| **GO:0071526** | semaphorin-plexin signaling pathway | 40 | 34 | 0.001997 |
| **GO:0050771** | negative regulation of axonogenesis | 43 | 36 | 0.00202 |
| **GO:0022898** | regulation of transmembrane transporter activity | 229 | 145 | 0.002021 |
| **GO:0055001** | muscle cell development | 118 | 82 | 0.002235 |
| **GO:1903522** | regulation of blood circulation | 188 | 122 | 0.002262 |
| **GO:0043412** | macromolecule modification | 3198 | 1641 | 0.002376 |
| **GO:0043408** | regulation of MAPK cascade | 534 | 308 | 0.002429 |
| **GO:0043087** | regulation of GTPase activity | 301 | 184 | 0.002542 |
| **GO:0031401** | positive regulation of protein modification process | 814 | 453 | 0.002569 |
| **GO:0046578** | regulation of Ras protein signal transduction | 139 | 94 | 0.002601 |
| **GO:1901699** | cellular response to nitrogen compound | 468 | 273 | 0.002746 |
| **GO:0032940** | secretion by cell | 571 | 327 | 0.002793 |
| **GO:0051966** | regulation of synaptic transmission, glutamatergic | 51 | 41 | 0.002884 |
| **GO:0008015** | blood circulation | 363 | 217 | 0.003065 |
| **GO:0051246** | regulation of protein metabolic process | 2072 | 1085 | 0.0031 |
| **GO:0098703** | calcium ion import across plasma membrane | 36 | 31 | 0.003323 |
| **GO:0060079** | excitatory postsynaptic potential | 45 | 37 | 0.003335 |
| **GO:0048813** | dendrite morphogenesis | 97 | 69 | 0.003994 |
| **GO:0010594** | regulation of endothelial cell migration | 154 | 102 | 0.004056 |
| **GO:0071417** | cellular response to organonitrogen compound | 413 | 243 | 0.004126 |
| **GO:0051146** | striated muscle cell differentiation | 170 | 111 | 0.004196 |
| **GO:0070588** | calcium ion transmembrane transport | 235 | 147 | 0.004516 |
| **GO:0070848** | response to growth factor | 516 | 297 | 0.004593 |
| **GO:0008360** | regulation of cell shape | 116 | 80 | 0.004638 |
| **GO:0021953** | central nervous system neuron differentiation | 74 | 55 | 0.004743 |

### Supplemental Table 6 (continued).

| GO ID | GO Description | Universe | COSMIC | Adjusted |
|---|---|---|---|---|
| **GO:0045229** | external encapsulating structure organization | 237 | 148 | 0.004792 |
| **GO:0051649** | establishment of localization in cell | 1698 | 897 | 0.004864 |
| **GO:0060048** | cardiac muscle contraction | 111 | 77 | 0.004996 |
| **GO:1901888** | regulation of cell junction assembly | 144 | 96 | 0.005129 |
| **GO:0090090** | negative regulation of canonical Wnt signaling pathway | 118 | 81 | 0.005334 |
| **GO:0033036** | macromolecule localization | 2245 | 1168 | 0.005414 |
| **GO:0006816** | calcium ion transport | 313 | 189 | 0.005544 |
| **GO:0060341** | regulation of cellular localization | 766 | 426 | 0.005581 |
| **GO:0043542** | endothelial cell migration | 194 | 124 | 0.005618 |
| **GO:0040013** | negative regulation of locomotion | 278 | 170 | 0.00585 |
| **GO:0006898** | receptor-mediated endocytosis | 214 | 135 | 0.005907 |
| **GO:0030198** | extracellular matrix organization | 234 | 146 | 0.005907 |
| **GO:0031400** | negative regulation of protein modification process | 409 | 240 | 0.006128 |
| **GO:0110053** | regulation of actin filament organization | 216 | 136 | 0.006298 |
| **GO:0060429** | epithelium development | 689 | 386 | 0.006326 |
| **GO:0030041** | actin filament polymerization | 129 | 87 | 0.007112 |
| **GO:0031098** | stress-activated protein kinase signaling cascade | 202 | 128 | 0.007442 |
| **GO:0060291** | long-term synaptic potentiation | 49 | 39 | 0.007819 |
| **GO:0043062** | extracellular structure organization | 235 | 146 | 0.008136 |
| **GO:1901379** | regulation of potassium ion transmembrane transport | 70 | 52 | 0.00878 |
| **GO:1902903** | regulation of supramolecular fiber organization | 311 | 187 | 0.008847 |
| **GO:0032880** | regulation of protein localization | 672 | 376 | 0.009649 |
| **GO:0043549** | regulation of kinase activity | 599 | 338 | 0.010664 |
| **GO:2001257** | regulation of cation channel activity | 153 | 100 | 0.011535 |
| **GO:0000122** | negative regulation of transcription by RNA polymerase II | 751 | 416 | 0.011781 |
| **GO:0002009** | morphogenesis of an epithelium | 275 | 167 | 0.012154 |
| **GO:0051403** | stress-activated MAPK cascade | 198 | 125 | 0.012154 |
| **GO:1990573** | potassium ion import across plasma membrane | 48 | 38 | 0.012766 |
| **GO:0120031** | plasma membrane bounded cell projection assembly | 483 | 277 | 0.013638 |
| **GO:0051258** | protein polymerization | 233 | 144 | 0.01366 |
| **GO:0045664** | regulation of neuron differentiation | 96 | 67 | 0.014735 |
| **GO:0007160** | cell-matrix adhesion | 195 | 123 | 0.015007 |
| **GO:0061572** | actin filament bundle organization | 136 | 90 | 0.015215 |
| **GO:0042327** | positive regulation of phosphorylation | 663 | 370 | 0.015238 |
| **GO:0007612** | learning | 53 | 41 | 0.015592 |
| **GO:0070252** | actin-mediated cell contraction | 86 | 61 | 0.016224 |
| **GO:0051338** | regulation of transferase activity | 718 | 398 | 0.016558 |
| **GO:0098901** | regulation of cardiac muscle cell action potential | 30 | 26 | 0.016851 |
| **GO:0051963** | regulation of synapse assembly | 58 | 44 | 0.017849 |
| **GO:0086065** | cell communication involved in cardiac conduction | 58 | 44 | 0.017849 |
| **GO:0051017** | actin filament bundle assembly | 133 | 88 | 0.018951 |
| **GO:0050678** | regulation of epithelial cell proliferation | 236 | 145 | 0.019404 |
| **GO:1990778** | protein localization to cell periphery | 279 | 168 | 0.021945 |
| **GO:0042592** | homeostatic process | 1249 | 666 | 0.022003 |
| **GO:0009792** | embryo development ending in birth or egg hatching | 198 | 124 | 0.022257 |
| **GO:0071363** | cellular response to growth factor stimulus | 499 | 284 | 0.022775 |
| **GO:0007215** | glutamate receptor signaling pathway | 44 | 35 | 0.023272 |
| **GO:0048640** | negative regulation of developmental growth | 60 | 45 | 0.023304 |
| **GO:0051961** | negative regulation of nervous system development | 85 | 60 | 0.023796 |
| **GO:0010959** | regulation of metal ion transport | 298 | 178 | 0.023909 |
| **GO:0097553** | calcium ion transmembrane import into cytosol | 146 | 95 | 0.024141 |
| **GO:0048762** | mesenchymal cell differentiation | 191 | 120 | 0.024283 |
| **GO:0048863** | stem cell differentiation | 155 | 100 | 0.02487 |
| **GO:0050879** | multicellular organismal movement | 41 | 33 | 0.025765 |
| **GO:0086091** | regulation of heart rate by cardiac conduction | 41 | 33 | 0.025765 |
| **GO:0050881** | musculoskeletal movement | 41 | 33 | 0.025765 |
| **GO:0032878** | regulation of establishment or maintenance of cell polarity | 26 | 23 | 0.026744 |
| **GO:0030031** | cell projection assembly | 498 | 283 | 0.027121 |
| **GO:0010232** | vascular transport | 87 | 61 | 0.027857 |
| **GO:0150104** | transport across blood-brain barrier | 87 | 61 | 0.027857 |
| **GO:0050919** | negative chemotaxis | 35 | 29 | 0.029641 |
| **GO:0050773** | regulation of dendrite development | 62 | 46 | 0.029645 |
| **GO:0030517** | negative regulation of axon extension | 32 | 27 | 0.030265 |
| **GO:0042982** | amyloid precursor protein metabolic process | 67 | 49 | 0.030628 |
| **GO:0007423** | sensory organ development | 271 | 163 | 0.030834 |
| **GO:0045785** | positive regulation of cell adhesion | 369 | 215 | 0.032573 |

### Supplemental Table 6 (continued).

| GO ID | GO Description | Universe | COSMIC | Adjusted |
|---|---|---|---|---|
| **GO:0009611** | response to wounding | 392 | 227 | 0.034051 |
| **GO:0050890** | cognition | 156 | 100 | 0.035799 |
| **GO:0001932** | regulation of protein phosphorylation | 882 | 479 | 0.035846 |
| **GO:0031323** | regulation of cellular metabolic process | 4966 | 2482 | 0.036928 |
| **GO:0051899** | membrane depolarization | 64 | 47 | 0.037241 |
| **GO:0001944** | vasculature development | 492 | 279 | 0.037414 |
| **GO:0002090** | regulation of receptor internalization | 43 | 34 | 0.037523 |
| **GO:0099643** | signal release from synapse | 91 | 63 | 0.037593 |
| **GO:0007269** | neurotransmitter secretion | 91 | 63 | 0.037593 |
| **GO:0010634** | positive regulation of epithelial cell migration | 133 | 87 | 0.038739 |
| **GO:0043254** | regulation of protein-containing complex assembly | 332 | 195 | 0.039459 |
| **GO:0045773** | positive regulation of axon extension | 22 | 20 | 0.039621 |
| **GO:0018108** | peptidyl-tyrosine phosphorylation | 246 | 149 | 0.040743 |
| **GO:0090596** | sensory organ morphogenesis | 119 | 79 | 0.040966 |
| **GO:0021954** | central nervous system neuron development | 40 | 32 | 0.042039 |
| **GO:2000249** | regulation of actin cytoskeleton reorganization | 40 | 32 | 0.042039 |
| **GO:0001933** | negative regulation of protein phosphorylation | 261 | 157 | 0.042119 |
| **GO:0018212** | peptidyl-tyrosine modification | 248 | 150 | 0.042279 |
| **GO:0010562** | positive regulation of phosphorus metabolic process | 722 | 397 | 0.042279 |
| **GO:0045937** | positive regulation of phosphate metabolic process | 722 | 397 | 0.042279 |
| **GO:0051668** | localization within membrane | 570 | 319 | 0.042433 |
| **GO:0048592** | eye morphogenesis | 81 | 57 | 0.042492 |
| **GO:0033554** | cellular response to stress | 1640 | 858 | 0.043396 |
| **GO:0045055** | regulated exocytosis | 162 | 103 | 0.044095 |
| **GO:1901021** | positive regulation of calcium ion transmembrane transporter activity | 37 | 30 | 0.046133 |
| **GO:0051347** | positive regulation of transferase activity | 447 | 255 | 0.046166 |
| **GO:0044092** | negative regulation of molecular function | 758 | 415 | 0.046963 |
| **GO:0006935** | chemotaxis | 480 | 272 | 0.049239 |
| **GO:0042330** | taxis | 480 | 272 | 0.049239 |

**Supplemental Table 7.** Significant GO:BP enrichments for all CLINVAR G4 mutations.

| GO ID | GO Description | Universe | CLINVAR | Adjusted P-value |
|---|---|---|---|---|
| **GO:0006941** | striated muscle contraction | 142 | 32 | 1.1E-11 |
| **GO:0060048** | cardiac muscle contraction | 111 | 27 | 2.1E-10 |
| **GO:0050905** | neuromuscular process | 73 | 22 | 4.5E-10 |
| **GO:0035637** | multicellular organismal signaling | 128 | 28 | 1.1E-09 |
| **GO:0001508** | action potential | 117 | 26 | 5.2E-09 |
| **GO:0030048** | actin filament-based movement | 113 | 25 | 1.5E-08 |
| **GO:0070252** | actin-mediated cell contraction | 86 | 20 | 7.0E-07 |
| **GO:0061337** | cardiac conduction | 91 | 20 | 1.9E-06 |
| **GO:0086001** | cardiac muscle cell action potential | 73 | 18 | 2.0E-06 |
| **GO:0006937** | regulation of muscle contraction | 119 | 22 | 7.3E-06 |
| **GO:0086003** | cardiac muscle cell contraction | 62 | 15 | 5.8E-05 |
| **GO:0051899** | membrane depolarization | 64 | 15 | 8.9E-05 |
| **GO:1903115** | regulation of actin filament-based movement | 32 | 11 | 9.0E-05 |
| **GO:0060415** | muscle tissue morphogenesis | 47 | 13 | 9.1E-05 |
| **GO:0048644** | muscle organ morphogenesis | 47 | 13 | 9.1E-05 |
| **GO:0002027** | regulation of heart rate | 84 | 17 | 1.1E-04 |
| **GO:0098900** | regulation of action potential | 48 | 13 | 1.2E-04 |
| **GO:0086002** | cardiac muscle cell action potential involved in contraction | 48 | 13 | 1.2E-04 |
| **GO:0050954** | sensory perception of mechanical stimulus | 106 | 18 | 5.7E-04 |
| **GO:0086065** | cell communication involved in cardiac conduction | 58 | 13 | 1.0E-03 |
| **GO:0006942** | regulation of striated muscle contraction | 78 | 15 | 1.1E-03 |
| **GO:0007605** | sensory perception of sound | 100 | 17 | 1.1E-03 |
| **GO:0086091** | regulation of heart rate by cardiac conduction | 41 | 11 | 1.2E-03 |
| **GO:0050881** | musculoskeletal movement | 41 | 11 | 1.2E-03 |
| **GO:0050879** | multicellular organismal movement | 41 | 11 | 1.2E-03 |
| **GO:0019226** | transmission of nerve impulse | 42 | 11 | 1.5E-03 |
| **GO:0055008** | cardiac muscle tissue morphogenesis | 42 | 11 | 1.5E-03 |
| **GO:0048738** | cardiac muscle tissue development | 138 | 20 | 1.5E-03 |
| **GO:0055001** | muscle cell development | 118 | 18 | 2.5E-03 |
| **GO:0003009** | skeletal muscle contraction | 36 | 10 | 2.5E-03 |
| **GO:0014706** | striated muscle tissue development | 144 | 20 | 2.8E-03 |
| **GO:0086005** | ventricular cardiac muscle cell action potential | 29 | 9 | 3.1E-03 |
| **GO:0050885** | neuromuscular process controlling balance | 16 | 7 | 3.4E-03 |
| **GO:0098901** | regulation of cardiac muscle cell action potential | 30 | 9 | 4.2E-03 |
| **GO:0003229** | ventricular cardiac muscle tissue development | 39 | 10 | 5.3E-03 |
| **GO:0048592** | eye morphogenesis | 81 | 14 | 8.1E-03 |
| **GO:0055010** | ventricular cardiac muscle tissue morphogenesis | 33 | 9 | 9.3E-03 |
| **GO:0055117** | regulation of cardiac muscle contraction | 61 | 12 | 9.8E-03 |
| **GO:0086019** | cell-cell signaling involved in cardiac conduction | 34 | 9 | 1.2E-02 |
| **GO:0008306** | associative learning | 19 | 7 | 1.2E-02 |
| **GO:0086010** | membrane depolarization during action potential | 35 | 9 | 1.5E-02 |
| **GO:0050953** | sensory perception of light stimulus | 138 | 18 | 1.9E-02 |
| **GO:0086004** | regulation of cardiac muscle cell contraction | 28 | 8 | 2.0E-02 |
| **GO:0043589** | skin morphogenesis | 5 | 4 | 2.6E-02 |
| **GO:0002028** | regulation of sodium ion transport | 68 | 12 | 2.7E-02 |
| **GO:0048661** | positive regulation of smooth muscle cell proliferation | 49 | 10 | 3.6E-02 |
| **GO:0048483** | autonomic nervous system development | 31 | 8 | 4.1E-02 |
| **GO:0060348** | bone development | 108 | 15 | 4.5E-02 |

**Supplemental Table 8**. Significant GO:BP enrichments for COSMIC and CLINVAR G4 mutations leading to the loss of a G4.

| GO ID | GO Description | Universe | COSMIC and CLINVAR | Adjusted P-value |
|---|---|---|---|---|
| **GO:0032502** | developmental process | 4584 | 1138 | 3.23E-24 |
| **GO:0048856** | anatomical structure development | 4152 | 1043 | 2.82E-23 |
| **GO:0007399** | nervous system development | 1648 | 488 | 1.15E-22 |
| **GO:0009653** | anatomical structure morphogenesis | 1871 | 539 | 2.18E-22 |
| **GO:0048731** | system development | 2976 | 784 | 9.30E-22 |
| **GO:0007275** | multicellular organism development | 3266 | 842 | 1.00E-20 |
| **GO:0032501** | multicellular organismal process | 5319 | 1270 | 1.40E-20 |
| **GO:0030154** | cell differentiation | 2860 | 742 | 3.10E-18 |
| **GO:0048699** | generation of neurons | 969 | 309 | 3.31E-18 |
| **GO:0048869** | cellular developmental process | 2879 | 745 | 5.24E-18 |
| **GO:0016043** | cellular component organization | 5370 | 1259 | 6.78E-17 |
| **GO:0022008** | neurogenesis | 1096 | 335 | 1.06E-16 |
| **GO:0030182** | neuron differentiation | 923 | 292 | 1.76E-16 |
| **GO:0071840** | cellular component organization or biogenesis | 5539 | 1277 | 2.38E-14 |
| **GO:0023051** | regulation of signaling | 2733 | 694 | 3.06E-14 |
| **GO:0010646** | regulation of cell communication | 2727 | 692 | 4.16E-14 |
| **GO:0048666** | neuron development | 729 | 236 | 4.22E-14 |
| **GO:0000904** | cell morphogenesis involved in differentiation | 495 | 175 | 7.97E-14 |
| **GO:0048468** | cell development | 1355 | 385 | 9.16E-14 |
| **GO:0048513** | animal organ development | 2176 | 567 | 3.84E-13 |
| **GO:0023052** | signaling | 5190 | 1196 | 9.50E-13 |
| **GO:0032989** | cellular component morphogenesis | 549 | 186 | 1.02E-12 |
| **GO:0007154** | cell communication | 5221 | 1200 | 2.02E-12 |
| **GO:0031175** | neuron projection development | 656 | 212 | 2.53E-12 |
| **GO:0048667** | cell morphogenesis involved in neuron differentiation | 381 | 140 | 3.85E-12 |
| **GO:0034330** | cell junction organization | 518 | 176 | 4.81E-12 |
| **GO:0000902** | cell morphogenesis | 718 | 226 | 6.91E-12 |
| **GO:0048812** | neuron projection morphogenesis | 441 | 155 | 9.70E-12 |
| **GO:0007010** | cytoskeleton organization | 1309 | 365 | 1.42E-11 |
| **GO:0009966** | regulation of signal transduction | 2464 | 621 | 1.59E-11 |
| **GO:0009887** | animal organ morphogenesis | 582 | 190 | 2.87E-11 |
| **GO:0007155** | cell adhesion | 1216 | 342 | 3.26E-11 |
| **GO:0120036** | plasma membrane bounded cell projection organization | 1144 | 325 | 3.84E-11 |
| **GO:0120039** | plasma membrane bounded cell projection morphogenesis | 455 | 157 | 3.85E-11 |
| **GO:0048858** | cell projection morphogenesis | 459 | 157 | 8.90E-11 |
| **GO:0030030** | cell projection organization | 1164 | 328 | 9.33E-11 |
| **GO:0032990** | cell part morphogenesis | 469 | 159 | 1.42E-10 |
| **GO:0030029** | actin filament-based process | 717 | 219 | 5.92E-10 |
| **GO:0061564** | axon development | 323 | 118 | 1.06E-09 |
| **GO:0007409** | axonogenesis | 298 | 111 | 1.25E-09 |
| **GO:0050808** | synapse organization | 275 | 104 | 2.31E-09 |
| **GO:0050793** | regulation of developmental process | 1810 | 465 | 4.44E-09 |
| **GO:0051128** | regulation of cellular component organization | 1929 | 490 | 6.77E-09 |
| **GO:0007165** | signal transduction | 4776 | 1078 | 4.27E-08 |
| **GO:0016477** | cell migration | 1210 | 325 | 7.79E-08 |
| **GO:0048870** | cell motility | 1362 | 359 | 8.30E-08 |
| **GO:0098609** | cell-cell adhesion | 743 | 217 | 8.39E-08 |
| **GO:0051716** | cellular response to stimulus | 5982 | 1316 | 9.92E-08 |
| **GO:0035556** | intracellular signal transduction | 2168 | 534 | 1.26E-07 |
| **GO:0048583** | regulation of response to stimulus | 3327 | 777 | 1.74E-07 |
| **GO:0051239** | regulation of multicellular organismal process | 2117 | 522 | 1.84E-07 |
| **GO:0099537** | trans-synaptic signaling | 501 | 157 | 2.02E-07 |
| **GO:0099536** | synaptic signaling | 522 | 162 | 2.28E-07 |
| **GO:0009888** | tissue development | 1239 | 329 | 2.43E-07 |
| **GO:0098916** | anterograde trans-synaptic signaling | 495 | 155 | 2.81E-07 |
| **GO:0007268** | chemical synaptic transmission | 495 | 155 | 2.81E-07 |
| **GO:0065007** | biological regulation | 10721 | 2221 | 3.01E-07 |
| **GO:0072359** | circulatory system development | 729 | 211 | 3.69E-07 |
| **GO:0065008** | regulation of biological quality | 2937 | 692 | 5.65E-07 |
| **GO:0050794** | regulation of cellular process | 9523 | 1993 | 7.40E-07 |
| **GO:0007267** | cell-cell signaling | 1269 | 333 | 7.84E-07 |
| **GO:0003008** | system process | 1358 | 351 | 1.64E-06 |
| **GO:0007417** | central nervous system development | 584 | 173 | 2.68E-06 |
| **GO:0009987** | cellular process | 14783 | 2936 | 4.03E-06 |

### Supplemental Table 8 (continued).

| GO ID | GO Description | Universe | COSMIC and CLINVAR | Adjusted P-value |
|---|---|---|---|---|
| **GO:0040011** | locomotion | 1075 | 285 | 5.65E-06 |
| **GO:0050789** | regulation of biological process | 10085 | 2089 | 8.77E-06 |
| **GO:1905114** | cell surface receptor signaling pathway involved in cell-cell signaling | 388 | 123 | 1.03E-05 |
| **GO:0030036** | actin cytoskeleton organization | 637 | 183 | 1.15E-05 |
| **GO:0050804** | modulation of chemical synaptic transmission | 252 | 88 | 1.18E-05 |
| **GO:0030048** | actin filament-based movement | 113 | 49 | 1.44E-05 |
| **GO:0099177** | regulation of trans-synaptic signaling | 253 | 88 | 1.46E-05 |
| **GO:0006812** | cation transport | 886 | 240 | 1.55E-05 |
| **GO:0030001** | metal ion transport | 678 | 192 | 1.58E-05 |
| **GO:0010975** | regulation of neuron projection development | 288 | 97 | 1.61E-05 |
| **GO:0060047** | heart contraction | 187 | 70 | 1.80E-05 |
| **GO:0006811** | ion transport | 1187 | 307 | 1.99E-05 |
| **GO:0048518** | positive regulation of biological process | 5294 | 1158 | 2.05E-05 |
| **GO:0034329** | cell junction assembly | 341 | 110 | 2.35E-05 |
| **GO:0032879** | regulation of localization | 1615 | 400 | 2.54E-05 |
| **GO:0044057** | regulation of system process | 392 | 122 | 3.78E-05 |
| **GO:0007411** | axon guidance | 169 | 64 | 4.59E-05 |
| **GO:0097485** | neuron projection guidance | 169 | 64 | 4.59E-05 |
| **GO:0051179** | localization | 4343 | 964 | 4.59E-05 |
| **GO:0048522** | positive regulation of cellular process | 4704 | 1036 | 5.30E-05 |
| **GO:0007507** | heart development | 338 | 108 | 5.46E-05 |
| **GO:0006936** | muscle contraction | 260 | 88 | 6.00E-05 |
| **GO:0055085** | transmembrane transport | 1060 | 276 | 6.09E-05 |
| **GO:0034220** | ion transmembrane transport | 816 | 221 | 6.22E-05 |
| **GO:0042391** | regulation of membrane potential | 319 | 103 | 6.34E-05 |
| **GO:0048523** | negative regulation of cellular process | 3896 | 872 | 6.65E-05 |
| **GO:0003015** | heart process | 193 | 70 | 7.53E-05 |
| **GO:0007166** | cell surface receptor signaling pathway | 2271 | 536 | 8.29E-05 |
| **GO:0003012** | muscle system process | 305 | 99 | 8.73E-05 |
| **GO:1902531** | regulation of intracellular signal transduction | 1429 | 356 | 9.28E-05 |
| **GO:0061061** | muscle structure development | 407 | 124 | 1.07E-04 |
| **GO:0040012** | regulation of locomotion | 842 | 225 | 1.46E-04 |
| **GO:0098655** | cation transmembrane transport | 656 | 182 | 1.81E-04 |
| **GO:0048646** | anatomical structure formation involved in morphogenesis | 751 | 203 | 2.65E-04 |
| **GO:0048519** | negative regulation of biological process | 4381 | 964 | 2.73E-04 |
| **GO:0048729** | tissue morphogenesis | 348 | 108 | 2.78E-04 |
| **GO:0007416** | synapse assembly | 126 | 50 | 2.85E-04 |
| **GO:0022603** | regulation of anatomical structure morphogenesis | 700 | 191 | 3.10E-04 |
| **GO:0051960** | regulation of nervous system development | 257 | 85 | 3.10E-04 |
| **GO:0030334** | regulation of cell migration | 767 | 206 | 3.51E-04 |
| **GO:0050905** | neuromuscular process | 73 | 34 | 3.51E-04 |
| **GO:0031589** | cell-substrate adhesion | 290 | 93 | 3.99E-04 |
| **GO:0050896** | response to stimulus | 7117 | 1503 | 4.02E-04 |
| **GO:0031344** | regulation of cell projection organization | 467 | 136 | 4.37E-04 |
| **GO:0003013** | circulatory system process | 443 | 130 | 5.12E-04 |
| **GO:0065009** | regulation of molecular function | 2121 | 498 | 5.42E-04 |
| **GO:0006810** | transport | 3620 | 807 | 5.65E-04 |
| **GO:0120035** | regulation of plasma membrane bounded cell projection organization | 453 | 132 | 6.34E-04 |
| **GO:0008015** | blood circulation | 363 | 110 | 7.79E-04 |
| **GO:0030111** | regulation of Wnt signaling pathway | 274 | 88 | 7.79E-04 |
| **GO:0007420** | brain development | 384 | 115 | 8.24E-04 |
| **GO:0051130** | positive regulation of cellular component organization | 844 | 221 | 9.67E-04 |
| **GO:2000145** | regulation of cell motility | 818 | 215 | 1.04E-03 |
| **GO:0006996** | organelle organization | 3147 | 708 | 1.04E-03 |
| **GO:0070252** | actin-mediated cell contraction | 86 | 37 | 1.15E-03 |
| **GO:0051056** | regulation of small GTPase mediated signal transduction | 238 | 78 | 1.44E-03 |
| **GO:0050877** | nervous system process | 755 | 200 | 1.46E-03 |
| **GO:0051234** | establishment of localization | 3775 | 834 | 1.55E-03 |
| **GO:0060048** | cardiac muscle contraction | 111 | 44 | 1.57E-03 |
| **GO:0051094** | positive regulation of developmental process | 967 | 247 | 1.64E-03 |
| **GO:0098662** | inorganic cation transmembrane transport | 572 | 158 | 1.65E-03 |
| **GO:0051963** | regulation of synapse assembly | 58 | 28 | 1.69E-03 |
| **GO:0060322** | head development | 402 | 118 | 1.74E-03 |
| **GO:0001667** | ameboidal-type cell migration | 333 | 101 | 2.14E-03 |

### Supplemental Table 8 (continued).

| GO ID | GO Description | Universe | COSMIC and CLINVAR | Adjusted P-value |
|---|---|---|---|---|
| **GO:0048638** | regulation of developmental growth | 167 | 59 | 2.18E-03 |
| **GO:0007167** | enzyme-linked receptor protein signaling pathway | 795 | 208 | 2.19E-03 |
| **GO:0008016** | regulation of heart contraction | 164 | 58 | 2.57E-03 |
| **GO:0051049** | regulation of transport | 1327 | 324 | 2.69E-03 |
| **GO:0006941** | striated muscle contraction | 142 | 52 | 2.76E-03 |
| **GO:0045595** | regulation of cell differentiation | 1129 | 281 | 2.82E-03 |
| **GO:0097435** | supramolecular fiber organization | 710 | 188 | 3.17E-03 |
| **GO:0035637** | multicellular organismal signaling | 128 | 48 | 3.20E-03 |
| **GO:0098660** | inorganic ion transmembrane transport | 622 | 168 | 3.22E-03 |
| **GO:0001505** | regulation of neurotransmitter levels | 143 | 52 | 3.47E-03 |
| **GO:0048598** | embryonic morphogenesis | 308 | 94 | 3.66E-03 |
| **GO:0010647** | positive regulation of cell communication | 1374 | 333 | 3.80E-03 |
| **GO:0006816** | calcium ion transport | 313 | 95 | 4.16E-03 |
| **GO:0009967** | positive regulation of signal transduction | 1255 | 307 | 4.32E-03 |
| **GO:0099565** | chemical synaptic transmission, postsynaptic | 51 | 25 | 4.41E-03 |
| **GO:0009719** | response to endogenous stimulus | 1076 | 268 | 4.62E-03 |
| **GO:0099587** | inorganic ion import across plasma membrane | 112 | 43 | 5.31E-03 |
| **GO:0098659** | inorganic cation import across plasma membrane | 112 | 43 | 5.31E-03 |
| **GO:0009790** | embryo development | 505 | 140 | 5.37E-03 |
| **GO:0098742** | cell-cell adhesion via plasma-membrane adhesion molecules | 195 | 65 | 5.93E-03 |
| **GO:0007169** | transmembrane receptor protein tyrosine kinase signaling pathway | 511 | 141 | 6.44E-03 |
| **GO:0023057** | negative regulation of signaling | 1067 | 265 | 6.57E-03 |
| **GO:1903522** | regulation of blood circulation | 188 | 63 | 6.84E-03 |
| **GO:0050807** | regulation of synapse organization | 124 | 46 | 7.05E-03 |
| **GO:0001508** | action potential | 117 | 44 | 7.54E-03 |
| **GO:0023056** | positive regulation of signaling | 1380 | 332 | 7.60E-03 |
| **GO:0051962** | positive regulation of nervous system development | 147 | 52 | 8.38E-03 |
| **GO:0198738** | cell-cell signaling by wnt | 343 | 101 | 8.47E-03 |
| **GO:0010631** | epithelial cell migration | 261 | 81 | 8.86E-03 |
| **GO:0043269** | regulation of ion transport | 458 | 128 | 9.01E-03 |
| **GO:0016358** | dendrite development | 155 | 54 | 9.18E-03 |
| **GO:0050803** | regulation of synapse structure or activity | 129 | 47 | 9.61E-03 |
| **GO:1903115** | regulation of actin filament-based movement | 32 | 18 | 9.75E-03 |
| **GO:1901888** | regulation of cell junction assembly | 144 | 51 | 9.80E-03 |
| **GO:0040008** | regulation of growth | 416 | 118 | 9.86E-03 |
| **GO:0007612** | learning | 53 | 25 | 1.00E-02 |
| **GO:0022607** | cellular component assembly | 2547 | 575 | 1.03E-02 |
| **GO:0010648** | negative regulation of cell communication | 1060 | 262 | 1.06E-02 |
| **GO:0042221** | response to chemical | 2943 | 656 | 1.07E-02 |
| **GO:0086001** | cardiac muscle cell action potential | 73 | 31 | 1.16E-02 |
| **GO:0071495** | cellular response to endogenous stimulus | 974 | 243 | 1.17E-02 |
| **GO:0090130** | tissue migration | 267 | 82 | 1.17E-02 |
| **GO:0090132** | epithelium migration | 263 | 81 | 1.19E-02 |
| **GO:0090596** | sensory organ morphogenesis | 119 | 44 | 1.21E-02 |
| **GO:0048738** | cardiac muscle tissue development | 138 | 49 | 1.34E-02 |
| **GO:0007269** | neurotransmitter secretion | 91 | 36 | 1.45E-02 |
| **GO:0099643** | signal release from synapse | 91 | 36 | 1.45E-02 |
| **GO:0016055** | Wnt signaling pathway | 339 | 99 | 1.47E-02 |
| **GO:0040007** | growth | 515 | 140 | 1.52E-02 |
| **GO:0048589** | developmental growth | 273 | 83 | 1.55E-02 |
| **GO:0098703** | calcium ion import across plasma membrane | 36 | 19 | 1.76E-02 |
| **GO:0099003** | vesicle-mediated transport in synapse | 121 | 44 | 1.91E-02 |
| **GO:0014706** | striated muscle tissue development | 144 | 50 | 2.13E-02 |
| **GO:0016310** | phosphorylation | 1444 | 342 | 2.13E-02 |
| **GO:0007517** | muscle organ development | 179 | 59 | 2.24E-02 |
| **GO:0034762** | regulation of transmembrane transport | 385 | 109 | 2.30E-02 |
| **GO:0006836** | neurotransmitter transport | 137 | 48 | 2.39E-02 |
| **GO:0044087** | regulation of cellular component biogenesis | 774 | 197 | 2.41E-02 |
| **GO:0086003** | cardiac muscle cell contraction | 62 | 27 | 2.50E-02 |
| **GO:0007611** | learning or memory | 115 | 42 | 2.61E-02 |
| **GO:0009968** | negative regulation of signal transduction | 1009 | 248 | 2.74E-02 |
| **GO:0090257** | regulation of muscle system process | 157 | 53 | 2.86E-02 |
| **GO:0035249** | synaptic transmission, glutamatergic | 66 | 28 | 3.09E-02 |
| **GO:0060560** | developmental growth involved in morphogenesis | 135 | 47 | 3.44E-02 |

### Supplemental Table 8 (continued).

| GO ID | GO Description | Universe | COSMIC and CLINVAR | Adjusted P-value |
|---|---|---|---|---|
| **GO:0002009** | morphogenesis of an epithelium | 275 | 82 | 3.61E-02 |
| **GO:0007264** | small GTPase mediated signal transduction | 389 | 109 | 3.61E-02 |
| **GO:0002027** | regulation of heart rate | 84 | 33 | 3.80E-02 |
| **GO:0043542** | endothelial cell migration | 194 | 62 | 3.82E-02 |
| **GO:0051668** | localization within membrane | 570 | 150 | 4.30E-02 |
| **GO:0086091** | regulation of heart rate by cardiac conduction | 41 | 20 | 4.31E-02 |
| **GO:0060828** | regulation of canonical Wnt signaling pathway | 211 | 66 | 4.44E-02 |
| **GO:0051965** | positive regulation of synapse assembly | 35 | 18 | 4.46E-02 |
| **GO:0048588** | developmental cell growth | 129 | 45 | 4.73E-02 |
| **GO:0099504** | synaptic vesicle cycle | 114 | 41 | 4.75E-02 |
| **GO:0042127** | regulation of cell population proliferation | 1218 | 291 | 4.82E-02 |
| **GO:0050890** | cognition | 156 | 52 | 4.83E-02 |
| **GO:0007158** | neuron cell-cell adhesion | 16 | 11 | 4.94E-02 |

**Supplemental Table 9.** Significant GO:BP enrichments for COSMIC G4 mutations leading to the loss of a G4.

| GO ID | GO Description | Universe | COSMIC | Adjusted P-value |
|---|---|---|---|---|
| **GO:0009653** | anatomical structure morphogenesis | 1871 | 381 | 1.50E-10 |
| **GO:0016043** | cellular component organization | 5370 | 931 | 4.07E-10 |
| **GO:0032502** | developmental process | 4584 | 804 | 5.37E-09 |
| **GO:0071840** | cellular component organization or biogenesis | 5539 | 945 | 1.77E-08 |
| **GO:0048856** | anatomical structure development | 4152 | 734 | 1.85E-08 |
| **GO:0048699** | generation of neurons | 969 | 216 | 1.86E-08 |
| **GO:0007399** | nervous system development | 1648 | 333 | 2.19E-08 |
| **GO:0030182** | neuron differentiation | 923 | 204 | 1.65E-07 |
| **GO:0032501** | multicellular organismal process | 5319 | 904 | 2.10E-07 |
| **GO:0048731** | system development | 2976 | 543 | 2.40E-07 |
| **GO:0022008** | Neurogenesis | 1096 | 233 | 3.56E-07 |
| **GO:0007275** | multicellular organism development | 3266 | 587 | 4.31E-07 |
| **GO:0048666** | neuron development | 729 | 167 | 5.80E-07 |
| **GO:0030154** | cell differentiation | 2860 | 521 | 8.72E-07 |
| **GO:0048869** | cellular developmental process | 2879 | 523 | 1.24E-06 |
| **GO:0048468** | cell development | 1355 | 273 | 2.92E-06 |
| **GO:0010646** | regulation of cell communication | 2727 | 493 | 9.55E-06 |
| **GO:0023051** | regulation of signaling | 2733 | 492 | 1.83E-05 |
| **GO:0034330** | cell junction organization | 518 | 122 | 2.98E-05 |
| **GO:0031175** | neuron projection development | 656 | 147 | 3.28E-05 |
| **GO:0009966** | regulation of signal transduction | 2464 | 447 | 4.09E-05 |
| **GO:0048513** | animal organ development | 2176 | 401 | 4.17E-05 |
| **GO:0051128** | regulation of cellular component organization | 1929 | 360 | 6.48E-05 |
| **GO:0000904** | cell morphogenesis involved in differentiation | 495 | 116 | 9.06E-05 |
| **GO:0007010** | cytoskeleton organization | 1309 | 256 | 0.000164105 |
| **GO:0023052** | Signaling | 5190 | 860 | 0.000200273 |
| **GO:0007154** | cell communication | 5221 | 863 | 0.000302145 |
| **GO:0007155** | cell adhesion | 1216 | 239 | 0.000305302 |
| **GO:0050808** | synapse organization | 275 | 72 | 0.000421197 |
| **GO:0009887** | animal organ morphogenesis | 582 | 129 | 0.000483951 |
| **GO:0032989** | cellular component morphogenesis | 549 | 123 | 0.000504882 |
| **GO:0048667** | cell morphogenesis involved in neuron differentiation | 381 | 91 | 0.001066523 |
| **GO:0061564** | axon development | 323 | 80 | 0.001103984 |
| **GO:0065007** | biological regulation | 10721 | 1650 | 0.001653583 |
| **GO:0048812** | neuron projection morphogenesis | 441 | 101 | 0.002100476 |
| **GO:0120036** | plasma membrane bounded cell projection organization | 1144 | 222 | 0.002358738 |
| **GO:0048583** | regulation of response to stimulus | 3327 | 568 | 0.002363352 |
| **GO:0007409** | Axonogenesis | 298 | 74 | 0.002622931 |
| **GO:0003008** | system process | 1358 | 257 | 0.002627434 |
| **GO:0030030** | cell projection organization | 1164 | 225 | 0.002676372 |
| **GO:0035556** | intracellular signal transduction | 2168 | 387 | 0.002733973 |
| **GO:0032879** | regulation of localization | 1615 | 298 | 0.003442374 |
| **GO:0051239** | regulation of multicellular organismal process | 2117 | 378 | 0.003636668 |
| **GO:0000902** | cell morphogenesis | 718 | 149 | 0.003637049 |
| **GO:0007165** | signal transduction | 4776 | 786 | 0.003677398 |
| **GO:0098609** | cell-cell adhesion | 743 | 153 | 0.004206573 |
| **GO:0120039** | plasma membrane bounded cell projection morphogenesis | 455 | 102 | 0.00520267 |
| **GO:0006811** | ion transport | 1187 | 227 | 0.005379723 |
| **GO:0051179** | Localization | 4343 | 718 | 0.007233245 |
| **GO:0044057** | regulation of system process | 392 | 90 | 0.007238929 |
| **GO:0051716** | cellular response to stimulus | 5982 | 962 | 0.007709961 |
| **GO:0048858** | cell projection morphogenesis | 459 | 102 | 0.007839606 |
| **GO:0065008** | regulation of biological quality | 2937 | 503 | 0.00859953 |
| **GO:0097485** | neuron projection guidance | 169 | 47 | 0.009226901 |
| **GO:0007411** | axon guidance | 169 | 47 | 0.009226901 |
| **GO:0009888** | tissue development | 1239 | 234 | 0.009565902 |
| **GO:0050794** | regulation of cellular process | 9523 | 1473 | 0.010601198 |
| **GO:0006812** | cation transport | 886 | 175 | 0.011607738 |
| **GO:0032990** | cell part morphogenesis | 469 | 103 | 0.012164938 |
| **GO:0055085** | transmembrane transport | 1060 | 203 | 0.017138186 |
| **GO:0048598** | embryonic morphogenesis | 308 | 73 | 0.018555392 |
| **GO:0007417** | central nervous system development | 584 | 122 | 0.022813937 |
| **GO:0009790** | embryo development | 505 | 108 | 0.02527435 |
| **GO:0031589** | cell-substrate adhesion | 290 | 69 | 0.028200129 |

### Supplemental Table 9 (continued).

| GO ID | GO Description | Universe Count | COSMIC Count | Adjusted P-value |
|---|---|---|---|---|
| **GO:0040011** | Locomotion | 1075 | 204 | 0.029753799 |
| **GO:0050793** | regulation of developmental process | 1810 | 322 | 0.033576014 |
| **GO:0050905** | neuromuscular process | 73 | 25 | 0.036424405 |
| **GO:0048523** | negative regulation of cellular process | 3896 | 643 | 0.039903379 |
| **GO:0010975** | regulation of neuron projection development | 288 | 68 | 0.042350124 |
| **GO:0052697** | xenobiotic glucuronidation | 11 | 8 | 0.042820644 |
| **GO:0016477** | cell migration | 1210 | 225 | 0.043542025 |
| **GO:0003013** | circulatory system process | 443 | 96 | 0.043990329 |
| **GO:0048646** | anatomical structure formation involved in morphogenesis | 751 | 149 | 0.045881349 |
| **GO:0003012** | muscle system process | 305 | 71 | 0.046497064 |
| **GO:0048519** | negative regulation of biological process | 4381 | 715 | 0.049876587 |

**Supplemental Table 10.** Significant GO:BP enrichments for CLINVAR G4 mutations leading to the loss of a G4.

| GO ID | GO Description | Universe | CLINVAR | Adjusted P-value |
|---|---|---|---|---|
| **GO:0006936** | muscle contraction | 260 | 21 | 1.07E-06 |
| **GO:0003012** | muscle system process | 305 | 22 | 1.86E-06 |
| **GO:0060048** | cardiac muscle contraction | 111 | 14 | 5.31E-06 |
| **GO:0006941** | striated muscle contraction | 142 | 15 | 9.47E-06 |
| **GO:0060047** | heart contraction | 187 | 16 | 3.82E-05 |
| **GO:0003015** | heart process | 193 | 16 | 4.82E-05 |
| **GO:0086003** | cardiac muscle cell contraction | 62 | 10 | 6.95E-05 |
| **GO:0086002** | cardiac muscle cell action potential involved in contraction | 48 | 9 | 9.09E-05 |
| **GO:0070252** | actin-mediated cell contraction | 86 | 11 | 1.12E-04 |
| **GO:0030048** | actin filament-based movement | 113 | 12 | 1.84E-04 |
| **GO:0086001** | cardiac muscle cell action potential | 73 | 10 | 2.19E-04 |
| **GO:0008016** | regulation of heart contraction | 164 | 13 | 8.37E-04 |
| **GO:0001508** | action potential | 117 | 11 | 1.31E-03 |
| **GO:0035637** | multicellular organismal signaling | 128 | 11 | 2.48E-03 |
| **GO:0008015** | blood circulation | 363 | 18 | 2.56E-03 |
| **GO:0003013** | circulatory system process | 443 | 20 | 2.56E-03 |
| **GO:1903522** | regulation of blood circulation | 188 | 13 | 2.64E-03 |
| **GO:0060348** | bone development | 108 | 10 | 3.32E-03 |
| **GO:0086005** | ventricular cardiac muscle cell action potential | 29 | 6 | 4.45E-03 |
| **GO:0061337** | cardiac conduction | 91 | 9 | 5.54E-03 |
| **GO:0044057** | regulation of system process | 392 | 17 | 1.65E-02 |
| **GO:0002027** | regulation of heart rate | 84 | 8 | 2.02E-02 |
| **GO:0001501** | skeletal system development | 289 | 14 | 2.76E-02 |
| **GO:0030279** | negative regulation of ossification | 26 | 5 | 2.93E-02 |
| **GO:0060537** | muscle tissue development | 218 | 12 | 3.16E-02 |

**Supplemental Table 11:** Significant GO:BP enrichments for COSMIC and CLINVAR G4 mutations leading to the gain of a G4.

| GO ID | GO Description | Universe | COSMIC And CLINVAR | Adjusted P-value |
|---|---|---|---|---|
| **GO:0048856** | anatomical structure development | 4152 | 588 | 5.78E-18 |
| **GO:0032502** | developmental process | 4584 | 634 | 2.01E-17 |
| **GO:0048731** | system development | 2976 | 448 | 5.64E-17 |
| **GO:0007275** | multicellular organism development | 3266 | 480 | 1.76E-16 |
| **GO:0007399** | nervous system development | 1648 | 279 | 1.92E-15 |
| **GO:0009653** | anatomical structure morphogenesis | 1871 | 303 | 2.95E-14 |
| **GO:0048869** | cellular developmental process | 2879 | 419 | 7.27E-13 |
| **GO:0030154** | cell differentiation | 2860 | 416 | 1.07E-12 |
| **GO:0007155** | cell adhesion | 1216 | 209 | 2.10E-11 |
| **GO:0032501** | multicellular organismal process | 5319 | 684 | 2.28E-11 |
| **GO:0030029** | actin filament-based process | 717 | 138 | 2.88E-10 |
| **GO:1903508** | positive regulation of nucleic acid-templated transcription | 1297 | 215 | 3.71E-10 |
| **GO:0045893** | positive regulation of DNA-templated transcription | 1297 | 215 | 3.71E-10 |
| **GO:1902680** | positive regulation of RNA biosynthetic process | 1303 | 215 | 5.95E-10 |
| **GO:0045944** | positive regulation of transcription by RNA polymerase II | 952 | 167 | 2.58E-09 |
| **GO:0010646** | regulation of cell communication | 2727 | 384 | 3.03E-09 |
| **GO:0000902** | cell morphogenesis | 718 | 135 | 3.17E-09 |
| **GO:0023051** | regulation of signaling | 2733 | 384 | 4.14E-09 |
| **GO:0050793** | regulation of developmental process | 1810 | 274 | 5.89E-09 |
| **GO:0032989** | cellular component morphogenesis | 549 | 110 | 7.78E-09 |
| **GO:0051254** | positive regulation of RNA metabolic process | 1433 | 227 | 7.96E-09 |
| **GO:0030036** | actin cytoskeleton organization | 637 | 121 | 2.50E-08 |
| **GO:0048858** | cell projection morphogenesis | 459 | 95 | 3.88E-08 |
| **GO:0032990** | cell part morphogenesis | 469 | 96 | 5.73E-08 |
| **GO:0045935** | positive regulation of nucleobase-containing compound metabolic process | 1620 | 245 | 1.21E-07 |
| **GO:0010557** | positive regulation of macromolecule biosynthetic process | 1492 | 229 | 1.43E-07 |
| **GO:0048522** | positive regulation of cellular process | 4704 | 594 | 2.02E-07 |
| **GO:0031328** | positive regulation of cellular biosynthetic process | 1568 | 237 | 2.77E-07 |
| **GO:0048812** | neuron projection morphogenesis | 441 | 90 | 2.81E-07 |
| **GO:0007267** | cell-cell signaling | 1269 | 200 | 2.89E-07 |
| **GO:0120039** | plasma membrane bounded cell projection morphogenesis | 455 | 92 | 2.89E-07 |
| **GO:0048513** | animal organ development | 2176 | 309 | 3.69E-07 |
| **GO:0009891** | positive regulation of biosynthetic process | 1593 | 239 | 4.60E-07 |
| **GO:0009887** | animal organ morphogenesis | 582 | 109 | 6.23E-07 |
| **GO:0023052** | signaling | 5190 | 642 | 8.45E-07 |
| **GO:0048699** | generation of neurons | 969 | 160 | 8.71E-07 |
| **GO:0000904** | cell morphogenesis involved in differentiation | 495 | 96 | 1.14E-06 |
| **GO:0048468** | cell development | 1355 | 208 | 1.18E-06 |
| **GO:0007154** | cell communication | 5221 | 644 | 1.30E-06 |
| **GO:0030182** | neuron differentiation | 923 | 153 | 1.64E-06 |
| **GO:0022008** | neurogenesis | 1096 | 175 | 1.72E-06 |
| **GO:0030030** | cell projection organization | 1164 | 183 | 2.34E-06 |
| **GO:0120036** | plasma membrane bounded cell projection organization | 1144 | 180 | 3.05E-06 |
| **GO:0007010** | cytoskeleton organization | 1309 | 199 | 6.53E-06 |
| **GO:0099536** | synaptic signaling | 522 | 96 | 1.86E-05 |
| **GO:0048667** | cell morphogenesis involved in neuron differentiation | 381 | 76 | 2.12E-05 |
| **GO:0048518** | positive regulation of biological process | 5294 | 642 | 2.88E-05 |
| **GO:0031175** | neuron projection development | 656 | 113 | 3.77E-05 |
| **GO:0050804** | modulation of chemical synaptic transmission | 252 | 56 | 4.32E-05 |
| **GO:0099177** | regulation of trans-synaptic signaling | 253 | 56 | 4.97E-05 |
| **GO:0009966** | regulation of signal transduction | 2464 | 330 | 5.42E-05 |
| **GO:0022603** | regulation of anatomical structure morphogenesis | 700 | 118 | 6.09E-05 |
| **GO:0051173** | positive regulation of nitrogen compound metabolic process | 2565 | 341 | 6.44E-05 |
| **GO:0051239** | regulation of multicellular organismal process | 2117 | 289 | 8.50E-05 |
| **GO:0016043** | cellular component organization | 5370 | 646 | 9.41E-05 |
| **GO:0048666** | neuron development | 729 | 121 | 9.62E-05 |
| **GO:0031325** | positive regulation of cellular metabolic process | 2522 | 335 | 9.64E-05 |
| **GO:0099537** | trans-synaptic signaling | 501 | 90 | 1.58E-04 |
| **GO:0048589** | developmental growth | 273 | 57 | 2.93E-04 |
| **GO:0048523** | negative regulation of cellular process | 3896 | 484 | 2.94E-04 |
| **GO:0048870** | cell motility | 1362 | 197 | 3.32E-04 |
| **GO:0007268** | chemical synaptic transmission | 495 | 88 | 3.49E-04 |
| **GO:0098916** | anterograde trans-synaptic signaling | 495 | 88 | 3.49E-04 |

### Supplemental Table 11 (continued).

| GO ID | GO Description | Universe | COSMIC And CLINVAR | Adjusted P-value |
|---|---|---|---|---|
| **GO:0009987** | cellular process | 14783 | 1556 | 6.39E-04 |
| **GO:0034330** | cell junction organization | 518 | 90 | 7.04E-04 |
| **GO:0009888** | tissue development | 1239 | 180 | 8.89E-04 |
| **GO:0035556** | intracellular signal transduction | 2168 | 288 | 1.05E-03 |
| **GO:0007417** | central nervous system development | 584 | 98 | 1.09E-03 |
| **GO:0010604** | positive regulation of macromolecule metabolic process | 2889 | 369 | 1.09E-03 |
| **GO:0045595** | regulation of cell differentiation | 1129 | 166 | 1.22E-03 |
| **GO:0051128** | regulation of cellular component organization | 1929 | 260 | 1.31E-03 |
| **GO:0099587** | inorganic ion import across plasma membrane | 112 | 30 | 1.40E-03 |
| **GO:0098659** | inorganic cation import across plasma membrane | 112 | 30 | 1.40E-03 |
| **GO:0060322** | head development | 402 | 73 | 1.69E-03 |
| **GO:0071840** | cellular component organization or biogenesis | 5539 | 653 | 1.79E-03 |
| **GO:0048519** | negative regulation of biological process | 4381 | 529 | 2.36E-03 |
| **GO:0007409** | axonogenesis | 298 | 58 | 2.40E-03 |
| **GO:0009893** | positive regulation of metabolic process | 3177 | 398 | 2.59E-03 |
| **GO:0016477** | cell migration | 1210 | 174 | 2.61E-03 |
| **GO:0051716** | cellular response to stimulus | 5982 | 697 | 3.27E-03 |
| **GO:0060560** | developmental growth involved in morphogenesis | 135 | 33 | 3.51E-03 |
| **GO:0007420** | brain development | 384 | 69 | 4.70E-03 |
| **GO:0048729** | tissue morphogenesis | 348 | 64 | 4.93E-03 |
| **GO:0007165** | signal transduction | 4776 | 568 | 5.32E-03 |
| **GO:0061564** | axon development | 323 | 60 | 7.13E-03 |
| **GO:0014074** | response to purine-containing compound | 75 | 22 | 7.97E-03 |
| **GO:0048670** | regulation of collateral sprouting | 9 | 7 | 8.28E-03 |
| **GO:0048588** | developmental cell growth | 129 | 31 | 9.81E-03 |
| **GO:1990138** | neuron projection extension | 112 | 28 | 1.23E-02 |
| **GO:0050794** | regulation of cellular process | 9523 | 1052 | 1.23E-02 |
| **GO:0007167** | enzyme-linked receptor protein signaling pathway | 795 | 120 | 1.26E-02 |
| **GO:0098657** | import into cell | 216 | 44 | 1.26E-02 |
| **GO:0009719** | response to endogenous stimulus | 1076 | 154 | 1.31E-02 |
| **GO:0032970** | regulation of actin filament-based process | 330 | 60 | 1.36E-02 |
| **GO:0031344** | regulation of cell projection organization | 467 | 78 | 1.63E-02 |
| **GO:0071495** | cellular response to endogenous stimulus | 974 | 141 | 1.80E-02 |
| **GO:0120035** | regulation of plasma membrane bounded cell projection organization | 453 | 76 | 1.80E-02 |
| **GO:0098739** | import across plasma membrane | 172 | 37 | 1.85E-02 |
| **GO:0034329** | cell junction assembly | 341 | 61 | 1.88E-02 |
| **GO:0051094** | positive regulation of developmental process | 967 | 140 | 1.91E-02 |
| **GO:0048583** | regulation of response to stimulus | 3327 | 407 | 1.94E-02 |
| **GO:0007156** | homophilic cell adhesion via plasma membrane adhesion molecules | 63 | 19 | 2.15E-02 |
| **GO:0007015** | actin filament organization | 395 | 68 | 2.20E-02 |
| **GO:0032535** | regulation of cellular component size | 256 | 49 | 2.21E-02 |
| **GO:0048638** | regulation of developmental growth | 167 | 36 | 2.28E-02 |
| **GO:0007169** | transmembrane receptor protein tyrosine kinase signaling pathway | 511 | 83 | 2.39E-02 |
| **GO:0098742** | cell-cell adhesion via plasma-membrane adhesion molecules | 195 | 40 | 2.62E-02 |
| **GO:0051093** | negative regulation of developmental process | 619 | 96 | 3.43E-02 |
| **GO:0050789** | regulation of biological process | 10085 | 1103 | 3.55E-02 |
| **GO:0010243** | response to organonitrogen compound | 613 | 95 | 3.86E-02 |
| **GO:0098609** | cell-cell adhesion | 743 | 111 | 4.07E-02 |
| **GO:0048639** | positive regulation of developmental growth | 83 | 22 | 4.10E-02 |
| **GO:1905114** | cell surface receptor signaling pathway involved in cell-cell signaling | 388 | 66 | 4.13E-02 |
| **GO:0051172** | negative regulation of nitrogen compound metabolic process | 1927 | 249 | 4.44E-02 |
| **GO:0072359** | circulatory system development | 729 | 109 | 4.60E-02 |

**Supplemental Table 12.** Significant GO:BP enrichments for COSMIC G4 mutations leading to the gain of a G4.

| GO ID | GO Description | Universe | COSMIC | Adjusted P-value |
|---|---|---|---|---|
| **GO:1903508** | positive regulation of nucleic acid-templated transcription | 1297 | 121 | 2.51E-05 |
| **GO:0045893** | positive regulation of DNA-templated transcription | 1297 | 121 | 2.51E-05 |
| **GO:1902680** | positive regulation of RNA biosynthetic process | 1303 | 121 | 3.28E-05 |
| **GO:0051254** | positive regulation of RNA metabolic process | 1433 | 129 | 6.57E-05 |
| **GO:0007155** | cell adhesion | 1216 | 111 | 0.00034137 |
| **GO:0045944** | positive regulation of transcription by RNA polymerase II | 952 | 92 | 0.00035644 |
| **GO:0045935** | positive regulation of nucleobase-containing compound metabolic process | 1620 | 137 | 0.00095303 |
| **GO:0046907** | intracellular transport | 1373 | 118 | 0.00307086 |
| **GO:0010557** | positive regulation of macromolecule biosynthetic process | 1492 | 126 | 0.00321361 |
| **GO:0031328** | positive regulation of cellular biosynthetic process | 1568 | 130 | 0.00562818 |
| **GO:0048856** | anatomical structure development | 4152 | 292 | 0.00721462 |
| **GO:0048522** | positive regulation of cellular process | 4704 | 325 | 0.00728079 |
| **GO:0009891** | positive regulation of biosynthetic process | 1593 | 130 | 0.01262319 |
| **GO:0010604** | positive regulation of macromolecule metabolic process | 2889 | 213 | 0.01326194 |
| **GO:0007275** | multicellular organism development | 3266 | 236 | 0.01514072 |
| **GO:0048731** | system development | 2976 | 218 | 0.01552671 |
| **GO:0051649** | establishment of localization in cell | 1698 | 136 | 0.01953263 |
| **GO:0009653** | anatomical structure morphogenesis | 1871 | 147 | 0.02240799 |
| **GO:0051173** | positive regulation of nitrogen compound metabolic process | 2565 | 191 | 0.02564038 |
| **GO:0032502** | developmental process | 4584 | 314 | 0.02643181 |
| **GO:0030036** | actin cytoskeleton organization | 637 | 62 | 0.03246111 |
| **GO:0051641** | cellular localization | 2721 | 200 | 0.03443678 |

**Supplemental Table 13.** Significant GO:BP enrichments for CLINVAR G4 mutations leading to the gain of a G4.

| GO ID | GO Description | Universe | CLINVAR | Adjusted P-value |
|---|---|---|---|---|
| **GO:0001508** | action potential | 117 | 6 | 4.30E-02 |
| **GO:0086001** | cardiac muscle cell action potential | 73 | 5 | 4.96E-02 |

**Supplemental Table 14.** Significant GO:CC enrichments for COSMIC and CLINVAR G4 mutations.

| GO ID | GO Description | Universe | COSMIC and CLINVAR | Adjusted P-value |
|---|---|---|---|---|
| **GO:0030054** | cell junction | 1385 | 893 | 2.16E-34 |
| **GO:0071944** | cell periphery | 5199 | 2884 | 2.79E-34 |
| **GO:0042995** | cell projection | 1567 | 979 | 3.89E-30 |
| **GO:0005886** | plasma membrane | 4762 | 2634 | 7.87E-29 |
| **GO:0120025** | plasma membrane bounded cell projection | 1540 | 959 | 1.11E-28 |
| **GO:0098590** | plasma membrane region | 794 | 529 | 4.09E-24 |
| **GO:0043005** | neuron projection | 839 | 553 | 1.42E-23 |
| **GO:0045202** | synapse | 652 | 445 | 2.85E-23 |
| **GO:0016020** | membrane | 7611 | 3986 | 9.36E-21 |
| **GO:0098797** | plasma membrane protein complex | 540 | 364 | 2.72E-17 |
| **GO:0030424** | axon | 339 | 243 | 4.48E-16 |
| **GO:0031226** | intrinsic component of plasma membrane | 1632 | 951 | 2.93E-15 |
| **GO:0005887** | integral component of plasma membrane | 1560 | 913 | 2.97E-15 |
| **GO:0015629** | actin cytoskeleton | 441 | 300 | 7.75E-15 |
| **GO:0036477** | somatodendritic compartment | 444 | 300 | 3.31E-14 |
| **GO:0005856** | cytoskeleton | 1812 | 1033 | 5.15E-13 |
| **GO:0031252** | cell leading edge | 289 | 206 | 5.23E-13 |
| **GO:0097060** | synaptic membrane | 160 | 126 | 9.52E-13 |
| **GO:0070161** | anchoring junction | 707 | 442 | 2.07E-12 |
| **GO:0043235** | receptor complex | 395 | 266 | 3.53E-12 |
| **GO:0098794** | postsynapse | 293 | 204 | 3.32E-11 |
| **GO:0005911** | cell-cell junction | 369 | 247 | 8.94E-11 |
| **GO:0097447** | dendritic tree | 341 | 230 | 1.79E-10 |
| **GO:0030425** | dendrite | 338 | 228 | 2.23E-10 |
| **GO:0005737** | cytoplasm | 10633 | 5335 | 1.52E-09 |
| **GO:0005912** | adherens junction | 144 | 110 | 1.96E-09 |
| **GO:0043228** | non-membrane-bounded organelle | 4394 | 2308 | 4.06E-09 |
| **GO:0043232** | intracellular non-membrane-bounded organelle | 4393 | 2307 | 4.57E-09 |
| **GO:0031224** | intrinsic component of membrane | 2600 | 1406 | 1.40E-08 |
| **GO:0005938** | cell cortex | 209 | 147 | 2.90E-08 |
| **GO:0099572** | postsynaptic specialization | 163 | 119 | 4.37E-08 |
| **GO:0005829** | cytosol | 5137 | 2662 | 6.95E-08 |
| **GO:0099081** | supramolecular polymer | 659 | 396 | 1.20E-07 |
| **GO:0098793** | presynapse | 241 | 164 | 1.37E-07 |
| **GO:0016021** | integral component of membrane | 2485 | 1339 | 1.60E-07 |
| **GO:0014069** | postsynaptic density | 151 | 110 | 2.64E-07 |
| **GO:0045211** | postsynaptic membrane | 106 | 82 | 3.26E-07 |
| **GO:0098984** | neuron to neuron synapse | 163 | 117 | 3.38E-07 |
| **GO:0099512** | supramolecular fiber | 650 | 389 | 3.45E-07 |
| **GO:0031982** | vesicle | 3535 | 1861 | 4.51E-07 |
| **GO:0042383** | sarcolemma | 77 | 63 | 5.11E-07 |
| **GO:0098978** | glutamatergic synapse | 82 | 66 | 7.67E-07 |
| **GO:0012505** | endomembrane system | 3988 | 2083 | 8.67E-07 |
| **GO:0032279** | asymmetric synapse | 158 | 113 | 9.15E-07 |
| **GO:0043025** | neuronal cell body | 209 | 143 | 1.00E-06 |
| **GO:0098588** | bounding membrane of organelle | 1644 | 905 | 1.02E-06 |
| **GO:0031253** | cell projection membrane | 215 | 146 | 1.50E-06 |
| **GO:0097708** | intracellular vesicle | 1967 | 1068 | 1.67E-06 |
| **GO:0031410** | cytoplasmic vesicle | 1966 | 1067 | 1.92E-06 |
| **GO:0043226** | organelle | 12740 | 6294 | 1.95E-06 |
| **GO:0019898** | extrinsic component of membrane | 282 | 184 | 1.95E-06 |
| **GO:0034702** | ion channel complex | 247 | 164 | 2.08E-06 |
| **GO:0044297** | cell body | 237 | 158 | 2.57E-06 |
| **GO:0034703** | cation channel complex | 218 | 147 | 2.71E-06 |
| **GO:0150034** | distal axon | 144 | 103 | 4.39E-06 |
| **GO:0005884** | actin filament | 105 | 79 | 5.83E-06 |
| **GO:0012506** | vesicle membrane | 881 | 504 | 8.55E-06 |
| **GO:0042734** | presynaptic membrane | 53 | 45 | 1.25E-05 |
| **GO:0030027** | lamellipodium | 135 | 96 | 2.09E-05 |
| **GO:0045177** | apical part of cell | 267 | 172 | 2.18E-05 |
| **GO:0005622** | intracellular anatomical structure | 13666 | 6714 | 2.42E-05 |
| **GO:0030659** | cytoplasmic vesicle membrane | 868 | 494 | 2.79E-05 |
| **GO:0019897** | extrinsic component of plasma membrane | 156 | 108 | 3.06E-05 |
| **GO:0099080** | supramolecular complex | 978 | 549 | 6.16E-05 |

**Supplemental Table 14 (continued).**

| GO ID | GO Description | Universe | COSMIC and CLINVAR | Adjusted P-value |
|---|---|---|---|---|
| **GO:0009925** | basal plasma membrane | 177 | 119 | 8.22E-05 |
| **GO:0045178** | basal part of cell | 184 | 123 | 8.39E-05 |
| **GO:0042641** | actomyosin | 64 | 51 | 8.50E-05 |
| **GO:0031256** | leading edge membrane | 111 | 80 | 1.00E-04 |
| **GO:0098802** | plasma membrane signaling receptor complex | 176 | 118 | 1.14E-04 |
| **GO:0030863** | cortical cytoskeleton | 82 | 62 | 1.36E-04 |
| **GO:0044304** | main axon | 39 | 34 | 1.57E-04 |
| **GO:0048786** | presynaptic active zone | 39 | 34 | 1.57E-04 |
| **GO:0099634** | postsynaptic specialization membrane | 45 | 38 | 1.83E-04 |
| **GO:0016323** | basolateral plasma membrane | 158 | 107 | 1.85E-04 |
| **GO:0008328** | ionotropic glutamate receptor complex | 32 | 29 | 1.85E-04 |
| **GO:0098839** | postsynaptic density membrane | 35 | 31 | 2.38E-04 |
| **GO:0030055** | cell-substrate junction | 395 | 238 | 2.60E-04 |
| **GO:0030426** | growth cone | 88 | 65 | 2.96E-04 |
| **GO:0030427** | site of polarized growth | 93 | 68 | 3.11E-04 |
| **GO:0099513** | polymeric cytoskeletal fiber | 482 | 284 | 3.57E-04 |
| **GO:0005901** | caveola | 61 | 48 | 3.61E-04 |
| **GO:0005925** | focal adhesion | 387 | 233 | 3.62E-04 |
| **GO:0098796** | membrane protein complex | 1115 | 614 | 3.62E-04 |
| **GO:0016324** | apical plasma membrane | 234 | 149 | 4.01E-04 |
| **GO:0097517** | contractile actin filament bundle | 58 | 46 | 4.01E-04 |
| **GO:0001725** | stress fiber | 58 | 46 | 4.01E-04 |
| **GO:0000785** | chromatin | 1214 | 663 | 5.89E-04 |
| **GO:0098858** | actin-based cell projection | 133 | 91 | 6.17E-04 |
| **GO:0005604** | basement membrane | 65 | 50 | 7.08E-04 |
| **GO:0009898** | cytoplasmic side of plasma membrane | 160 | 106 | 9.53E-04 |
| **GO:0032432** | actin filament bundle | 64 | 49 | 1.11E-03 |
| **GO:0043292** | contractile fiber | 173 | 113 | 1.26E-03 |
| **GO:0005794** | Golgi apparatus | 1365 | 737 | 1.26E-03 |
| **GO:0045121** | membrane raft | 209 | 133 | 1.45E-03 |
| **GO:0005768** | endosome | 810 | 452 | 1.57E-03 |
| **GO:0030016** | myofibril | 165 | 108 | 1.74E-03 |
| **GO:0098862** | cluster of actin-based cell projections | 96 | 68 | 1.83E-03 |
| **GO:0098857** | membrane microdomain | 210 | 133 | 2.05E-03 |
| **GO:0044309** | neuron spine | 90 | 64 | 2.72E-03 |
| **GO:0044853** | plasma membrane raft | 87 | 62 | 3.30E-03 |
| **GO:0070382** | exocytic vesicle | 151 | 99 | 3.62E-03 |
| **GO:0043197** | dendritic spine | 89 | 63 | 3.94E-03 |
| **GO:0030017** | sarcomere | 146 | 96 | 4.06E-03 |
| **GO:0043229** | intracellular organelle | 11898 | 5852 | 4.14E-03 |
| **GO:0034705** | potassium channel complex | 91 | 64 | 4.68E-03 |
| **GO:0044291** | cell-cell contact zone | 50 | 39 | 4.80E-03 |
| **GO:0048471** | perinuclear region of cytoplasm | 440 | 255 | 5.80E-03 |
| **GO:0032589** | neuron projection membrane | 38 | 31 | 7.16E-03 |
| **GO:0031674** | I band | 97 | 67 | 7.44E-03 |
| **GO:0016342** | catenin complex | 32 | 27 | 7.57E-03 |
| **GO:0005769** | early endosome | 295 | 177 | 7.71E-03 |
| **GO:0030864** | cortical actin cytoskeleton | 62 | 46 | 8.23E-03 |
| **GO:0001726** | ruffle | 118 | 79 | 8.51E-03 |
| **GO:1902495** | transmembrane transporter complex | 331 | 196 | 8.99E-03 |
| **GO:0098878** | neurotransmitter receptor complex | 37 | 30 | 1.18E-02 |
| **GO:0030018** | Z disc | 86 | 60 | 1.19E-02 |
| **GO:0032420** | stereocilium | 31 | 26 | 1.29E-02 |
| **GO:0008021** | synaptic vesicle | 128 | 84 | 1.42E-02 |
| **GO:0031234** | extrinsic component of cytoplasmic side of plasma membrane | 95 | 65 | 1.48E-02 |
| **GO:0015630** | microtubule cytoskeleton | 1159 | 622 | 1.58E-02 |
| **GO:0030315** | T-tubule | 36 | 29 | 1.92E-02 |
| **GO:0016010** | dystrophin-associated glycoprotein complex | 17 | 16 | 2.25E-02 |
| **GO:0032281** | AMPA glutamate receptor complex | 20 | 18 | 3.07E-02 |
| **GO:0031941** | filamentous actin | 29 | 24 | 3.62E-02 |
| **GO:0005815** | microtubule organizing center | 724 | 397 | 3.66E-02 |
| **GO:0043194** | axon initial segment | 12 | 12 | 4.29E-02 |
| **GO:0008076** | voltage-gated potassium channel complex | 82 | 56 | 4.75E-02 |
| **GO:0034399** | nuclear periphery | 96 | 64 | 5.00E-02 |

**Supplemental Table 15.** Significant GO:CC enrichments for COSMIC G4 mutations.

| GO ID | GO Description | Universe | COSMIC | Adjusted P-value |
|---|---|---|---|---|
| **GO:0071944** | cell periphery | 5199 | 2843 | 5.77E-34 |
| **GO:0030054** | cell junction | 1385 | 879 | 3.22E-33 |
| **GO:0042995** | cell projection | 1567 | 967 | 4.83E-30 |
| **GO:0005886** | plasma membrane | 4762 | 2598 | 7.93E-29 |
| **GO:0120025** | plasma membrane bounded cell projection | 1540 | 947 | 1.55E-28 |
| **GO:0045202** | synapse | 652 | 440 | 3.61E-23 |
| **GO:0098590** | plasma membrane region | 794 | 520 | 5.03E-23 |
| **GO:0043005** | neuron projection | 839 | 545 | 5.77E-23 |
| **GO:0016020** | membrane | 7611 | 3925 | 2.66E-20 |
| **GO:0030424** | axon | 339 | 242 | 1.07E-16 |
| **GO:0098797** | plasma membrane protein complex | 540 | 358 | 1.46E-16 |
| **GO:0005887** | integral component of plasma membrane | 1560 | 900 | 5.34E-15 |
| **GO:0015629** | actin cytoskeleton | 441 | 297 | 6.5E-15 |
| **GO:0031226** | intrinsic component of plasma membrane | 1632 | 936 | 9.92E-15 |
| **GO:0036477** | somatodendritic compartment | 444 | 296 | 6.2E-14 |
| **GO:0097060** | synaptic membrane | 160 | 126 | 2.17E-13 |
| **GO:0031252** | cell leading edge | 289 | 204 | 4.97E-13 |
| **GO:0005856** | cytoskeleton | 1812 | 1018 | 8.96E-13 |
| **GO:0070161** | anchoring junction | 707 | 436 | 3.3E-12 |
| **GO:0098794** | postsynapse | 293 | 202 | 3.02E-11 |
| **GO:0043235** | receptor complex | 395 | 259 | 8.91E-11 |
| **GO:0097447** | dendritic tree | 341 | 227 | 2.74E-10 |
| **GO:0030425** | dendrite | 338 | 225 | 3.47E-10 |
| **GO:0005911** | cell-cell junction | 369 | 242 | 5.32E-10 |
| **GO:0005912** | adherens junction | 144 | 110 | 5.79E-10 |
| **GO:0043228** | non-membrane-bounded organelle | 4394 | 2279 | 1.51E-09 |
| **GO:0043232** | intracellular non-membrane-bounded organelle | 4393 | 2278 | 1.7E-09 |
| **GO:0005737** | cytoplasm | 10633 | 5250 | 4.16E-09 |
| **GO:0099572** | postsynaptic specialization | 163 | 119 | 1.28E-08 |
| **GO:0005938** | cell cortex | 209 | 146 | 1.83E-08 |
| **GO:0031224** | intrinsic component of membrane | 2600 | 1385 | 2.07E-08 |
| **GO:0014069** | postsynaptic density | 151 | 110 | 8.66E-08 |
| **GO:0016021** | integral component of membrane | 2485 | 1322 | 1.01E-07 |
| **GO:0045211** | postsynaptic membrane | 106 | 82 | 1.31E-07 |
| **GO:0005829** | cytosol | 5137 | 2619 | 1.67E-07 |
| **GO:0098793** | presynapse | 241 | 162 | 1.7E-07 |
| **GO:0098984** | neuron to neuron synapse | 163 | 116 | 2.92E-07 |
| **GO:0099081** | supramolecular polymer | 659 | 389 | 3.35E-07 |
| **GO:0098978** | glutamatergic synapse | 82 | 66 | 3.5E-07 |
| **GO:0019898** | extrinsic component of membrane | 282 | 184 | 4.45E-07 |
| **GO:0034702** | ion channel complex | 247 | 164 | 5.27E-07 |
| **GO:0034703** | cation channel complex | 218 | 147 | 7.57E-07 |
| **GO:0032279** | asymmetric synapse | 158 | 112 | 8.12E-07 |
| **GO:0099512** | supramolecular fiber | 650 | 382 | 9.7E-07 |
| **GO:0042383** | sarcolemma | 77 | 62 | 1.08E-06 |
| **GO:0043025** | neuronal cell body | 209 | 141 | 1.5E-06 |
| **GO:0043226** | organelle | 12740 | 6199 | 1.51E-06 |
| **GO:0012505** | endomembrane system | 3988 | 2050 | 1.58E-06 |
| **GO:0031253** | cell projection membrane | 215 | 144 | 2.15E-06 |
| **GO:0005884** | actin filament | 105 | 79 | 2.5E-06 |
| **GO:0098588** | bounding membrane of organelle | 1644 | 889 | 3E-06 |
| **GO:0044297** | cell body | 237 | 156 | 3.16E-06 |
| **GO:0031982** | vesicle | 3535 | 1825 | 3.48E-06 |
| **GO:0097708** | intracellular vesicle | 1967 | 1050 | 3.97E-06 |
| **GO:0150034** | distal axon | 144 | 102 | 4.31E-06 |
| **GO:0031410** | cytoplasmic vesicle | 1966 | 1049 | 4.54E-06 |
| **GO:0042734** | presynaptic membrane | 53 | 45 | 7.12E-06 |
| **GO:0012506** | vesicle membrane | 881 | 497 | 9.91E-06 |
| **GO:0019897** | extrinsic component of plasma membrane | 156 | 108 | 1.15E-05 |
| **GO:0030027** | lamellipodium | 135 | 95 | 2.2E-05 |
| **GO:0005622** | intracellular anatomical structure | 13666 | 6611 | 2.35E-05 |
| **GO:0030659** | cytoplasmic vesicle membrane | 868 | 487 | 3.36E-05 |
| **GO:0042641** | actomyosin | 64 | 51 | 4.76E-05 |
| **GO:0099080** | supramolecular complex | 978 | 542 | 5.51E-05 |
| **GO:0030863** | cortical cytoskeleton | 82 | 62 | 7.1E-05 |

**Supplemental Table 15 (continued).**

| GO ID | GO Description | Universe | COSMIC | Adjusted P-value |
|---|---|---|---|---|
| **GO:0048786** | presynaptic active zone | 39 | 34 | 0.000103 |
| **GO:0044304** | main axon | 39 | 34 | 0.000103 |
| **GO:0099634** | postsynaptic specialization membrane | 45 | 38 | 0.000116 |
| **GO:0031256** | leading edge membrane | 111 | 79 | 0.000126 |
| **GO:0008328** | ionotropic glutamate receptor complex | 32 | 29 | 0.000128 |
| **GO:0045177** | apical part of cell | 267 | 167 | 0.000152 |
| **GO:0030426** | growth cone | 88 | 65 | 0.000155 |
| **GO:0030427** | site of polarized growth | 93 | 68 | 0.000159 |
| **GO:0098839** | postsynaptic density membrane | 35 | 31 | 0.000161 |
| **GO:0001725** | stress fiber | 58 | 46 | 0.000242 |
| **GO:0097517** | contractile actin filament bundle | 58 | 46 | 0.000242 |
| **GO:0030055** | cell-substrate junction | 395 | 235 | 0.000271 |
| **GO:0045178** | basal part of cell | 184 | 120 | 0.000291 |
| **GO:0009925** | basal plasma membrane | 177 | 116 | 0.000304 |
| **GO:0098796** | membrane protein complex | 1115 | 606 | 0.000328 |
| **GO:0005925** | focal adhesion | 387 | 230 | 0.000389 |
| **GO:0009898** | cytoplasmic side of plasma membrane | 160 | 106 | 0.000405 |
| **GO:0005604** | basement membrane | 65 | 50 | 0.000416 |
| **GO:0098802** | plasma membrane signaling receptor complex | 176 | 115 | 0.000416 |
| **GO:0098858** | actin-based cell projection | 133 | 90 | 0.000646 |
| **GO:0099513** | polymeric cytoskeletal fiber | 482 | 279 | 0.000661 |
| **GO:0032432** | actin filament bundle | 64 | 49 | 0.000663 |
| **GO:0000785** | chromatin | 1214 | 653 | 0.000752 |
| **GO:0016323** | basolateral plasma membrane | 158 | 104 | 0.000776 |
| **GO:0005901** | caveola | 61 | 47 | 0.000777 |
| **GO:0016324** | apical plasma membrane | 234 | 146 | 0.000888 |
| **GO:0043292** | contractile fiber | 173 | 112 | 0.001057 |
| **GO:0005768** | endosome | 810 | 447 | 0.001123 |
| **GO:0030016** | myofibril | 165 | 107 | 0.001521 |
| **GO:0070382** | exocytic vesicle | 151 | 99 | 0.001679 |
| **GO:0098862** | cluster of actin-based cell projections | 96 | 67 | 0.002472 |
| **GO:0034705** | potassium channel complex | 91 | 64 | 0.002626 |
| **GO:0043229** | intracellular organelle | 11898 | 5765 | 0.002825 |
| **GO:1902495** | transmembrane transporter complex | 331 | 196 | 0.002936 |
| **GO:0045121** | membrane raft | 209 | 130 | 0.003563 |
| **GO:0030017** | sarcomere | 146 | 95 | 0.003962 |
| **GO:0005769** | early endosome | 295 | 176 | 0.004405 |
| **GO:0001726** | ruffle | 118 | 79 | 0.004484 |
| **GO:0044853** | plasma membrane raft | 87 | 61 | 0.004808 |
| **GO:0098857** | membrane microdomain | 210 | 130 | 0.00492 |
| **GO:0032589** | neuron projection membrane | 38 | 31 | 0.005071 |
| **GO:0030864** | cortical actin cytoskeleton | 62 | 46 | 0.005237 |
| **GO:0016342** | catenin complex | 32 | 27 | 0.005544 |
| **GO:0005794** | Golgi apparatus | 1365 | 720 | 0.006418 |
| **GO:0008021** | synaptic vesicle | 128 | 84 | 0.007447 |
| **GO:0098878** | neurotransmitter receptor complex | 37 | 30 | 0.008507 |
| **GO:0031234** | extrinsic component of cytoplasmic side of plasma membrane | 95 | 65 | 0.008596 |
| **GO:0044309** | neuron spine | 90 | 62 | 0.009397 |
| **GO:0031674** | I band | 97 | 66 | 0.009766 |
| **GO:0048471** | perinuclear region of cytoplasm | 440 | 250 | 0.011477 |
| **GO:0043197** | dendritic spine | 89 | 61 | 0.013481 |
| **GO:0030315** | T-tubule | 36 | 29 | 0.014156 |
| **GO:0030018** | Z disc | 86 | 59 | 0.016795 |
| **GO:0032281** | AMPA glutamate receptor complex | 20 | 18 | 0.02496 |
| **GO:0015630** | microtubule cytoskeleton | 1159 | 611 | 0.027121 |
| **GO:1990351** | transporter complex | 356 | 204 | 0.02757 |
| **GO:0031941** | filamentous actin | 29 | 24 | 0.027799 |
| **GO:0008076** | voltage-gated potassium channel complex | 82 | 56 | 0.029877 |
| **GO:0044291** | cell-cell contact zone | 50 | 37 | 0.033809 |
| **GO:0043194** | axon initial segment | 12 | 12 | 0.036977 |
| **GO:0016363** | nuclear matrix | 79 | 54 | 0.037414 |
| **GO:0099240** | intrinsic component of synaptic membrane | 34 | 27 | 0.037971 |
| **GO:0032420** | stereocilium | 31 | 25 | 0.042583 |
| **GO:0005815** | microtubule organizing center | 724 | 391 | 0.043147 |

**Supplemental Table 16.** Significant GO:CC enrichments for CLINVAR G4 mutations.

| GO ID | GO Description | Universe | CLINVAR | Adjusted P-value |
|---|---|---|---|---|
| **GO:0030017** | sarcomere | 146 | 26 | 9.1E-08 |
| **GO:0031674** | I band | 97 | 16 | 5.6E-04 |
| **GO:0042383** | sarcolemma | 77 | 14 | 7.7E-04 |
| **GO:0036379** | myofilament | 25 | 8 | 1.6E-03 |
| **GO:0030315** | T-tubule | 36 | 9 | 3.3E-03 |
| **GO:0005865** | striated muscle thin filament | 21 | 7 | 4.4E-03 |
| **GO:0014704** | intercalated disc | 32 | 8 | 9.8E-03 |
| **GO:0030018** | Z disc | 86 | 13 | 1.1E-02 |
| **GO:0043202** | lysosomal lumen | 87 | 13 | 1.2E-02 |
| **GO:0033268** | node of Ranvier | 12 | 5 | 2.1E-02 |
| **GO:0043194** | axon initial segment | 12 | 5 | 2.1E-02 |
| **GO:1990584** | cardiac Troponin complex | 3 | 3 | 2.2E-02 |
| **GO:0044291** | cell-cell contact zone | 50 | 9 | 4.0E-02 |
| **GO:0005861** | troponin complex | 8 | 4 | 4.8E-02 |

**Supplemental Table 17.** Significant KEGG enrichments for COSMIC and CLINVAR G4 mutations.

| KEGG ID | KEGG Description | Universe | COSMIC and CLINVAR | Adjusted P-value |
|---|---|---|---|---|
| **KEGG:04360** | Axon guidance | 181 | 137 | 9.52E-13 |
| **KEGG:04921** | Oxytocin signaling pathway | 154 | 120 | 1.19E-12 |
| **KEGG:05412** | Arrhythmogenic right ventricular cardiomyopathy | 77 | 68 | 1.05E-11 |
| **KEGG:04010** | MAPK signaling pathway | 294 | 201 | 2.43E-11 |
| **KEGG:04724** | Glutamatergic synapse | 114 | 92 | 4.31E-11 |
| **KEGG:04015** | Rap1 signaling pathway | 210 | 151 | 5.83E-11 |
| **KEGG:05200** | Pathways in cancer | 529 | 330 | 1.01E-10 |
| **KEGG:04510** | Focal adhesion | 200 | 143 | 5.00E-10 |
| **KEGG:04929** | GnRH secretion | 64 | 57 | 5.44E-10 |
| **KEGG:04728** | Dopaminergic synapse | 132 | 100 | 4.36E-09 |
| **KEGG:04261** | Adrenergic signaling in cardiomyocytes | 150 | 111 | 4.46E-09 |
| **KEGG:04072** | Phospholipase D signaling pathway | 147 | 109 | 5.47E-09 |
| **KEGG:04014** | Ras signaling pathway | 234 | 159 | 1.91E-08 |
| **KEGG:01522** | Endocrine resistance | 95 | 75 | 4.97E-08 |
| **KEGG:04725** | Cholinergic synapse | 113 | 86 | 7.34E-08 |
| **KEGG:04020** | Calcium signaling pathway | 239 | 160 | 9.32E-08 |
| **KEGG:04912** | GnRH signaling pathway | 93 | 73 | 1.37E-07 |
| **KEGG:05414** | Dilated cardiomyopathy | 95 | 74 | 1.96E-07 |
| **KEGG:04713** | Circadian entrainment | 97 | 75 | 2.75E-07 |
| **KEGG:04720** | Long-term potentiation | 67 | 55 | 9.00E-07 |
| **KEGG:05215** | Prostate cancer | 97 | 74 | 9.96E-07 |
| **KEGG:04934** | Cushing syndrome | 153 | 107 | 1.96E-06 |
| **KEGG:04750** | Inflammatory mediator regulation of TRP channels | 98 | 74 | 2.12E-06 |
| **KEGG:05410** | Hypertrophic cardiomyopathy | 90 | 69 | 2.26E-06 |
| **KEGG:04810** | Regulation of actin cytoskeleton | 216 | 143 | 2.26E-06 |
| **KEGG:04012** | ErbB signaling pathway | 84 | 65 | 3.13E-06 |
| **KEGG:04925** | Aldosterone synthesis and secretion | 98 | 73 | 6.95E-06 |
| **KEGG:04151** | PI3K-Akt signaling pathway | 353 | 216 | 1.11E-05 |
| **KEGG:04722** | Neurotrophin signaling pathway | 119 | 85 | 1.35E-05 |
| **KEGG:04911** | Insulin secretion | 86 | 65 | 1.49E-05 |
| **KEGG:04022** | cGMP-PKG signaling pathway | 166 | 112 | 1.84E-05 |
| **KEGG:05213** | Endometrial cancer | 58 | 47 | 2.40E-05 |
| **KEGG:04370** | VEGF signaling pathway | 59 | 47 | 6.10E-05 |
| **KEGG:04730** | Long-term depression | 59 | 47 | 6.10E-05 |
| **KEGG:04660** | T cell receptor signaling pathway | 103 | 74 | 6.17E-05 |
| **KEGG:05214** | Glioma | 75 | 57 | 6.46E-05 |
| **KEGG:04927** | Cortisol synthesis and secretion | 64 | 50 | 7.72E-05 |
| **KEGG:04070** | Phosphatidylinositol signaling system | 97 | 70 | 9.54E-05 |
| **KEGG:04919** | Thyroid hormone signaling pathway | 121 | 84 | 1.09E-04 |
| **KEGG:04662** | B cell receptor signaling pathway | 79 | 59 | 1.11E-04 |
| **KEGG:01521** | EGFR tyrosine kinase inhibitor resistance | 79 | 59 | 1.11E-04 |
| **KEGG:05165** | Human papillomavirus infection | 331 | 200 | 1.17E-04 |
| **KEGG:04928** | Parathyroid hormone synthesis, secretion and action | 106 | 75 | 1.33E-04 |
| **KEGG:05166** | Human T-cell leukemia virus 1 infection | 219 | 139 | 1.37E-04 |
| **KEGG:04330** | Notch signaling pathway | 59 | 46 | 2.38E-04 |
| **KEGG:05224** | Breast cancer | 147 | 98 | 2.42E-04 |
| **KEGG:05031** | Amphetamine addiction | 69 | 52 | 3.10E-04 |
| **KEGG:04727** | GABAergic synapse | 89 | 64 | 3.38E-04 |
| **KEGG:04380** | Osteoclast differentiation | 125 | 85 | 3.47E-04 |
| **KEGG:04390** | Hippo signaling pathway | 157 | 103 | 3.97E-04 |
| **KEGG:05225** | Hepatocellular carcinoma | 166 | 108 | 4.11E-04 |
| **KEGG:05205** | Proteoglycans in cancer | 205 | 129 | 6.14E-04 |
| **KEGG:05222** | Small cell lung cancer | 92 | 65 | 7.28E-04 |
| **KEGG:04935** | Growth hormone synthesis, secretion and action | 120 | 81 | 8.95E-04 |
| **KEGG:04658** | Th1 and Th2 cell differentiation | 89 | 63 | 9.14E-04 |
| **KEGG:04666** | Fc gamma R-mediated phagocytosis | 96 | 67 | 1.01E-03 |
| **KEGG:04611** | Platelet activation | 124 | 83 | 1.13E-03 |
| **KEGG:04971** | Gastric acid secretion | 76 | 55 | 1.22E-03 |
| **KEGG:05220** | Chronic myeloid leukemia | 76 | 55 | 1.22E-03 |
| **KEGG:04512** | ECM-receptor interaction | 88 | 62 | 1.36E-03 |
| **KEGG:05032** | Morphine addiction | 90 | 63 | 1.60E-03 |
| **KEGG:04310** | Wnt signaling pathway | 170 | 108 | 1.92E-03 |
| **KEGG:04926** | Relaxin signaling pathway | 129 | 85 | 2.15E-03 |
| **KEGG:04371** | Apelin signaling pathway | 138 | 90 | 2.24E-03 |

### Supplemental Table 17 (continued).

| KEGG ID | KEGG Description | Universe | COSMIC and CLINVAR | Adjusted P-value |
|---|---|---|---|---|
| **KEGG:05235** | PD-L1 expression and PD-1 checkpoint pathway in cancer | 89 | 62 | 2.34E-03 |
| **KEGG:05231** | Choline metabolism in cancer | 98 | 67 | 2.83E-03 |
| **KEGG:04910** | Insulin signaling pathway | 137 | 89 | 3.06E-03 |
| **KEGG:04922** | Glucagon signaling pathway | 107 | 72 | 3.22E-03 |
| **KEGG:04144** | Endocytosis | 251 | 151 | 3.24E-03 |
| **KEGG:05226** | Gastric cancer | 148 | 95 | 3.33E-03 |
| **KEGG:04659** | Th17 cell differentiation | 105 | 70 | 6.43E-03 |
| **KEGG:05210** | Colorectal cancer | 86 | 59 | 7.15E-03 |
| **KEGG:04540** | Gap junction | 88 | 60 | 8.16E-03 |
| **KEGG:05135** | Yersinia infection | 136 | 87 | 8.37E-03 |
| **KEGG:04520** | Adherens junction | 71 | 50 | 9.16E-03 |
| **KEGG:04930** | Type II diabetes mellitus | 46 | 35 | 9.21E-03 |
| **KEGG:00562** | Inositol phosphate metabolism | 73 | 51 | 1.07E-02 |
| **KEGG:04931** | Insulin resistance | 108 | 71 | 1.10E-02 |
| **KEGG:04916** | Melanogenesis | 101 | 67 | 1.14E-02 |
| **KEGG:04024** | cAMP signaling pathway | 220 | 132 | 1.16E-02 |
| **KEGG:04917** | Prolactin signaling pathway | 70 | 49 | 1.35E-02 |
| **KEGG:05230** | Central carbon metabolism in cancer | 70 | 49 | 1.35E-02 |
| **KEGG:04960** | Aldosterone-regulated sodium reabsorption | 37 | 29 | 1.51E-02 |
| **KEGG:05223** | Non-small cell lung cancer | 72 | 50 | 1.56E-02 |
| **KEGG:05218** | Melanoma | 72 | 50 | 1.56E-02 |
| **KEGG:05163** | Human cytomegalovirus infection | 223 | 133 | 1.57E-02 |
| **KEGG:04664** | Fc epsilon RI signaling pathway | 67 | 47 | 1.68E-02 |
| **KEGG:04625** | C-type lectin receptor signaling pathway | 104 | 68 | 1.90E-02 |
| **KEGG:04721** | Synaptic vesicle cycle | 78 | 53 | 2.32E-02 |
| **KEGG:04071** | Sphingolipid signaling pathway | 119 | 76 | 2.41E-02 |
| **KEGG:04152** | AMPK signaling pathway | 121 | 77 | 2.59E-02 |
| **KEGG:04726** | Serotonergic synapse | 112 | 72 | 2.62E-02 |
| **KEGG:05219** | Bladder cancer | 41 | 31 | 2.74E-02 |
| **KEGG:04270** | Vascular smooth muscle contraction | 134 | 84 | 2.83E-02 |
| **KEGG:04933** | AGE-RAGE signaling pathway in diabetic complications | 100 | 65 | 3.26E-02 |
| **KEGG:05221** | Acute myeloid leukemia | 67 | 46 | 4.10E-02 |

**Supplemental Table 18.** Significant KEGG enrichments for COSMIC G4 mutations.

| GO ID | GO Description | Universe | COSMIC | Adjusted P-value |
|---|---|---|---|---|
| **KEGG:04611** | Platelet activation | 124 | 81 | 0.002339 |
| **KEGG:05226** | Gastric cancer | 148 | 94 | 0.002757 |
| **KEGG:04144** | Endocytosis | 251 | 149 | 0.002928 |
| **KEGG:05166** | Human T-cell leukemia virus 1 infection | 219 | 132 | 0.003096 |
| **KEGG:04926** | Relaxin signaling pathway | 129 | 83 | 0.00414 |
| **KEGG:04024** | cAMP signaling pathway | 220 | 132 | 0.004174 |
| **KEGG:04935** | Growth hormone synthesis, secretion and action | 120 | 78 | 0.004212 |
| **KEGG:04520** | Adherens junction | 71 | 50 | 0.005182 |
| **KEGG:04930** | Type II diabetes mellitus | 46 | 35 | 0.005861 |
| **KEGG:00562** | Inositol phosphate metabolism | 73 | 51 | 0.006052 |
| **KEGG:04666** | Fc gamma R-mediated phagocytosis | 96 | 64 | 0.006737 |
| **KEGG:04658** | Th1 and Th2 cell differentiation | 89 | 60 | 0.007009 |
| **KEGG:05231** | Choline metabolism in cancer | 98 | 65 | 0.007447 |
| **KEGG:04960** | Aldosterone-regulated sodium reabsorption | 37 | 29 | 0.010324 |
| **KEGG:04910** | Insulin signaling pathway | 137 | 86 | 0.010713 |
| **KEGG:04916** | Melanogenesis | 101 | 66 | 0.012641 |
| **KEGG:04721** | Synaptic vesicle cycle | 78 | 53 | 0.013364 |
| **KEGG:05135** | Yersinia infection | 136 | 85 | 0.014406 |
| **KEGG:05235** | PD-L1 expression and PD-1 checkpoint pathway in cancer | 89 | 59 | 0.015883 |
| **KEGG:04922** | Glucagon signaling pathway | 107 | 69 | 0.015977 |
| **KEGG:05230** | Central carbon metabolism in cancer | 70 | 48 | 0.019532 |
| **KEGG:05210** | Colorectal cancer | 86 | 57 | 0.020522 |
| **KEGG:05218** | Melanoma | 72 | 49 | 0.022161 |
| **KEGG:04931** | Insulin resistance | 108 | 69 | 0.023655 |
| **KEGG:04152** | AMPK signaling pathway | 121 | 76 | 0.025197 |
| **KEGG:04270** | Vascular smooth muscle contraction | 134 | 83 | 0.025742 |
| **KEGG:05163** | Human cytomegalovirus infection | 223 | 130 | 0.026284 |
| **KEGG:05220** | Chronic myeloid leukemia | 76 | 51 | 0.027645 |
| **KEGG:04659** | Th17 cell differentiation | 105 | 67 | 0.030128 |
| **KEGG:04625** | C-type lectin receptor signaling pathway | 104 | 66 | 0.041098 |
| **KEGG:05030** | Cocaine addiction | 49 | 35 | 0.043077 |
| **KEGG:04071** | Sphingolipid signaling pathway | 119 | 74 | 0.045066 |
| **KEGG:04917** | Prolactin signaling pathway | 70 | 47 | 0.04558 |
| **KEGG:04540** | Gap junction | 88 | 57 | 0.047383 |

**Supplemental Table 19.** Significant KEGG enrichments for CLINVAR G4 mutations.

| KEGG ID | KEGG Description | Universe | CLINVAR | Adjusted P-value |
|---|---|---|---|---|
| **KEGG:05414** | Dilated cardiomyopathy | 95 | 21 | 6.7E-07 |
| **KEGG:05410** | Hypertrophic cardiomyopathy | 90 | 20 | 1.4E-06 |
| **KEGG:05412** | Arrhythmogenic right ventricular cardiomyopathy | 77 | 18 | 3.4E-06 |
| **KEGG:04261** | Adrenergic signaling in cardiomyocytes | 150 | 23 | 1.0E-04 |
| **KEGG:04512** | ECM-receptor interaction | 88 | 16 | 5.1E-04 |
| **KEGG:04260** | Cardiac muscle contraction | 87 | 15 | 1.8E-03 |
| **KEGG:05221** | Acute myeloid leukemia | 67 | 12 | 7.9E-03 |
| **KEGG:05230** | Central carbon metabolism in cancer | 70 | 12 | 1.2E-02 |
| **KEGG:04919** | Thyroid hormone signaling pathway | 121 | 16 | 2.0E-02 |
| **KEGG:05220** | Chronic myeloid leukemia | 76 | 12 | 2.4E-02 |
| **KEGG:04912** | GnRH signaling pathway | 93 | 13 | 4.1E-02 |

**Supplemental Table 20.** Significant KEGG enrichments for COSMIC and CLINVAR G4 mutations leading to a G4 loss.

| KEGG ID | KEGG Description | Universe | COSMIC and CLINVAR | Adjusted P-value |
|---|---|---|---|---|
| **KEGG:05412** | Arrhythmogenic right ventricular cardiomyopathy | 77 | 44 | 4.37E-11 |
| **KEGG:05414** | Dilated cardiomyopathy | 95 | 50 | 7.37E-11 |
| **KEGG:05410** | Hypertrophic cardiomyopathy | 90 | 48 | 1.15E-10 |
| **KEGG:04261** | Adrenergic signaling in cardiomyocytes | 150 | 61 | 1.83E-07 |
| **KEGG:04921** | Oxytocin signaling pathway | 154 | 60 | 1.62E-06 |
| **KEGG:04010** | MAPK signaling pathway | 294 | 95 | 7.23E-06 |
| **KEGG:04015** | Rap1 signaling pathway | 210 | 73 | 1.11E-05 |
| **KEGG:04510** | Focal adhesion | 200 | 70 | 1.52E-05 |
| **KEGG:04020** | Calcium signaling pathway | 239 | 80 | 1.63E-05 |
| **KEGG:04725** | Cholinergic synapse | 113 | 44 | 1.41E-04 |
| **KEGG:04151** | PI3K-Akt signaling pathway | 353 | 104 | 2.35E-04 |
| **KEGG:04514** | Cell adhesion molecules | 153 | 53 | 6.96E-04 |
| **KEGG:05200** | Pathways in cancer | 529 | 142 | 9.19E-04 |
| **KEGG:04022** | cGMP-PKG signaling pathway | 166 | 56 | 9.35E-04 |
| **KEGG:04024** | cAMP signaling pathway | 220 | 69 | 1.41E-03 |
| **KEGG:04512** | ECM-receptor interaction | 88 | 34 | 2.64E-03 |
| **KEGG:04810** | Regulation of actin cytoskeleton | 216 | 67 | 2.74E-03 |
| **KEGG:04925** | Aldosterone synthesis and secretion | 98 | 36 | 5.22E-03 |
| **KEGG:04360** | Axon guidance | 181 | 57 | 6.73E-03 |
| **KEGG:04929** | GnRH secretion | 64 | 26 | 8.78E-03 |
| **KEGG:04713** | Circadian entrainment | 97 | 35 | 9.76E-03 |
| **KEGG:04911** | Insulin secretion | 86 | 32 | 1.00E-02 |
| **KEGG:04662** | B cell receptor signaling pathway | 79 | 30 | 1.07E-02 |
| **KEGG:04934** | Cushing syndrome | 153 | 49 | 1.34E-02 |
| **KEGG:04724** | Glutamatergic synapse | 114 | 39 | 1.42E-02 |
| **KEGG:05165** | Human papillomavirus infection | 331 | 91 | 1.55E-02 |
| **KEGG:04014** | Ras signaling pathway | 234 | 68 | 1.95E-02 |
| **KEGG:04728** | Dopaminergic synapse | 132 | 43 | 2.21E-02 |
| **KEGG:05032** | Morphine addiction | 90 | 32 | 2.49E-02 |
| **KEGG:04390** | Hippo signaling pathway | 157 | 49 | 2.56E-02 |
| **KEGG:05224** | Breast cancer | 147 | 46 | 3.61E-02 |
| **KEGG:04730** | Long-term depression | 59 | 23 | 4.14E-02 |
| **KEGG:04727** | GABAergic synapse | 89 | 31 | 4.49E-02 |

**Supplemental Table 21.** Significant KEGG enrichments for COSMIC G4 mutations leading to a G4 loss.

| KEGG ID | KEGG Description | Universe | COSMIC | Adjusted P-value |
|---|---|---|---|---|
| **KEGG:05410** | Hypertrophic cardiomyopathy | 90 | 37 | 9.29E-08 |
| **KEGG:05414** | Dilated cardiomyopathy | 95 | 38 | 1.37E-07 |
| **KEGG:05412** | Arrhythmogenic right ventricular cardiomyopathy | 77 | 32 | 9.49E-07 |
| **KEGG:04261** | Adrenergic signaling in cardiomyocytes | 150 | 43 | 5.39E-04 |
| **KEGG:04020** | Calcium signaling pathway | 239 | 59 | 1.65E-03 |
| **KEGG:04921** | Oxytocin signaling pathway | 154 | 42 | 2.51E-03 |
| **KEGG:04360** | Axon guidance | 181 | 45 | 1.33E-02 |
| **KEGG:04022** | cGMP-PKG signaling pathway | 166 | 42 | 1.47E-02 |
| **KEGG:04015** | Rap1 signaling pathway | 210 | 50 | 1.76E-02 |
| **KEGG:04510** | Focal adhesion | 200 | 48 | 1.92E-02 |
| **KEGG:04024** | cAMP signaling pathway | 220 | 51 | 2.94E-02 |
| **KEGG:04151** | PI3K-Akt signaling pathway | 353 | 74 | 3.78E-02 |

**Supplemental Table 22.** Significant KEGG enrichments for CLINVAR G4 mutations leading to a G4 loss.

| KEGG ID | KEGG Description | Universe | CLINVAR | Adjusted P-value |
|---|---|---|---|---|
| **KEGG:05410** | Hypertrophic cardiomyopathy | 90 | 10 | 5.17E-04 |
| **KEGG:05414** | Dilated cardiomyopathy | 95 | 10 | 7.44E-04 |
| **KEGG:05412** | Arrhythmogenic right ventricular cardiomyopathy | 77 | 9 | 9.61E-04 |
| **KEGG:04261** | Adrenergic signaling in cardiomyocytes | 150 | 9 | 4.67E-02 |
| **KEGG:05221** | Acute myeloid leukemia | 67 | 6 | 5.00E-02 |

**Supplemental Table 23:** Significant GO:CC enrichments for COSMIC and CLINVAR G4 mutations leading to a G4 gain.

| KEGG ID | KEGG Description | Universe | COSMIC and CLINVAR | Adjusted P-value |
|---|---|---|---|---|
| **KEGG:04929** | GnRH secretion | 64 | 22 | 1.99E-05 |
| **KEGG:05200** | Pathways in cancer | 529 | 89 | 5.62E-05 |
| **KEGG:04724** | Glutamatergic synapse | 114 | 30 | 8.27E-05 |
| **KEGG:04725** | Cholinergic synapse | 113 | 28 | 6.58E-04 |
| **KEGG:04919** | Thyroid hormone signaling pathway | 121 | 29 | 8.96E-04 |
| **KEGG:04015** | Rap1 signaling pathway | 210 | 42 | 1.20E-03 |
| **KEGG:04261** | Adrenergic signaling in cardiomyocytes | 150 | 33 | 1.45E-03 |
| **KEGG:05202** | Transcriptional misregulation in cancer | 192 | 39 | 1.70E-03 |
| **KEGG:04713** | Circadian entrainment | 97 | 24 | 3.03E-03 |
| **KEGG:04930** | Type II diabetes mellitus | 46 | 15 | 3.49E-03 |
| **KEGG:05210** | Colorectal cancer | 86 | 22 | 3.76E-03 |
| **KEGG:04010** | MAPK signaling pathway | 294 | 52 | 3.76E-03 |
| **KEGG:04928** | Parathyroid hormone synthesis, secretion and action | 106 | 25 | 4.76E-03 |
| **KEGG:04072** | Phospholipase D signaling pathway | 147 | 31 | 5.89E-03 |
| **KEGG:04730** | Long-term depression | 59 | 17 | 5.93E-03 |
| **KEGG:04921** | Oxytocin signaling pathway | 154 | 32 | 6.02E-03 |
| **KEGG:05218** | Melanoma | 72 | 19 | 7.97E-03 |
| **KEGG:04728** | Dopaminergic synapse | 132 | 28 | 1.16E-02 |
| **KEGG:04934** | Cushing syndrome | 153 | 31 | 1.22E-02 |
| **KEGG:04512** | ECM-receptor interaction | 88 | 21 | 1.52E-02 |
| **KEGG:05213** | Endometrial cancer | 58 | 16 | 1.62E-02 |
| **KEGG:04961** | Endocrine and other factor-regulated calcium reabsorption | 53 | 15 | 1.89E-02 |
| **KEGG:04510** | Focal adhesion | 200 | 37 | 1.91E-02 |
| **KEGG:04974** | Protein digestion and absorption | 103 | 23 | 2.09E-02 |
| **KEGG:04810** | Regulation of actin cytoskeleton | 216 | 39 | 2.15E-02 |
| **KEGG:05030** | Cocaine addiction | 49 | 14 | 2.72E-02 |
| **KEGG:04720** | Long-term potentiation | 67 | 17 | 2.91E-02 |
| **KEGG:04911** | Insulin secretion | 86 | 20 | 3.02E-02 |
| **KEGG:04151** | PI3K-Akt signaling pathway | 353 | 56 | 3.34E-02 |
| **KEGG:05165** | Human papillomavirus infection | 331 | 53 | 3.80E-02 |
| **KEGG:04540** | Gap junction | 88 | 20 | 4.06E-02 |

**Supplemental Table 24.** Significant KEGG enrichments for COSMIC G4 mutations leading to a G4 gain.

| KEGG ID | KEGG Description | Universe | COSMIC | Adjusted P-value |
|---|---|---|---|---|
| **KEGG:05218** | Melanoma | 72 | 13 | 1.45E-02 |
| **KEGG:04072** | Phospholipase D signaling pathway | 147 | 20 | 1.59E-02 |
| **KEGG:05030** | Cocaine addiction | 49 | 10 | 2.90E-02 |

**Supplemental Table 25.** Significant INTERPRO enrichments for COSMIC and CLINVAR G4 mutations.

| INTERPRO ID | COSMIC and CLINVAR | UNIVERSE | FDR |
|---|---|---|---|
| **IPR011993:Pleckstrin homology-like domain** | 150 | 446 | 1.41E-10 |
| **IPR001849:Pleckstrin homology domain** | 95 | 277 | 1.4E-06 |
| **IPR011009:Protein kinase-like domain** | 158 | 547 | 4.88E-06 |
| **IPR000719:Protein kinase, catalytic domain** | 146 | 502 | 9.19E-06 |
| **IPR013098:Immunoglobulin I-set** | 52 | 140 | 0.000295 |
| **IPR008271:Serine/threonine-protein kinase, active site** | 96 | 316 | 0.000295 |
| **IPR017441:Protein kinase, ATP binding site** | 113 | 390 | 0.000331 |
| **IPR017970:Homeobox, conserved site** | 65 | 193 | 0.000418 |
| **IPR001781:Zinc finger, LIM-type** | 32 | 75 | 0.001657 |
| **IPR001452:Src homology-3 domain** | 72 | 230 | 0.001677 |
| **IPR002219:Protein kinase C-like, phorbol ester/diacylglycerol binding** | 29 | 67 | 0.003026 |
| **IPR003598:Immunoglobulin subtype 2** | 76 | 254 | 0.004198 |
| **IPR013164:Cadherin, N-terminal** | 28 | 65 | 0.004198 |
| **IPR008936:Rho GTPase activation protein** | 35 | 95 | 0.013732 |
| **IPR015425:Actin-binding FH2** | 11 | 15 | 0.013732 |
| **IPR000008:C2 calcium-dependent membrane targeting** | 48 | 148 | 0.018196 |
| **IPR020479:Homeodomain, metazoa** | 34 | 93 | 0.018196 |
| **IPR013088:Zinc finger, NHR/GATA-type** | 24 | 57 | 0.021614 |
| **IPR001025:Bromo adjacent homology (BAH) domain** | 9 | 11 | 0.025203 |
| **IPR000536:Nuclear hormone receptor, ligand-binding, core** | 21 | 48 | 0.031914 |
| **IPR009057:Homeodomain-like** | 95 | 360 | 0.04254 |
| **IPR001478:PDZ domain** | 50 | 163 | 0.04254 |
| **IPR001628:Zinc finger, nuclear hormone receptor-type** | 20 | 46 | 0.045727 |

**Supplemental Table 26.** Significant INTERPRO enrichments for COSMIC G4 mutations.

| INTERPRO ID | COSMIC | Universe | FDR |
|---|---|---|---|
| **IPR011993:Pleckstrin homology-like domain** | 147 | 446 | 1.30E-10 |
| **IPR001849:Pleckstrin homology domain** | 93 | 277 | 1.46E-06 |
| **IPR011009:Protein kinase-like domain** | 153 | 547 | 1.13E-05 |
| **IPR000719:Protein kinase, catalytic domain** | 141 | 502 | 2.55E-05 |
| **IPR013098:Immunoglobulin I-set** | 50 | 140 | 7.98E-04 |
| **IPR017441:Protein kinase, ATP binding site** | 109 | 390 | 7.98E-04 |
| **IPR017970:Homeobox, conserved site** | 63 | 193 | 7.98E-04 |
| **IPR008271:Serine/threonine-protein kinase, active site** | 91 | 316 | 1.39E-03 |
| **IPR002219:Protein kinase C-like, phorbol ester/diacylglycerol binding** | 29 | 67 | 1.98E-03 |
| **IPR001781:Zinc finger, LIM-type** | 31 | 75 | 2.42E-03 |
| **IPR013164:Cadherin, N-terminal** | 28 | 65 | 2.72E-03 |
| **IPR001452:Src homology-3 domain** | 69 | 230 | 3.73E-03 |
| **IPR008936:Rho GTPase activation protein** | 35 | 95 | 7.92E-03 |
| **IPR015425:Actin-binding FH2** | 11 | 15 | 1.09E-02 |
| **IPR020479:Homeodomain, metazoa** | 34 | 93 | 1.09E-02 |
| **IPR003598:Immunoglobulin subtype 2** | 72 | 254 | 1.39E-02 |
| **IPR000008:C2 calcium-dependent membrane targeting** | 47 | 148 | 1.68E-02 |
| **IPR001025:Bromo adjacent homology (BAH) domain** | 9 | 11 | 2.07E-02 |
| **IPR001478:PDZ domain** | 50 | 163 | 2.28E-02 |
| **IPR013088:Zinc finger, NHR/GATA-type** | 23 | 57 | 3.61E-02 |
| **IPR009057:Homeodomain-like** | 93 | 360 | 3.71E-02 |
| **IPR015919:Cadherin-like** | 39 | 121 | 4.08E-02 |
| **IPR002126:Cadherin** | 38 | 118 | 4.83E-02 |

**Supplemental Table 27.** Significant INTERPRO enrichments for CLINVAR G4 mutations.

| INTERPRO ID | CLINVAR | UNIVERSE | FDR |
|---|---|---|---|
| **IPR000595:Cyclic nucleotide-binding domain** | 7 | 36 | 0.009358 |
| **IPR018490:Cyclic nucleotide-binding-like** | 7 | 39 | 0.009358 |

**Supplemental Table 28:** Top 50 significant transcription factor enrichments for COSMIC and CLINVAR G4.

| Transcription Factor ID | Transcription Factor Description | Universe | COSMIC and CLINVAR | FDR |
|---|---|---|---|---|
| **TF:M09636_1** | Factor: MAZ; motif: GGGMGGGGSSGGGGGGGGGGGG; match class: 1 | 14379 | 7641 | 6.97E-262 |
| **TF:M09973_1** | Factor: CPBP; motif: GNNRGGGHGGGGNNGGGRN; match class: 1 | 6788 | 4243 | 1.01E-254 |
| **TF:M09826_1** | Factor: BTEB3; motif: CCNNSCCNSCCCCKCCCCC; match class: 1 | 7694 | 4675 | 1.41E-249 |
| **TF:M07289_1** | Factor: GKLF; motif: NNNRGGNGNGGSN; match class: 1 | 10800 | 6111 | 2.57E-248 |
| **TF:M07039_1** | Factor: ETF; motif: CCCCGCCCCYN; match class: 1 | 13890 | 7403 | 5.24E-241 |
| **TF:M09973** | Factor: CPBP; motif: GNNRGGGHGGGGNNGGGRN | 11087 | 6214 | 2.70E-238 |
| **TF:M09984** | Factor: MAZ; motif: GGGGGAGGGGGNGRGRRRGNRG | 9762 | 5614 | 4.46E-236 |
| **TF:M12351_1** | Factor: TIEG1; motif: NCCCNSNCCCCGCCCCC; match class: 1 | 8412 | 4966 | 9.71E-228 |
| **TF:M09723** | Factor: BTEB1; motif: GGGGGCGGGGCNGSGGGNGS | 10228 | 5801 | 4.64E-226 |
| **TF:M09826** | Factor: BTEB3; motif: CCNNSCCNSCCCCKCCCCC | 11731 | 6462 | 1.50E-224 |
| **TF:M09984_1** | Factor: MAZ; motif: GGGGGAGGGGGNGRGRRRGNRG; match class: 1 | 5696 | 3615 | 3.02E-220 |
| **TF:M10026** | Factor: PATZ; motif: GGGGNGGGGGMKGGRRNGGNRN | 8607 | 5037 | 2.42E-219 |
| **TF:M07040_1** | Factor: GKLF; motif: NNRRGRRNGNSNNN; match class: 1 | 8337 | 4909 | 3.67E-219 |
| **TF:M00986_1** | Factor: Churchill; motif: CGGGNN; match class: 1 | 10609 | 5947 | 4.14E-216 |
| **TF:M09723_1** | Factor: BTEB1; motif: GGGGGCGGGGCNGSGGGNGS; match class: 1 | 6131 | 3819 | 2.47E-213 |
| **TF:M12160_1** | Factor: KLF15; motif: RCCMCRCCCMCN; match class: 1 | 8212 | 4823 | 1.49E-208 |
| **TF:M10432_1** | Factor: MAZ; motif: GGGMGGGGS; match class: 1 | 4484 | 2948 | 2.74E-203 |
| **TF:M12351** | Factor: TIEG1; motif: NCCCNSNCCCCGCCCCC | 12580 | 6766 | 2.59E-200 |
| **TF:M10432** | Factor: MAZ; motif: GGGMGGGGS | 9496 | 5399 | 2.34E-199 |
| **TF:M00933** | Factor: Sp1; motif: CCCCGCCCCN | 9913 | 5589 | 4.40E-199 |
| **TF:M10529** | Factor: Sp1; motif: RGGGMGGRGSNGGGG | 7039 | 4230 | 1.45E-197 |
| **TF:M04953** | Factor: Sp1; motif: GGNDGGRGGCGGGG | 8852 | 5093 | 1.56E-196 |
| **TF:M02089_1** | Factor: E2F-3; motif: GGCGGGN; match class: 1 | 9606 | 5440 | 8.00E-196 |
| **TF:M10112** | Factor: Miz-1; motif: NNRGGWGGGGGAGGGGMRR | 8878 | 5103 | 9.30E-196 |
| **TF:M12160** | Factor: KLF15; motif: RCCMCRCCCMCN | 12959 | 6914 | 1.26E-195 |
| **TF:M09636** | Factor: MAZ; motif: GGGMGGGGSSGGGGGGGGGGGG | 16533 | 8315 | 1.88E-194 |
| **TF:M10026_1** | Factor: PATZ; motif: GGGGNGGGGGMKGGRRNGGNRN; match class: 1 | 5053 | 3219 | 9.11E-192 |
| **TF:M01104_1** | Factor: MOVO-B; motif: GNGGGGG; match class: 1 | 5798 | 3599 | 1.13E-191 |
| **TF:M00932_1** | Factor: Sp1; motif: NNGGGGCGGGGNN; match class: 1 | 6212 | 3805 | 5.40E-191 |
| **TF:M07395_1** | Factor: Sp1; motif: NGGGGCGGGGN; match class: 1 | 6529 | 3956 | 1.81E-188 |
| **TF:M00931** | Factor: Sp1; motif: GGGGCGGGGC | 10524 | 5832 | 8.08E-187 |
| **TF:M00933_1** | Factor: Sp1; motif: CCCCGCCCCN; match class: 1 | 5316 | 3340 | 7.30E-186 |
| **TF:M09834** | Factor: ZNF148; motif: NNNNNNCCNNCCCCTCCCCCACCCN | 7099 | 4227 | 3.40E-185 |
| **TF:M00932** | Factor: Sp1; motif: NNGGGGCGGGGNN | 10669 | 5892 | 5.01E-185 |
| **TF:M01303** | Factor: SP1; motif: GGGGYGGGGNS | 8089 | 4697 | 5.64E-183 |
| **TF:M03876_1** | Factor: Kaiso; motif: GCMGGGRGCRGS; match class: 1 | 9311 | 5267 | 3.63E-182 |
| **TF:M07436** | Factor: WT1; motif: NNGGGNGGGSGN | 6637 | 3990 | 4.64E-181 |
| **TF:M07226** | Factor: SP1; motif: NCCCCKCCCCC | 8460 | 4865 | 2.87E-180 |
| **TF:M07397** | Factor: ZBP89; motif: CCCCKCCCCCNN | 7289 | 4306 | 2.95E-180 |
| **TF:M07289** | Factor: GKLF; motif: NNNRGGNGNGGSN | 14918 | 7675 | 3.00E-180 |
| **TF:M00196** | Factor: Sp1; motif: NGGGGGCGGGGYN | 10479 | 5792 | 1.13E-179 |

**Supplemental Table 28 (continued).**

| Transcription Factor ID | Transcription Factor Description | Universe | COSMIC and CLINVAR | FDR |
|---|---|---|---|---|
| **TF:M10071** | Factor: Sp1; motif: NGGGGGCGGGGCCNGGGGGGGG | 8705 | 4978 | 1.41E-179 |
| **TF:M00931_1** | Factor: Sp1; motif: GGGGCGGGGC; match class: 1 | 6075 | 3707 | 2.15E-179 |
| **TF:M11529_1** | Factor: E2F-2; motif: GCGCGCGCNCS; match class: 1 | 14789 | 7621 | 5.14E-179 |
| **TF:M07395** | Factor: Sp1; motif: NGGGGCGGGGN | 10901 | 5975 | 1.19E-177 |
| **TF:M00196_1** | Factor: Sp1; motif: NGGGGGCGGGGYN; match class: 1 | 6084 | 3703 | 4.14E-176 |
| **TF:M09970** | Factor: KLF3; motif: NNNNNNGGGCGGGGCNNGN | 7907 | 4589 | 3.12E-175 |
| **TF:M07039** | Factor: ETF; motif: CCCCGCCCCYN | 16656 | 8320 | 3.08E-174 |
| **TF:M01104** | Factor: MOVO-B; motif: GNGGGGG | 10486 | 5777 | 3.21E-173 |
| **TF:M12703_1** | Factor: ZNF383; motif: SSNGGGMGGNGSNGGS; match class: 1 | 4453 | 2863 | 3.94E-173 |

**Supplemental Table 29**. Count and percentage of effect of SNV calculated by thermodynamic MFE and ED changes in the G quadruplex sequence.

| Change in Stability by SNV | Change in multiconfirm by SNV | Frequency | Percentage |
|---|---|---|---|
| **Further stabilized** | less diversity | 6,417 | 17.105 |
| **no change** | less diversity | 3,835 | 10.222 |
| **Destabilized** | less diversity | 5,383 | 14.349 |
| **Further stabilized** | no change | 34 | 0.091 |
| **no change** | no change | 5,378 | 14.335 |
| **Destabilized** | no change | 39 | 0.104 |
| **Further stabilized** | more diversity | 3,984 | 10.619 |
| **no change** | more diversity | 2,849 | 7.594 |
| **Destabilized** | more diversity | 9,597 | 25.581 |

**Supplemental Table 30**. Effect of transition mutation G→A in chr10:122,143,482 on potential binding for multiple transcription factors. All effects are strong.

| Motif Position | Gene Symbol | Transcription factor binding match | Reference P-value | Alternate P-value | Allele Difference | Allele Effect Size |
|---|---|---|---|---|---|---|
| -3\|6 | NHLH1 | tgtgtgggcAggtgggttg | 0.0018 | 2.86E-05 | 2.1672 | 0.1850 |
| -8\|2 | FOXO3 | atgtgtgggcAggtgggttgg | 0.0033 | 0.0001 | 2.3166 | 0.1431 |
| -3\|6 | TAL1 | tgtgtgggcAggtgggttg | 0.0045 | 0.0001 | 2.3166 | 0.1814 |
| -12\|7 | TP53 | gtccattccatgtgtgggcAggtgggttgggtggggtga | 0.0031 | 0.0001 | 2.3166 | 0.1149 |
| -4\|7 | HES5 | catgtgtgggcAggtgggttggg | 0.0040 | 0.0001 | 2.2697 | 0.1547 |
| -4\|7 | HES7 | catgtgtgggcAggtgggttggg | 0.0041 | 0.0002 | 2.2817 | 0.14880 |
| 1\|8 | USF2 | gtgtgggcAggtgggtt | 0.0026 | 0.0002 | 1.8992 | 0.1282 |
| -11\|3 | EGR3 | ttccatgtgtgggcGggtgggttgggtgg | 3.47E-06 | 0.0002 | -1.8709 | -0.1170 |
| -12\|2 | EGR3 | ttccatgtgtgggcGggtgggttgggtgg | 5.21E-06 | 0.0002 | -1.8638 | -0.1082 |
| -11\|2 | EGR1 | tccatgtgtgggcGggtgggttgggtg | 6.63E-06 | 0.0002 | -1.8447 | -0.1058 |
| -9\|1 | EGR2 | atgtgtgggcGggtgggttgg | 5.96E-06 | 0.0002 | -1.4785 | -0.1112 |
| -11\|2 | EGR1 | tccatgtgtgggcGggtgggttgggtg | 4.59E-06 | 0.0003 | -1.6940 | -0.1215 |
| -11\|3 | EGR2 | ttccatgtgtgggcGggtgggttgggtgg | 5.46E-06 | 0.0003 | -1.7277 | -0.1245 |
| -12\|3 | EGR1 | attccatgtgtgggcGggtgggttgggtggg | 1.35E-05 | 0.0004 | -1.9116 | -0.1130 |
| -8\|1 | EGR1 | tgtgtgggcGggtgggttg | 1.62E-05 | 0.0006 | -1.9401 | -0.1347 |
| -6\|3 | ZNF740 | tgtgtgggcGggtgggttg | 4.86E-05 | 0.0011 | -1.6642 | -0.1248 |
| -6\|3 | SP1 | tgtgtgggcGggtgggttg | 0.0001 | 0.0017 | -1.0496 | -0.1072 |
| -6\|4 | KLF16 | atgtgtgggcGggtgggttgg | 0.0001 | 0.0021894 | -1.5426264 | -0.12738754 |
| -7\|9 | SP4 | cattccatgtgtgggcGggtgggttgggtgggg | 0.0002 | 0.0023592 | -1.76102043 | -0.10166014 |
| -6\|4 | SP1 | atgtgtgggcGggtgggttgg | 0.0002 | 0.00291348 | -1.15354231 | -0.11339262 |
| -3\|6 | SP1 | tgtgtgggcGggtgggttg | 0.0002 | 0.00323868 | -1.21127823 | -0.11785306 |
| -6\|3 | ZNF740 | tgtgtgggcGggtgggttg | 0.0002 | 0.00376701 | -1.71699463 | -0.13807642 |
| -9\|2 | ZBTB7A | catgtgtgggcGggtgggttggg | 0.0004 | 0.0038684 | -1.3008656 | -0.11661897 |
| -11\|5 | SP4 | cattccatgtgtgggcGggtgggttgggtgggg | 0.0002 | 0.0047031 | -1.86702905 | -0.14590419 |
| -6\|4 | SP3 | atgtgtgggcGggtgggttgg | 0.0003 | 0.00499582 | -1.29081923 | -0.13319603 |

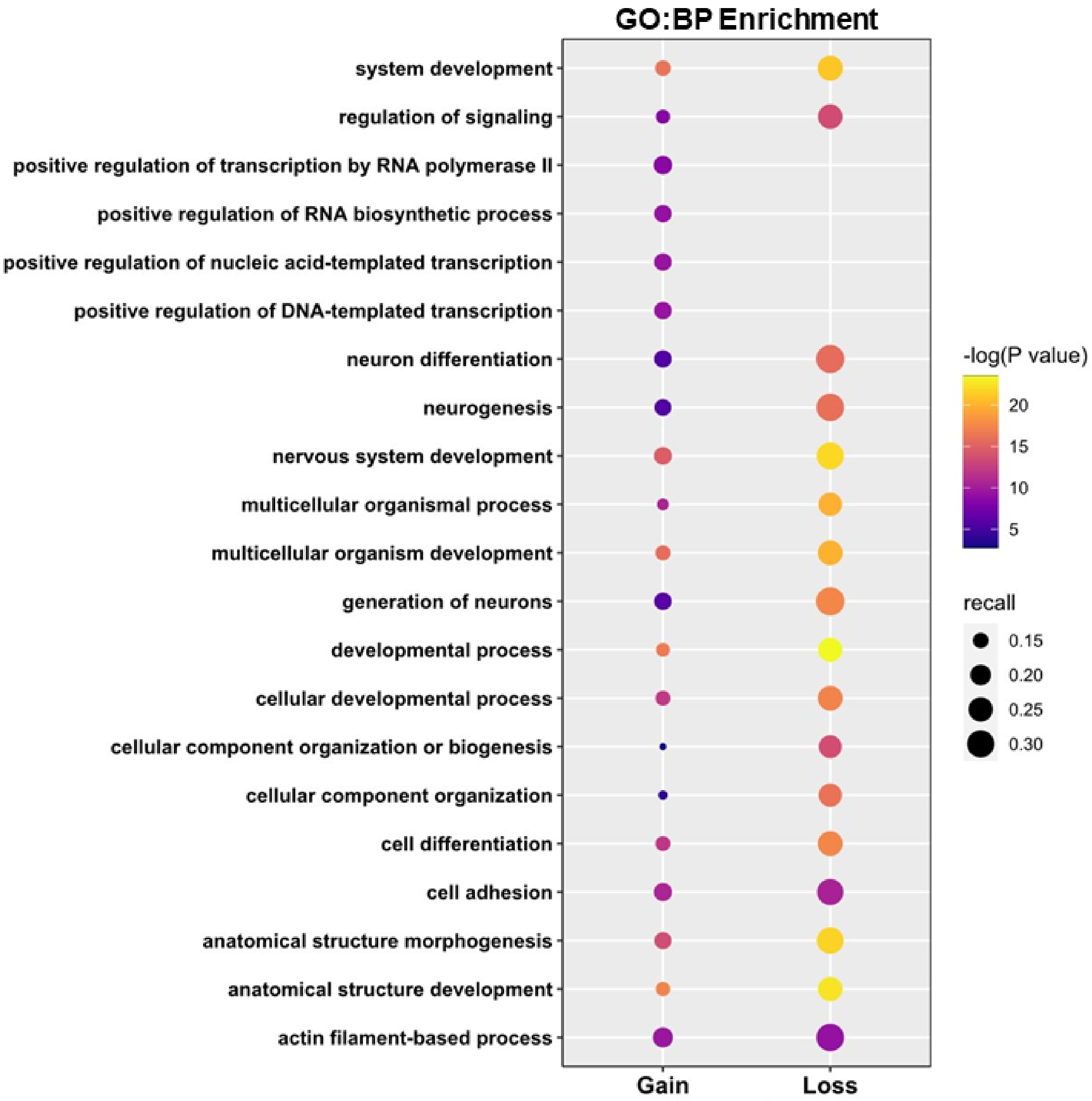


**Supplemental Figure 1.** Top 25 enriched GO:BP terms for COSMIC and CLINVAR G4 mutations.

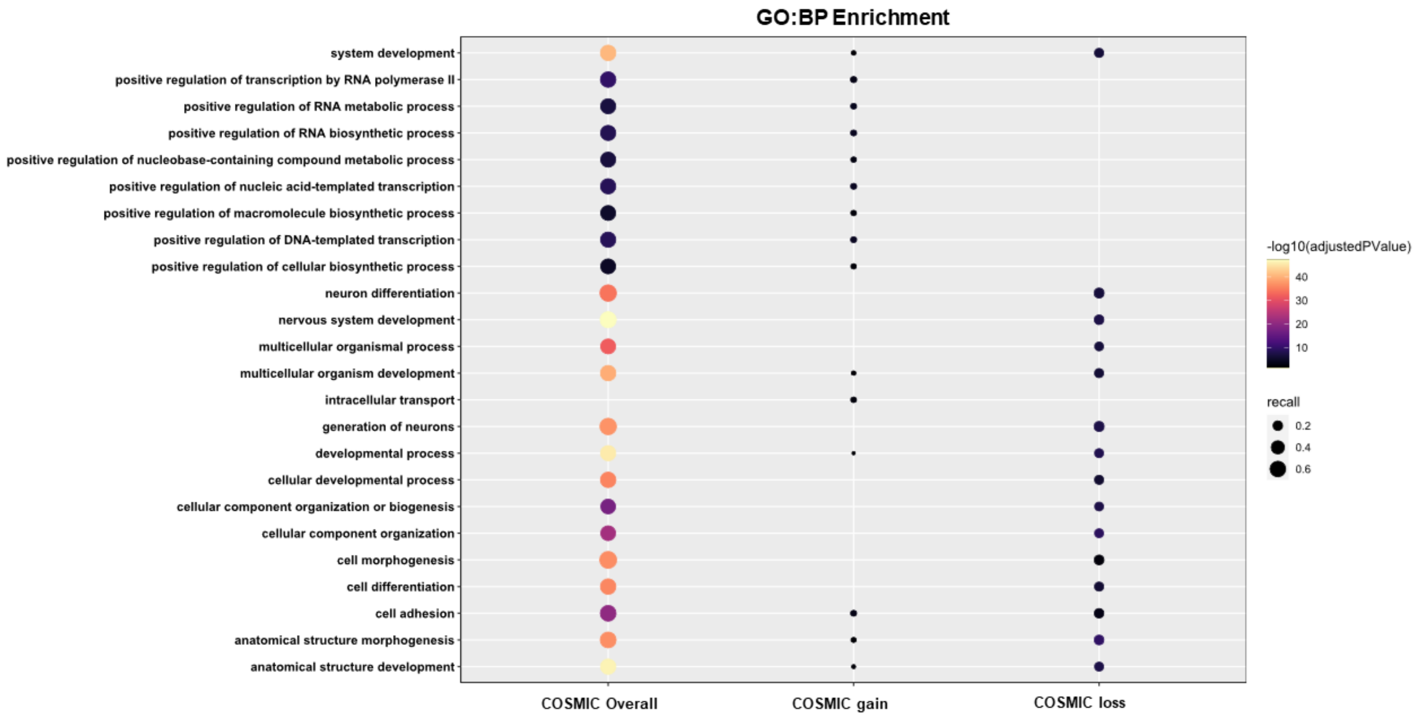


**Supplemental Figure 2.** Top 25 enriched GO:BP terms for COSMIC G4 mutations.

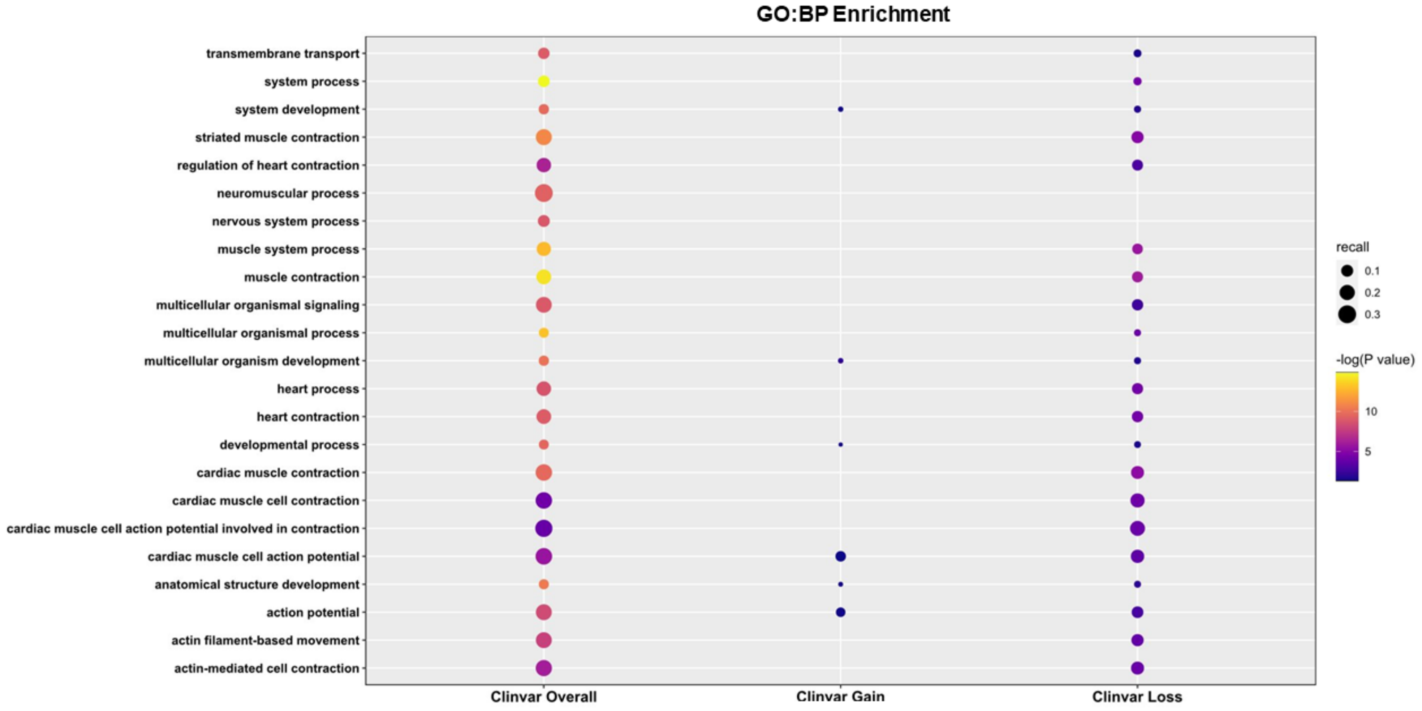


**Supplemental Figure 3.** Top 25 enriched GO:BP terms for CLINVAR G4 mutations.

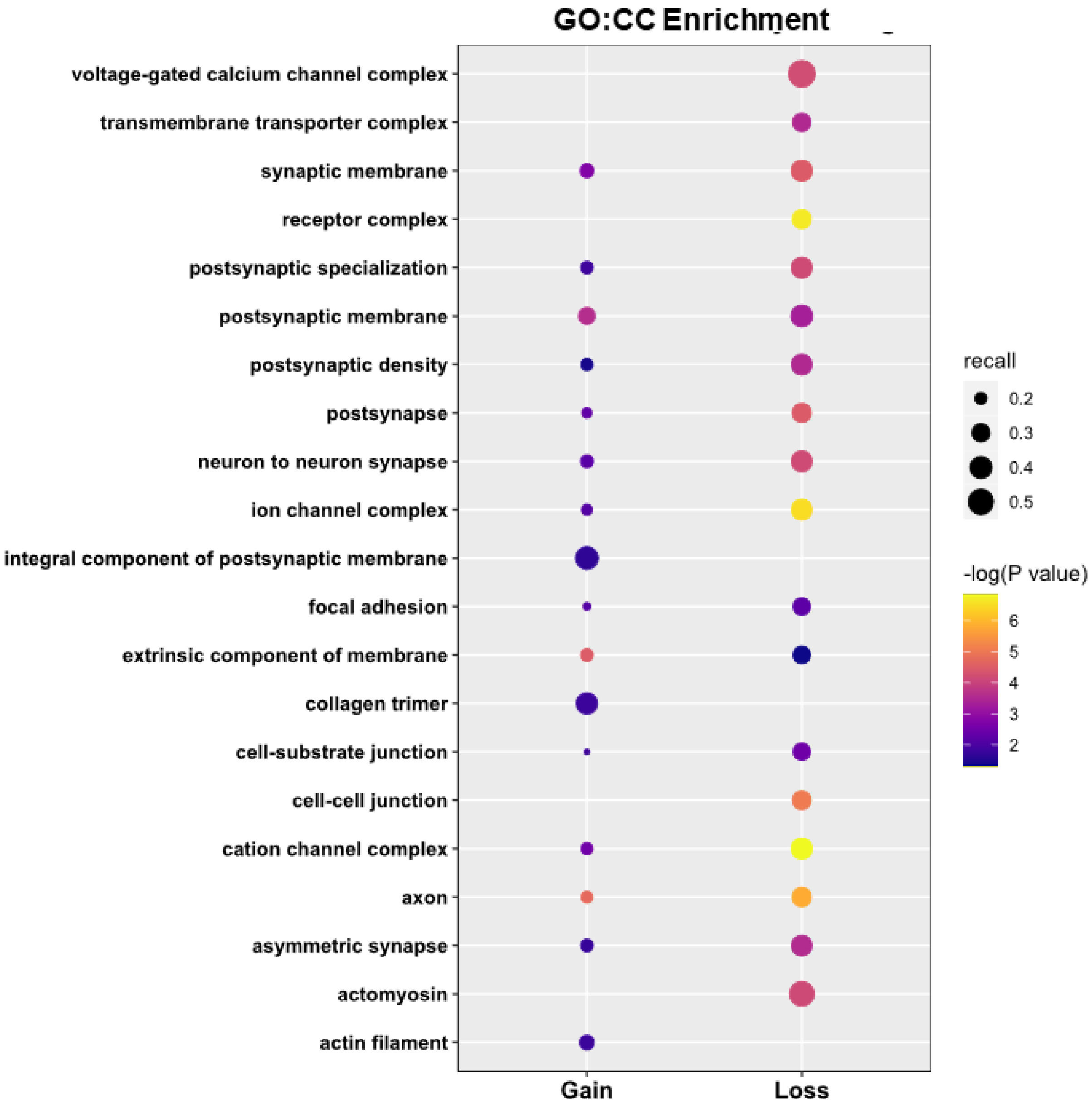


**Supplemental Figure 4.** Top 25 enriched GO:CC terms for COSMIC and CLINVAR G4 mutations.

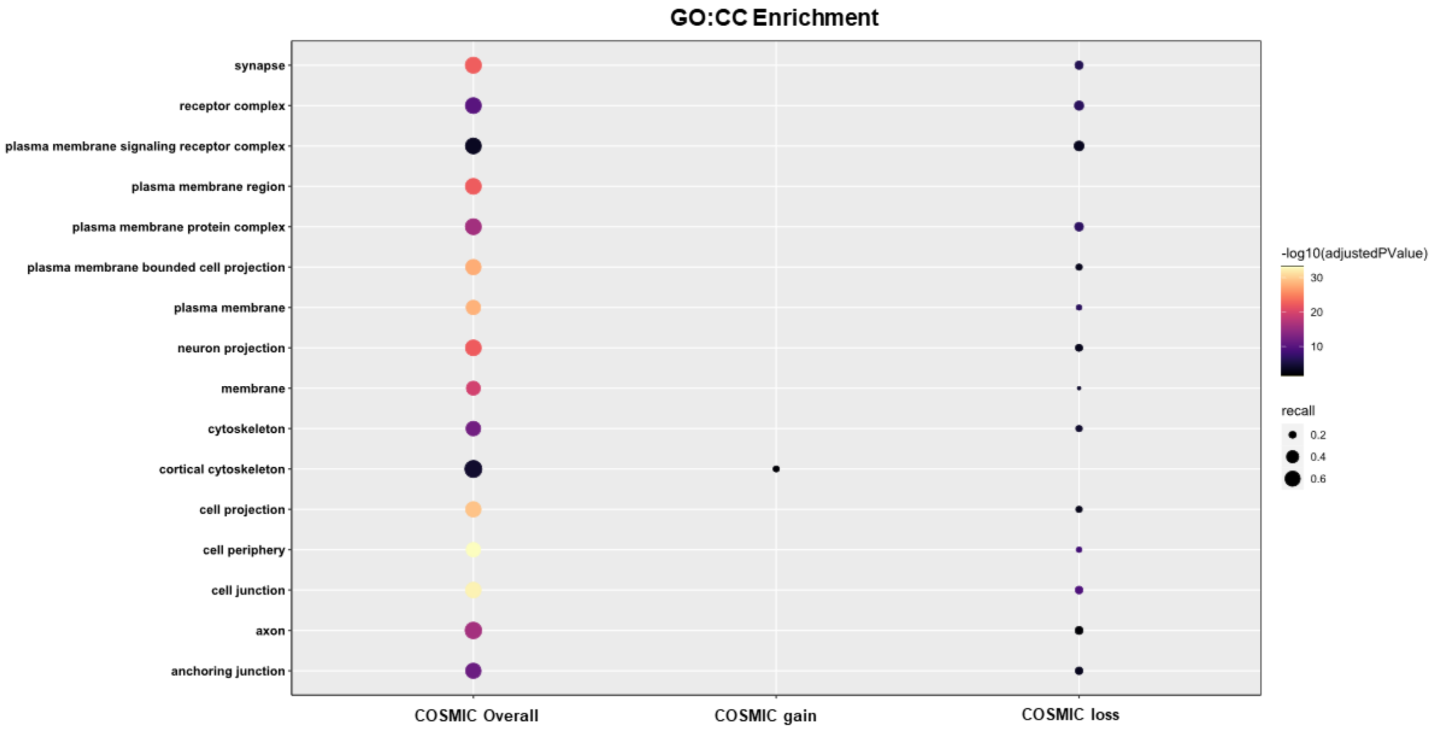


**Supplemental Figure 5.** Top 25 enriched GO:CC terms for COSMIC G4 mutations.

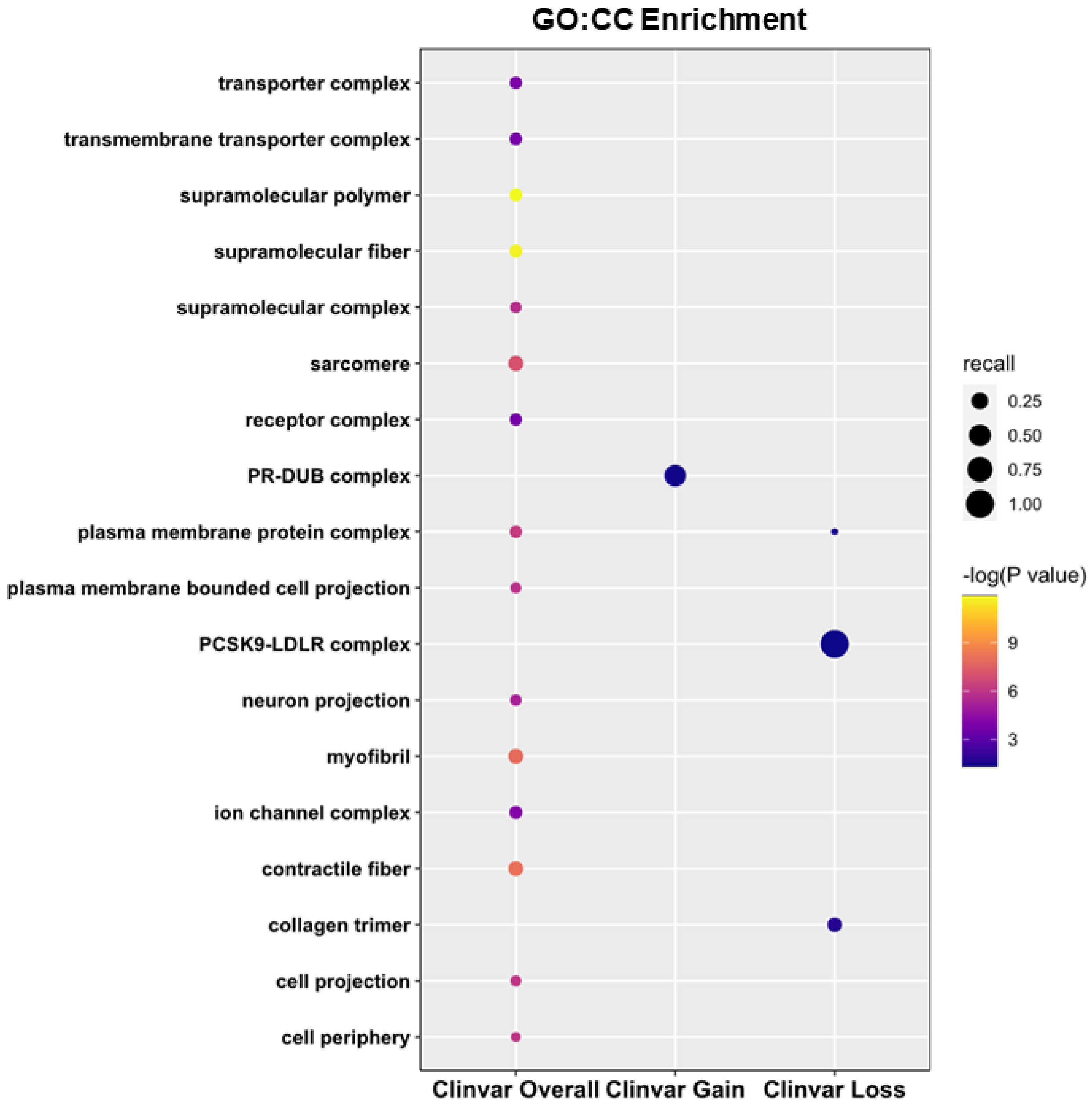


**Supplemental Figure 6.** Top 25 enriched GO:CC terms for CLINVAR G4 mutations.

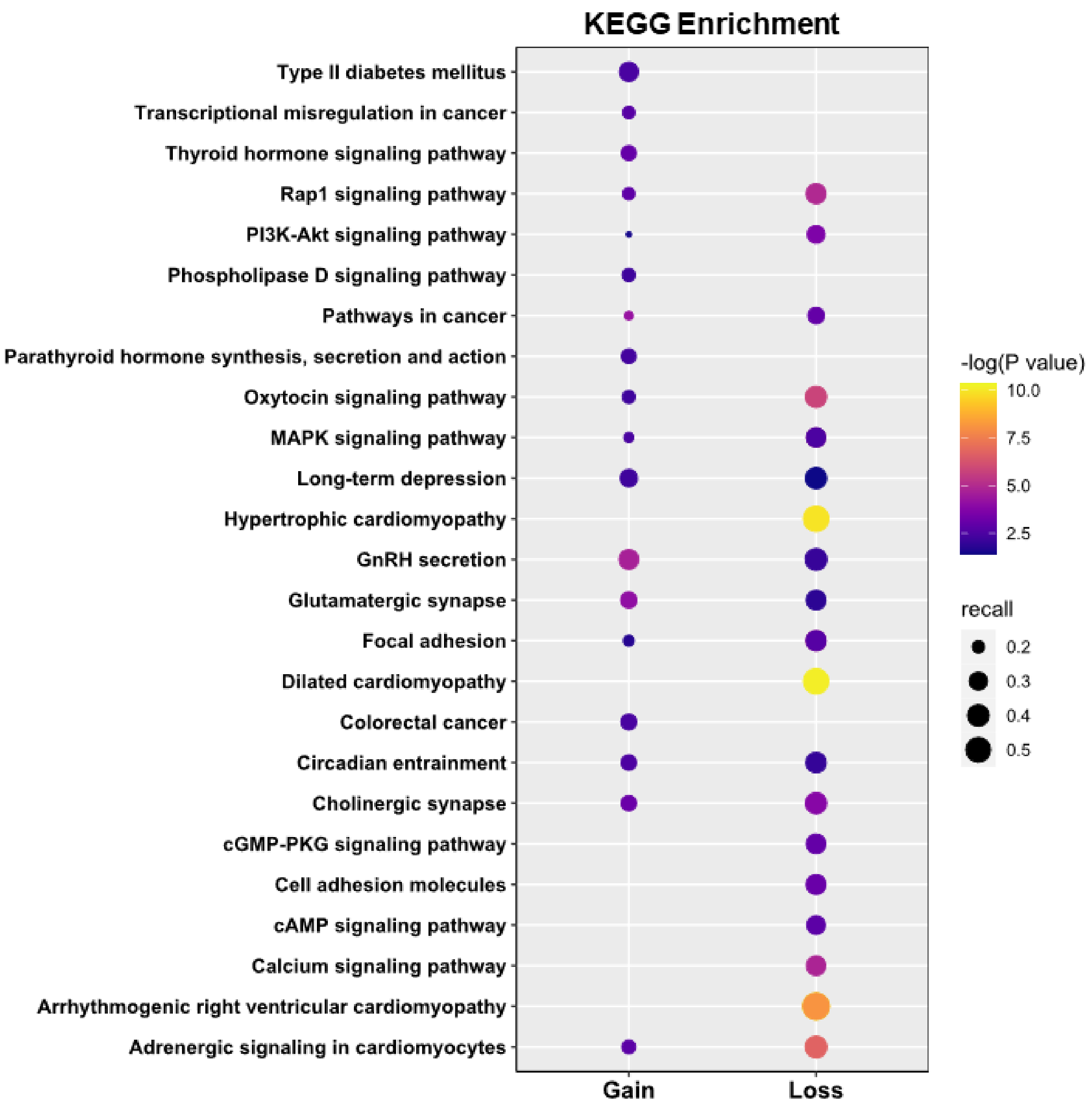


**Supplemental Figure 7.** Top 25 enriched KEGG terms for COSMIC and CLINVAR G4 mutations.

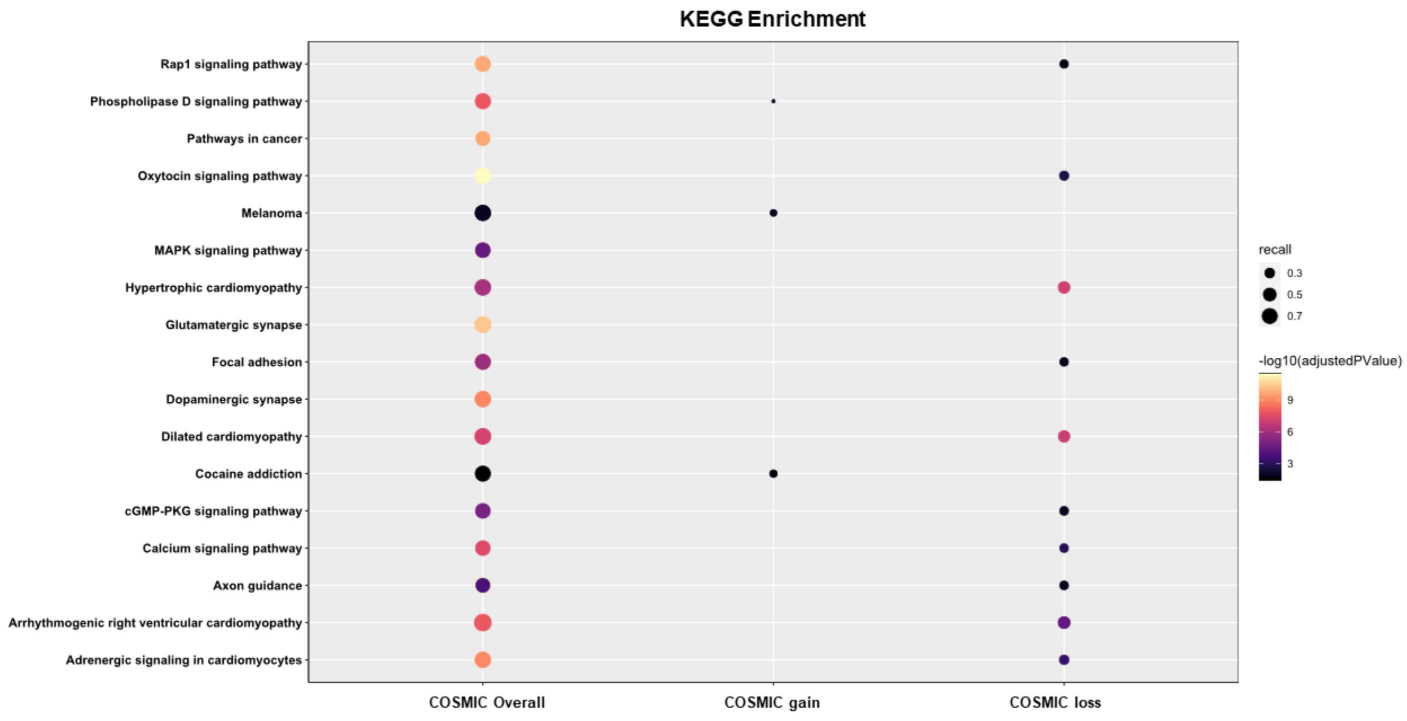


**Supplemental Figure 8.** Top 25 enriched KEGG terms for COSMIC G4 mutations.

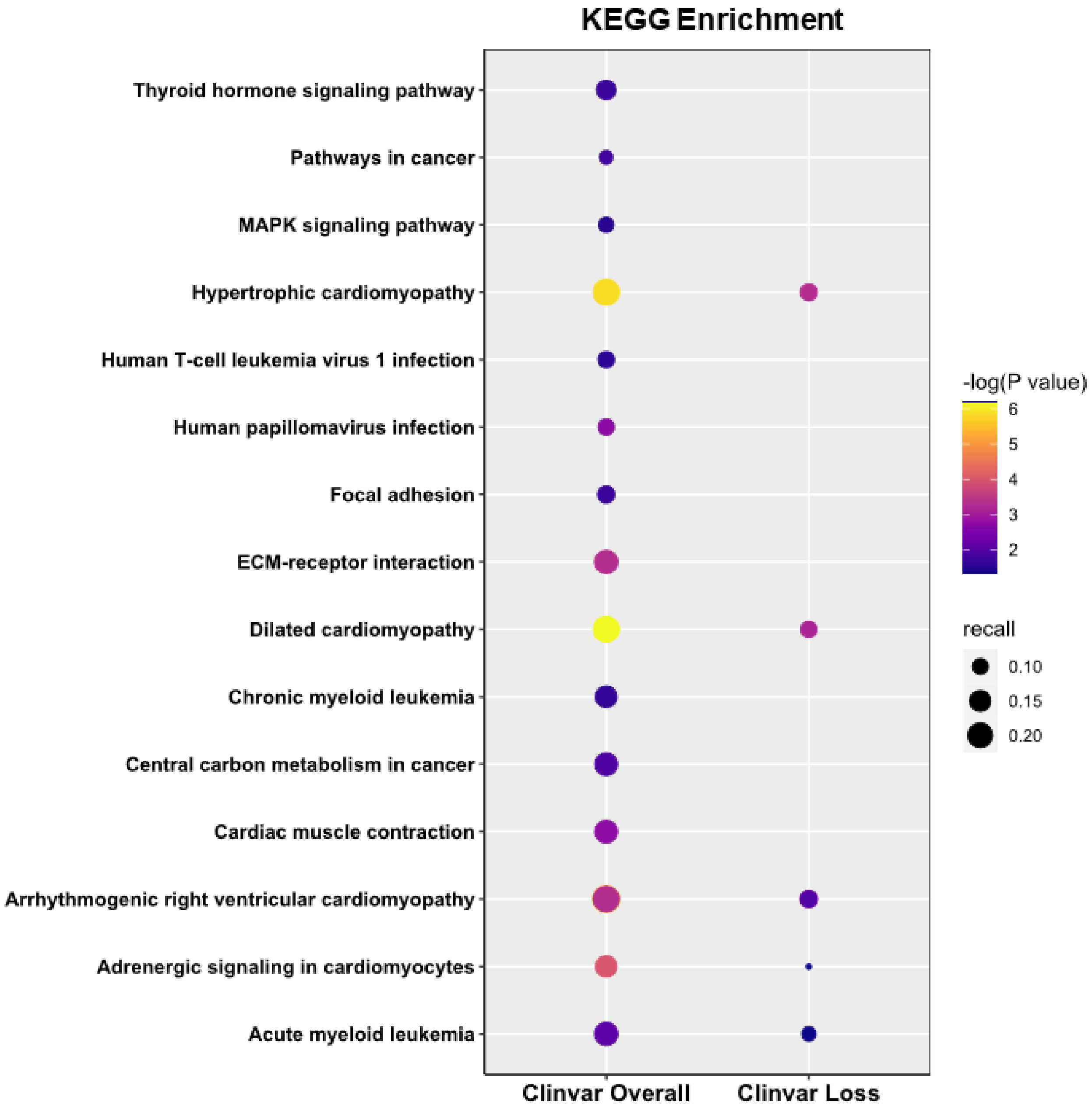


**Supplemental Figure 9.** Top 25 enriched KEGG terms for CLINVAR G4 mutations.

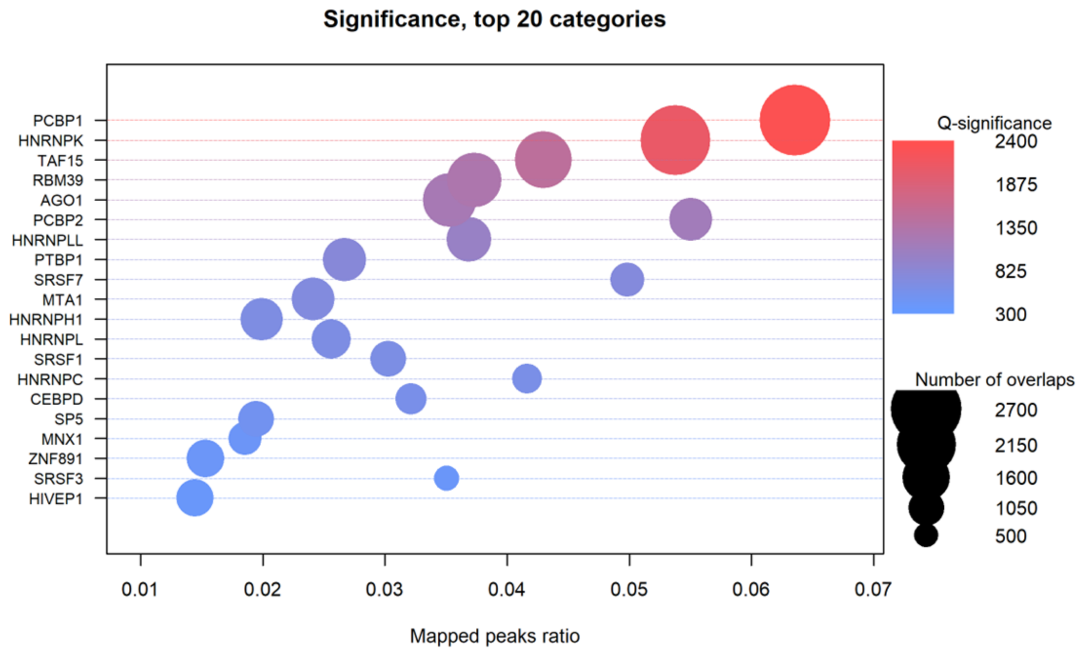


**Supplemental Figure 10.** Top 20 enriched transcription factors with overlapping ChIP-seq peaks for COSMIC and CLINVAR G4 SNVs in the HEK293 cell line.

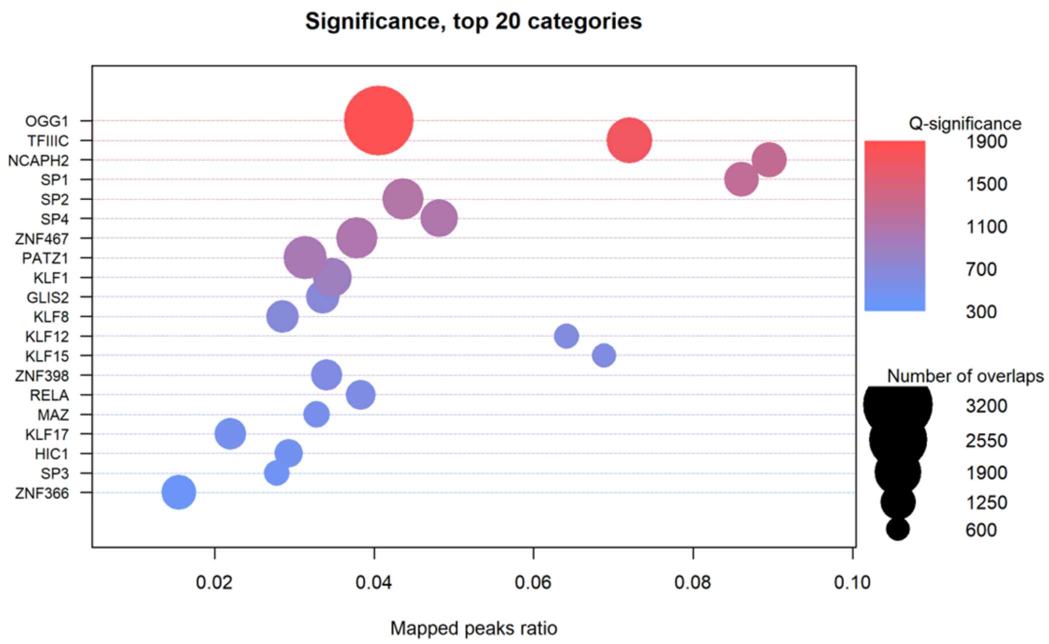


**Supplemental Figure 11.** Top 20 enriched transcription factors with overlapping ChIP-seq peaks for COSMIC and CLINVAR G4 SNVs in the K562 cell line.

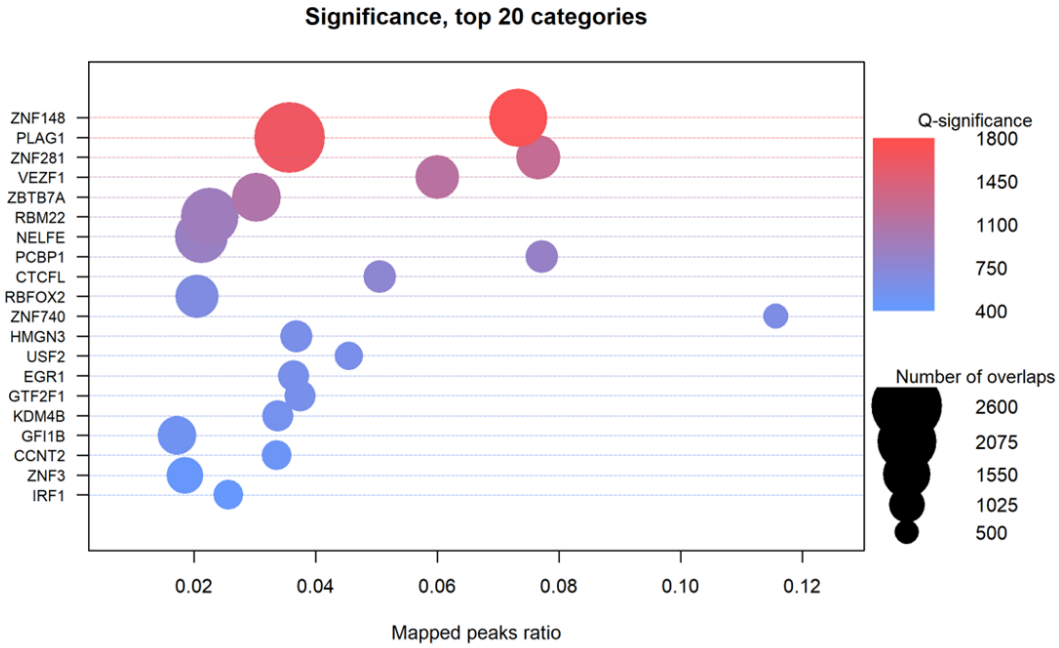


**Supplemental Figure 12.** Top 20 enriched transcription factors with overlapping ChIP-seq peaks for COSMIC and CLINVAR G4 SNVs in the Hep-G2 cell line.

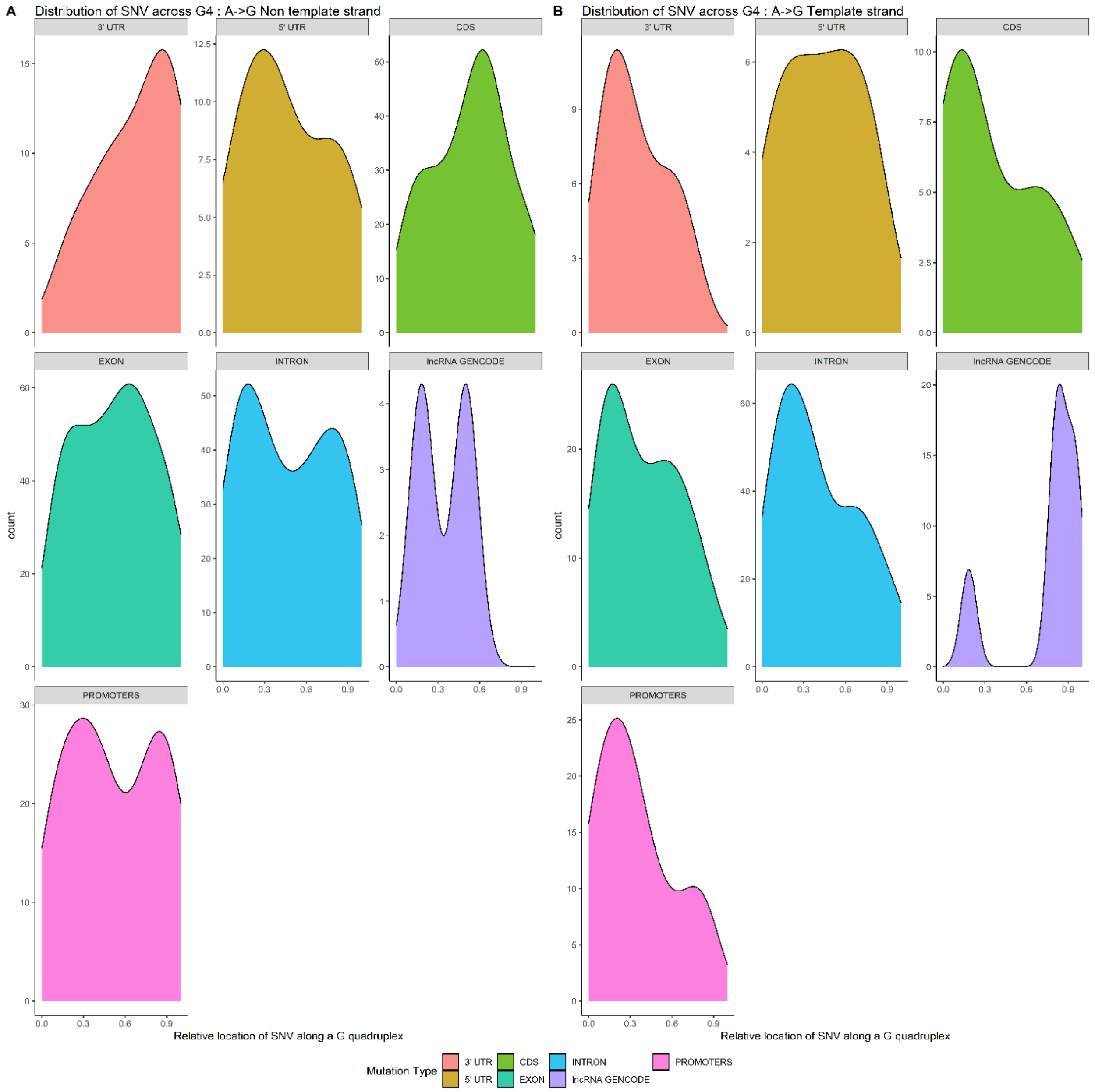


**Supplemental Figure 13**. Distribution of A→G SNVs across the G4 region for different features on (A) the non-template and (B) template strand.

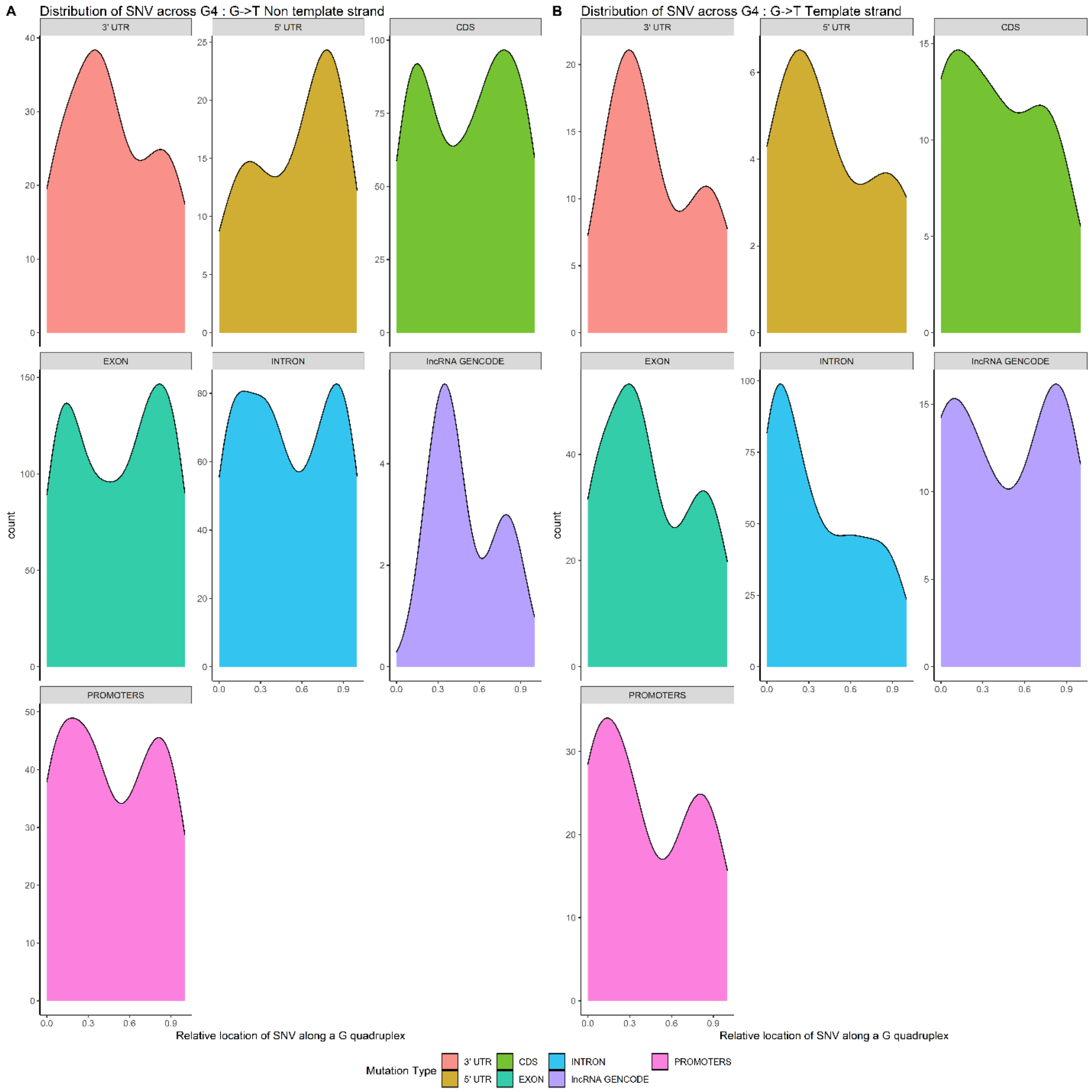


**Supplemental Figure 14**. Distribution of G→T SNVs across the G4 region for different features on (A) the non-template and (B) template strand.

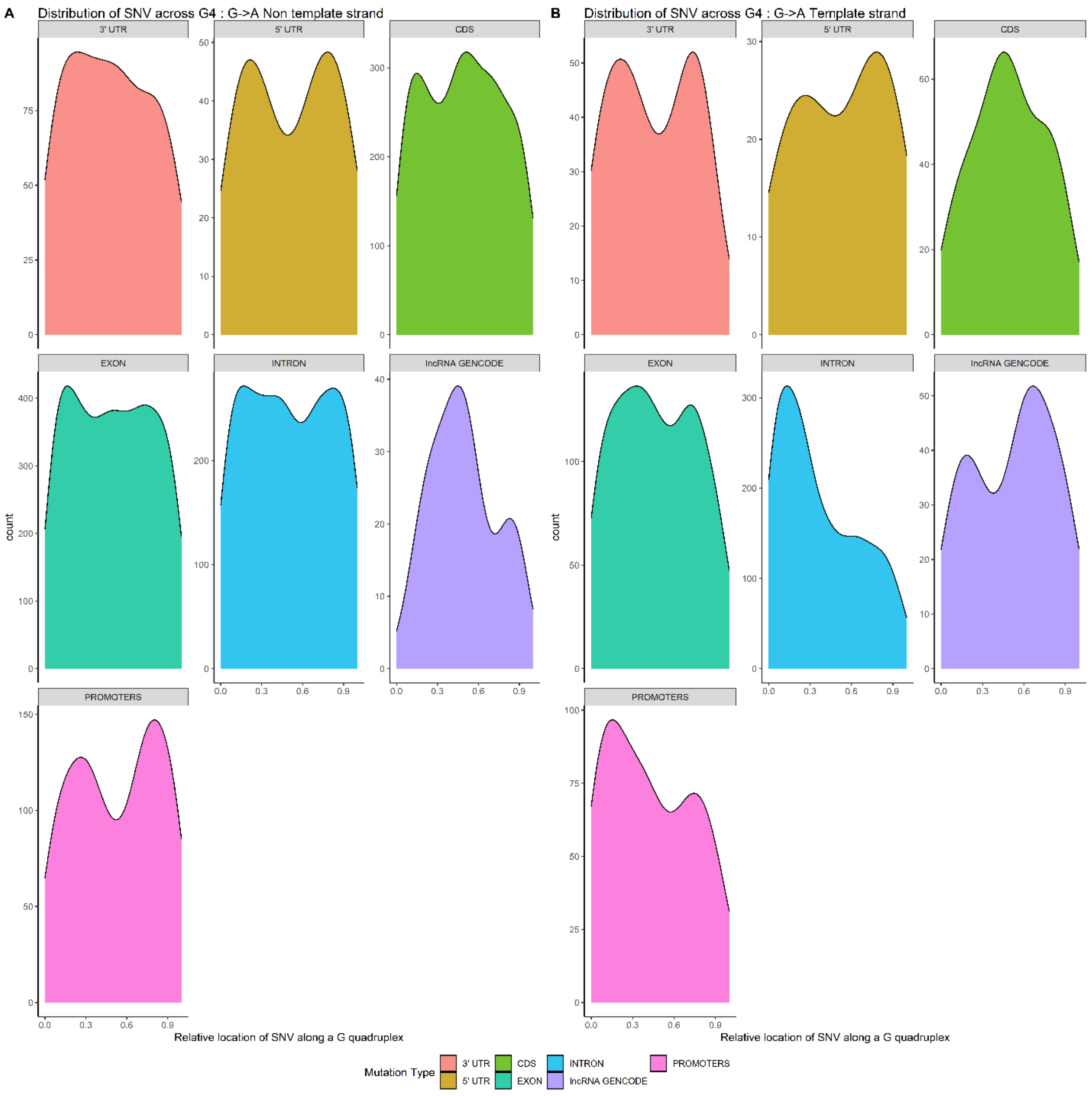


**Supplemental Figure 15**. Distribution of G→A SNVs across the G4 region for different features on (A) the non-template and (B) template strand.

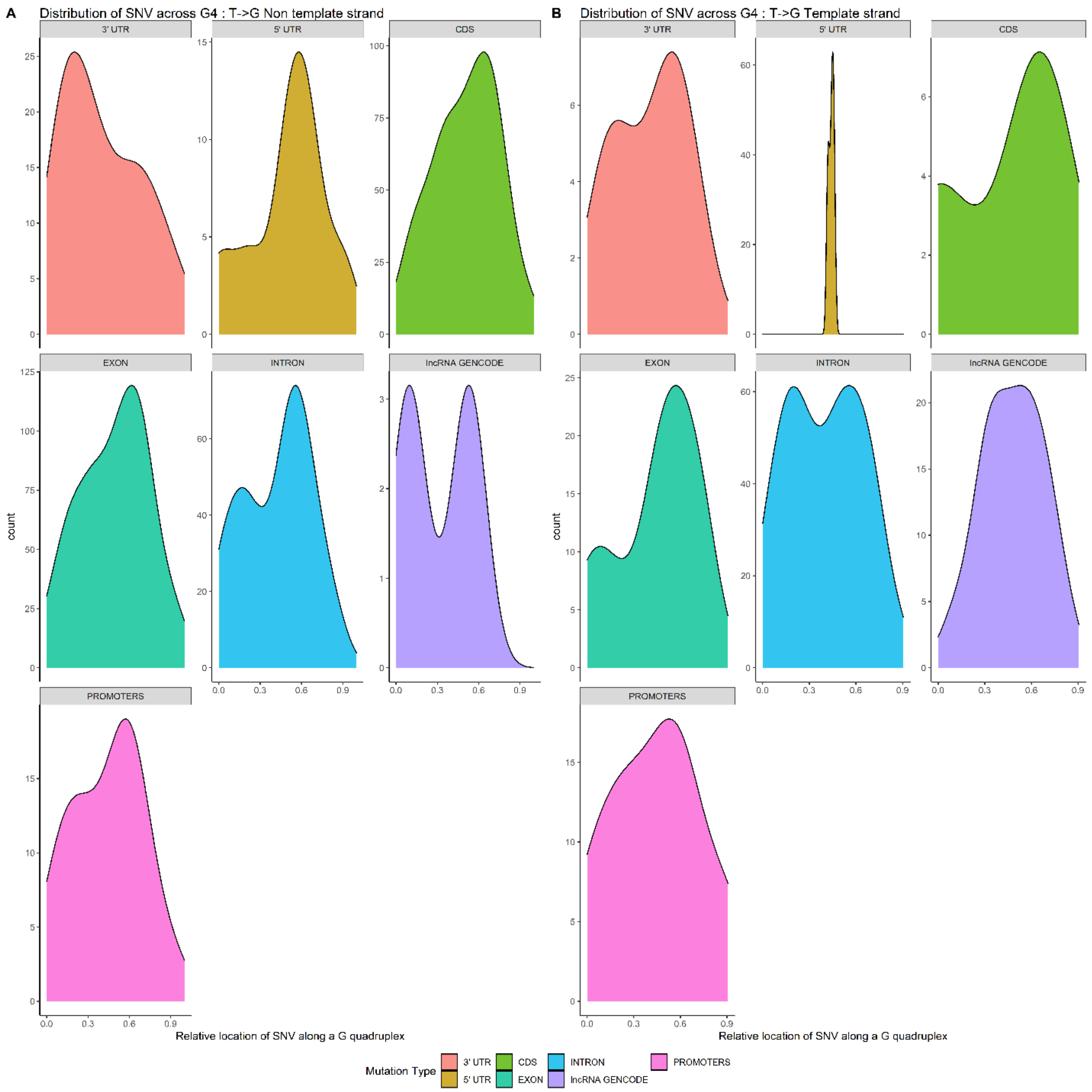


**Supplemental Figure 16**. Distribution of T→G SNVs across the G4 region for different features on (A) the non-template and (B) template strand.

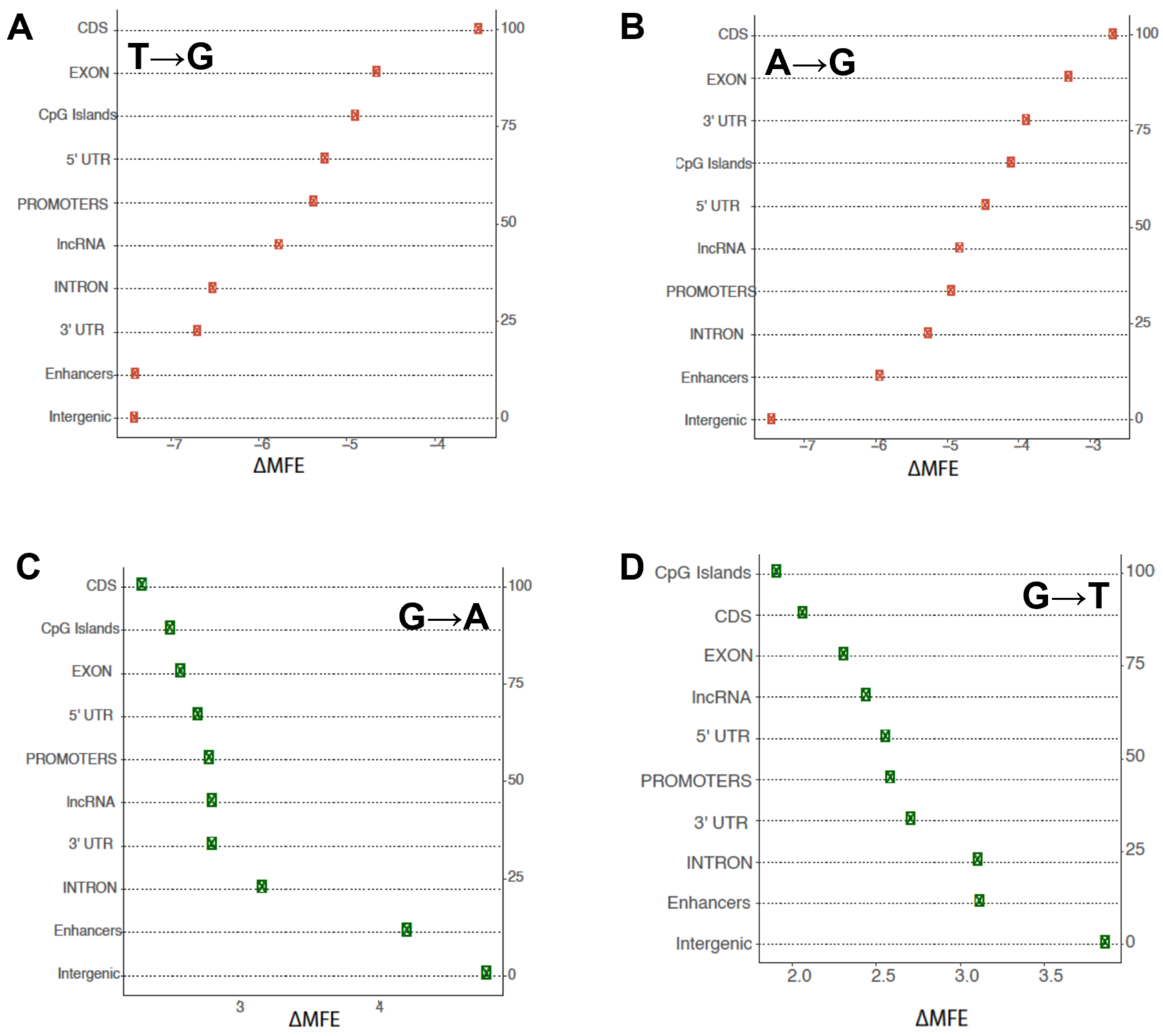


**Supplemental Figure 17.** Effect of each SNV on Δ MFE of G4 on different annotations with percentage of the counts shown in the secondary y axis. Shown is (A) T→G SNVs; (B) A→G SNVs; (C) G→A SNVs; and (D) G→T SNVs.

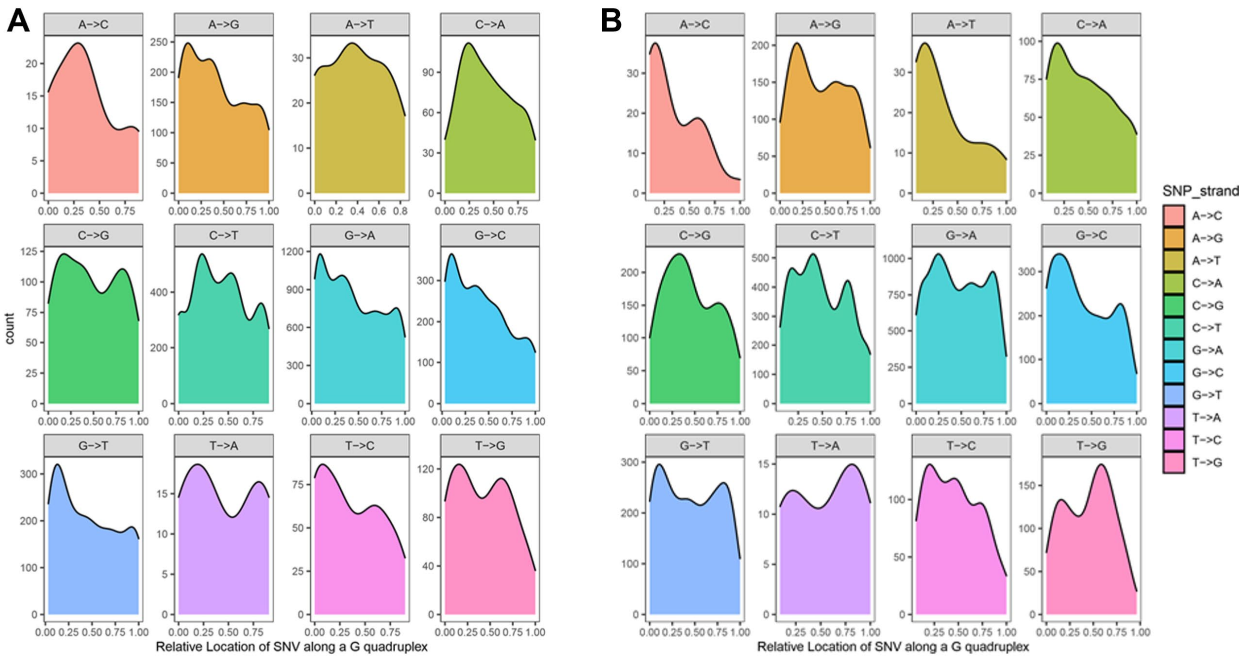


**Supplemental Figure 18**. Distribution of SNVs across G-quadruplex regions for the (A) forward and (B) reverse strands for SNVs detected in the CLINVAR database.
